# Mutation-agnostic gene insertion therapy for RHO-associated autosomal dominant retinitis pigmentosa using zinc finger nucleases

**DOI:** 10.64898/2026.07.13.738123

**Authors:** Akishi Onishi, Tetsushi Sakuma, Michiko Mandai, Takashi Watanabe, Takaho Endo, Wataru Nomura, Aiko Ishimaru, Ken-ichi Inoue, Junki Sho, Yoko Ohigashi, Yuki Nakano, Kazushi Yasuda, Atsuta Ozaki, Akiko Maeda, Chikako Morinaga, Tetsuo Itoh, Yuji Inomata, Yukihide Momozawa, Takashi Yamamoto, Hiroshi Kiyonari, Seiji Hori, Masayo Takahashi

**Author notes:** Correspondence: Akishi Onishi, VCGT Inc., Kobe, Japan. These authors contributed equally to this work (Akishi Onishi, Tetsushi Sakuma).

## Abstract

**Purpose:** Autosomal dominant retinitis pigmentosa caused by mutations in the rhodopsin gene (*RHO*-adRP) is among the most prevalent inherited retinal dystrophies. With nearly 100 distinct pathogenic variants identified to date, the mutational heterogeneity of *RHO*-adRP severely limits the clinical utility of mutation-specific therapeutic strategies. We aimed to develop a mutation-agnostic gene insertion therapy using homology-independent targeted integration (HITI) mediated by zinc finger nuclease ZF-ND1 targeting the human *RHO* 5′-UTR, and to validate its preclinical efficacy, safety, proof-of-concept, and proof-of-mechanism.

**Methods:** We developed two AAV serotype 5 (AAV5) vectors: one encoding the ZF-ND1 pair (AAV5-ZFN) and one carrying the therapeutic donor cassette (AAV5-RHO), delivered by subretinal co-injection. ZF pairs targeting the *RHO* 5′-UTR were arranged and refined by *in vitro* validation; AAV vector optimization and mechanistic verification were performed in human induced pluripotent stem cell (hiPSC)-derived retinal organoids-derived retinal organoids; longitudinal proof-of-concept efficacy and safety were assessed in a humanized *RHO*-T17M rat disease model by 6-month optical coherence tomography (OCT); and proof-of-mechanism was evaluated in non-human primate retina.

**Results:** We identified a ZF-ND1 pair achieving cleavage efficiency comparable to the SpCas9 RNP previously validated for HITI-mediated editing in mouse retina, and optimized the ZF array composition and NLS configuration for efficient editing especially in post-mitotic photoreceptors. HITI-mediated donor integration was confirmed across multiple cell types and ZFN:donor ratios. In the humanized rat disease model, the therapeutic vector provided outer nuclear layer (ONL) preservation by 6 months, with AAV5-ZFN:AAV5-RHO ratios of 1:1 and 1:2 maintaining ONL thickness above the preservation threshold. In the non-human primate retina, the fraction of HITI-edited rod photoreceptors exceeded the 20% therapeutic correction threshold in the successfully treated individual.

**Conclusions:** These findings support the advancement of this therapeutic vector to first-in-human trials as a mutation-agnostic insertion therapy applicable to all patients with *RHO*-adRP, irrespective of the specific causative variant.

## Introduction

Retinitis pigmentosa (RP) is the most prevalent inherited retinal dystrophy, affecting approximately 1 in 3,000–4,000 individuals worldwide and representing a leading cause of progressive blindness in the working-age population[1,2]. More than 250 causative genes have been identified[3], among which mutations in the rhodopsin gene (*RHO*) constitute the most common cause of autosomal dominant RP (adRP), accounting for up to 25% of dominant cases in Western cohorts[4,5], with lower prevalence reported in Japanese cohorts[6–8]. *RHO*-adRP is driven by two primary pathogenic mechanisms: haploinsufficiency, in which a single functional *RHO* allele cannot sustain photoreceptor viability[9]; and dominant toxicity arising from mutant rhodopsin through dominant-negative interference, gain-of-function, or both[5,10]. Thus, neither simple gene supplementation nor silencing of the mutant allele alone can fully neutralize the toxic effect of dominant pathogenic variants, and a fundamentally different therapeutic approach is therefore required.

Two therapeutic strategies are under clinical or advanced preclinical development for dominant *RHO* mutations. The first employs allele-specific suppression targeting prevalent variants such as P23H mutation (RHO1-2, QR-1123) [11,12]; however, this approach is restricted to patients carrying the targeted variant and fails to address the broad mutational heterogeneity of *RHO*-adRP, nor does it provide *RHO* supplementation to prevent haploinsufficiency-driven degeneration. The second strategy combines mutation-independent knockdown/knockout with exogenous *RHO* supplementation (EDIT-103, OPGx-RHO)[13,14]; while variant-agnostic in principle, the level of supplemented rhodopsin expression is proportional to AAV transduction efficiency, introducing a risk of overexpression toxicity at high vector doses[15] or haploinsufficiency under suboptimal transduction. Base editing, offering single-nucleotide precision without permanent double-strand breaks, is inherently variant-specific[16]. With nearly 100 pathogenic *RHO* variants reported to date, and with additional rare and *de novo* variants expected to emerge as genetic screening becomes more widely available[4,17,18], no single base editor can serve as a universal therapy[10,19]. A therapeutic design capable of treating all known and future *RHO* variants with a single product and protocol, while restoring *RHO* expression at physiologically appropriate levels under endogenous promoter control, therefore represents the most clinically rational approach.

Homology-independent targeted integration (HITI) is a genome editing strategy that uses non-homologous end joining (NHEJ) genome repairing machinery to achieve stable, site-specific donor insertion in post-mitotic cells, which is not achievable with HDR-dependent approaches that are infrequent in terminally differentiated tissues[20,21]. The underlying design principle of directional NHEJ-mediated integration through inverted target sequences flanking the donor was earlier demonstrated with heterodimeric ZFNs and plasmid donors in dividing cultured cells by the Obligate Ligation-Gated Recombination (ObLiGaRe) method[22]; HITI extended this principle to non-dividing cells in vivo through AAV-mediated delivery. We have previously demonstrated proof-of-mechanism and proof-of-concept for HITI-mediated therapy in a mouse model of *Rho* disease and explored its applicability to other autosomal dominant RP subtypes[23]. Here, we employ HITI at the human *RHO* 5′-UTR such that the therapeutic donor cassette is inserted under the transcriptional control of the endogenous *RHO* promoter, simultaneously restoring *RHO* expression and substantially suppressing transcription of the mutant allele via 5′-UTR disruption. Because the target site lies within the 5′-UTR rather than the mutation-specific coding sequence, this design is inherently variant-agnostic[24,25]. Translating this strategy to human clinical application, however, requires a genome editing tool that can be packaged within AAV without compromise to expression efficiency, a requirement that SpCas9 (∼4.2 kb) and even compact alternatives such as SaCas9 (∼3.2 kb) and CjCas9 (∼3.0 kb) fail to meet when combined with promoter, gRNA cassette, and ITRs[26].

In this study, we use ZF-ND1[27,28], whose combined coding length (∼2.6 kb) enables single-AAV packaging, whose obligate heterodimer architecture structurally suppresses non-specific cleavage[29], and whose 6-finger ZF array provides the extended target specificity required for precise editing at the *RHO* 5′-UTR[30–32]. We report the stepwise preclinical development and validation encompassing: (1) ZF module arrangement and *in vitro* refinement and validation of ZF-ND1; (2) vector optimization and mechanism verification in human induced pluripotent stem cell (hiPSC)-derived retinal organoids; (3) longitudinal proof-of-concept efficacy and safety validation in a rat disease model; and (4) proof-of-mechanism in non-human primate retina. These data support the advancement of this therapeutic vector to first-in-human trials as a mutation-agnostic gene insertion therapy for *RHO*-adRP.

## Results

### Development and Optimization of ZF-ND1 for HITI-Mediated Editing at the Human *RHO* 5**′**-UTR

Achieving high-efficiency, site-specific HITI editing in post-mitotic rod photoreceptors requires a ZF-ND1 pair with both stringent target specificity and robust cleavage activity in non-dividing cells. ZF-ND1 consists of a pair of 6-finger ZFN subunits, each recognizing an 18-bp half-site and connected to the heterodimeric FirmCut nuclease domain (ND1-DDD and ND1-RRR) via a TGGS linker. To identify and optimize such a pair, we implemented a four-step strategy (Fig. 1A): (1) arrangement of candidate ZF array pairs targeting a 95-bp window of the human *RHO* 5’-UTR using ZF Tools[30] (http://www.zincfingertools.org); (2) *in vitro* cleavage validation by single-strand annealing (SSA) assay; (3) ZF refinement by SSA, Cel-1, and Tracking of Indels by Decomposition (TIDE) assays (steps 1–3 addressed in this section); and (4) regulatory sequence optimization in hiPSC-derived retinal organoids (Fig. 2; addressed in the following section). NLS optimization in post-mitotic mouse rod photoreceptors was also evaluated in parallel (Supplementary Fig. S2).

**Figure 1.**
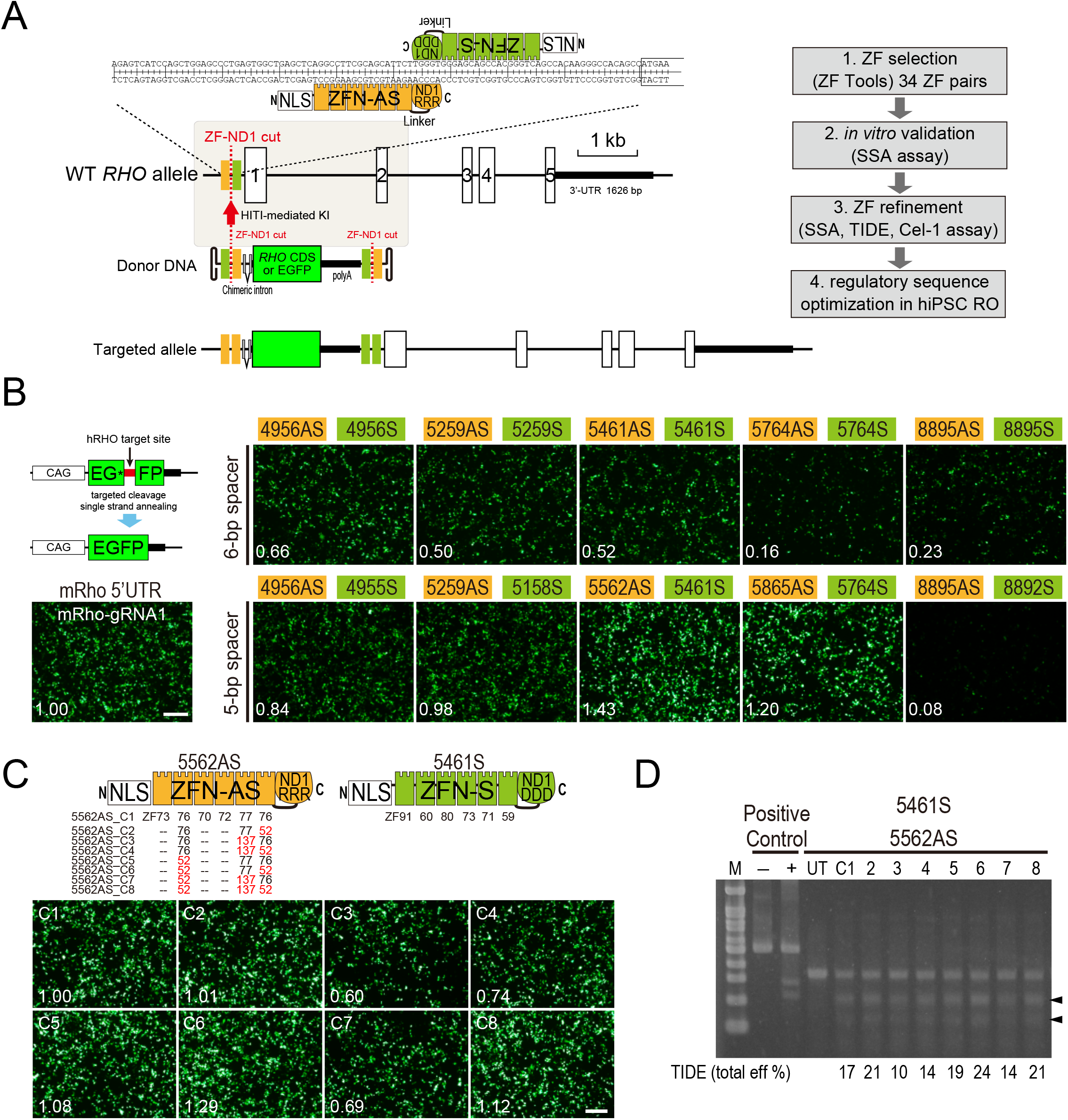
Development and Optimization of ZF-ND1 for HITI-Mediated Editing at the Human RHO 5′-UTR. **(A)** Schematic illustration of ZF-ND1-mediated HITI strategy and four-step development workflow. Left: structure of the human *RHO* allele (exons 1–5) and ZF-ND1 target site within the 5′-UTR. Each ZF-ND1 subunit comprises an N-terminal 3×multiNLS, a 6-finger ZF array, a TGGS linker, and the ND1-DDD or ND1-RRR heterodimeric nuclease domain. Obligate heterodimerization of ND1-DDD and ND1-RRR is required for nuclease activity. The donor DNA cassette, flanked by ZF-ND1 recognition sequences in reverse orientation, contains a chimeric intron, RHO CDS (or EGFP reporter), and polyA signal. HITI-mediated integration at the 5′-UTR places the donor under endogenous *RHO* promoter control. Right: four-step workflow for ZF-ND1 development: (1) ZF array design using ZF Tools (34 candidate pairs); (2) *in vitro* validation by SSA assay; (3) ZF refinement by SSA, Cel-1, and TIDE; (4) regulatory sequence optimization in hiPSC-derived retinal organoids (Fig. 2). **(B)** SSA assay screening of 34 candidate ZF pairs targeting the human RHO 5′-UTR. Representative fluorescence microscopy images of HEK293T cells 48 hours after co-transfection with pCAG-EGxxFP, ZF-ND1 expression plasmid pairs, and pCAG-mCherry (mCherry channel not shown for clarity). EGFP reconstitution (green) indicates targeted cleavage. SpCas9/mRho-gRNA1 included as a positive control used in our previous study. Normalized EGFP:mCherry ratios (relative to SpCas9/mRho-gRNA1 = 1.00) are shown in the lower-left corner of each panel. Panels grouped by spacer length: 6-bp spacer pairs (upper row) and 5-bp spacer pairs (lower row). Scale bar = 500 µm. **(C)** ZF module compositions of 5461S and 5562AS variants C1–C8. Upper: protein domain architecture of 5562AS (left) and 5461S (right) subunits, with ZF module numbers indicated for each array position; compatible ZF module substitutions at each position of 5562AS generate eight variants (C1–C8). Lower: SSA fluorescence images of HEK293T cells co-transfected with 5461S and 5562AS variants C1–C8. Normalized EGFP:mCherry ratios (relative to C1 = 1.00) are shown in the lower-left corner of each panel. Scale bar = 500 µm. **(D)** Cel-1 (Surveyor) nuclease assay and TIDE analysis of 5461S + 5562AS-C1 through C8. Top: Cel-1 gel. Lanes: M, DNA size marker; −, kit positive control without Detection Enzyme; +, kit positive control with Detection Enzyme; UT, untransfected cell lysate; C1–C8, HEK293T cells transfected with 5461S paired with 5562AS-C1 through C8, treated with Detection Enzyme. Arrowheads indicate cleavage bands. Band intensity assessed qualitatively. Bottom: TIDE indel efficiencies for C1–C8 shown beneath the corresponding gel lanes (see also Supplementary Fig. S1).

**Figure 2.**
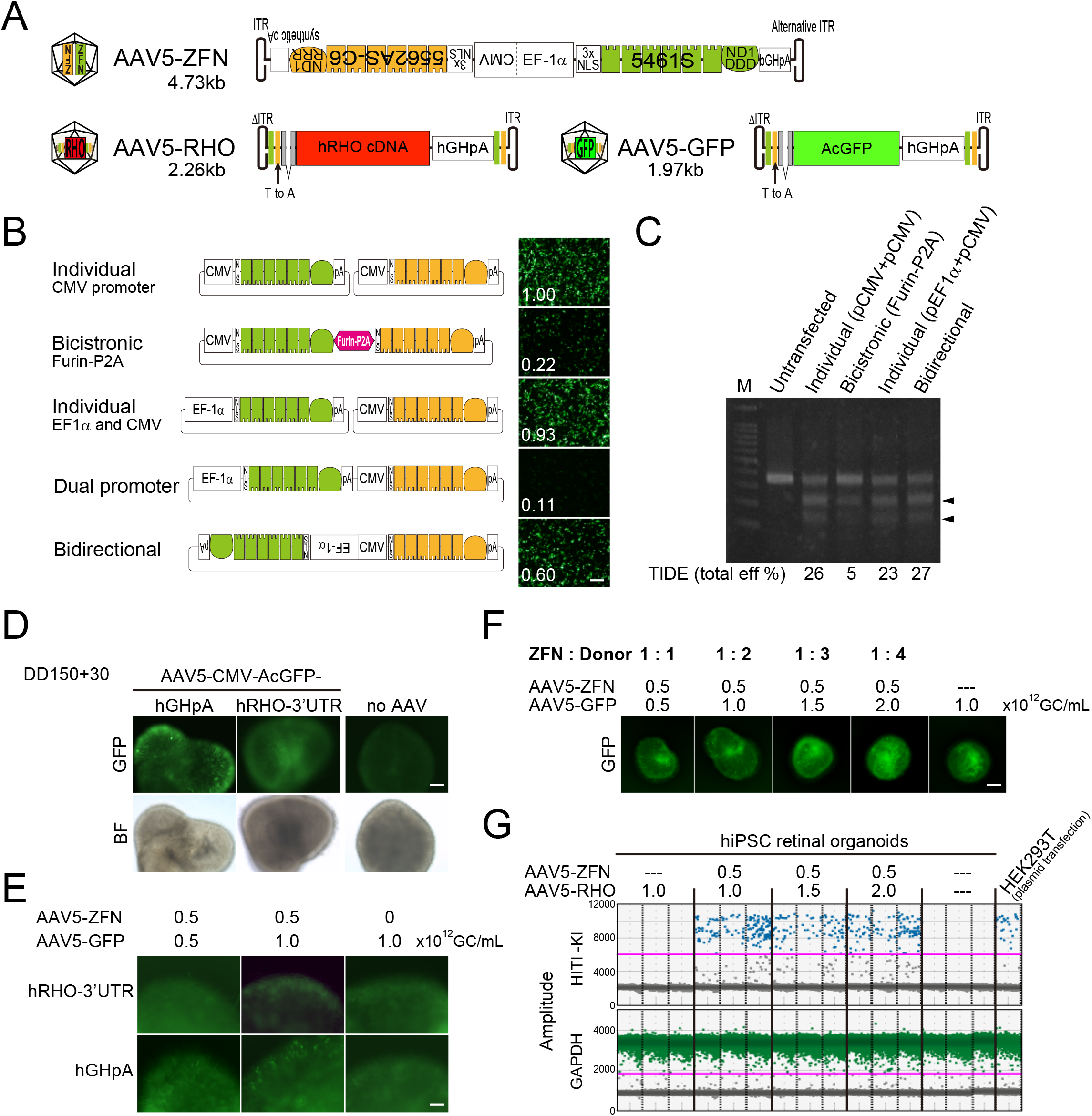
AAV Vector Optimization for HITI-Mediated Editing in Human Retinal Cells. **(A)** Final optimized AAV vector pair schematics. AAV5-ZFN (4.73 kb): bidirectional EF-1α/CMV promoter driving 5461S-ND1-DDD and 5562AS-C6-ND1-RRR; the 5′ ITR has an 11-bp deletion for efficient AAV packaging (Supplementary Table S1). AAV5-RHO (2.26 kb, scAAV): Δ-ITR–[ZFN target]–Chimeric intron–hRHO CDS–hGH polyA–[ZFN target]–ITR. AAV5-GFP (1.97 kb, scAAV): identical architecture with AcGFP replacing hRHO CDS (PoM verification vector). **(B)** Schematic of five promoter configurations for dual ZFN subunit co-expression: Individual CMV (two separate plasmids), Bicistronic Furin-P2A (single plasmid), Individual EF-1α and CMV (two plasmids), Dual promoter, and Bidirectional EF-1α/CMV (single plasmid, opposing orientations). PolyA signals indicated. Right: representative SSA fluorescence images for each configuration. Normalized EGFP:mCherry ratios (relative to Individual CMV = 1.00) are shown in the lower-left corner of each panel. Scale bar = 500 µm (SSA). **(C)** Cel-1 (Surveyor) nuclease assay for four promoter configurations. Lanes: M, ladder; Untransfected, no-plasmid control; conditions as indicated. Arrowheads indicate cleavage bands. Bottom: TIDE indel efficiencies (total eff %). **(D)** PolyA signal optimization in hiPSC-derived retinal organoids (DD150–200). From left to right: organoids transduced with AAV5-CMV-AcGFP-hGHpA (internal transduction control, confirming AAV5 delivery), AAV5-ZFN + AAV5-GFP carrying hRHO 3′-UTR, or untreated. GFP (top row) and BF (bottom row) images at DD150+30. Scale bar = 500 µm. **(E)** Comparison of hRHO 3′-UTR and hGH polyA in hiPSC-derived retinal organoids. Organoids transduced with AAV5-ZFN (0.5 × 10¹² GC/mL) and AAV5-GFP at 0.5 or 1.0 × 10¹² GC/mL, or AAV5-GFP alone (no ZFN), carrying either hRHO 3′-UTR (upper row) or hGH polyA (lower row). GFP images at DD150+30. Scale bar = 200 µm. **(F)** ZFN:donor ratio optimization in hiPSC-derived retinal organoids (DD150–200). Organoids transduced with AAV5-ZFN (0.5 × 10¹² GC/mL) and AAV5-GFP at 0.5–2.0 × 10¹² GC/mL (ZFN:GFP = 1:1 to 1:4), or AAV5-GFP alone (negative control). GFP fluorescence images at 30 days post-transduction. Scale bar = 200 µm. **(G)** ddPCR amplitude plots showing HITI-mediated knock-in allele frequency. Upper panels: hiPSC-derived retinal organoids transduced with AAV5-ZFN and AAV5-RHO at the indicated ratios, or AAV5-RHO alone. Right: HEK293T cells co-transfected with AAV5-ZFN and AAV5-RHO plasmids. FAM-positive droplets (upper cluster) = KI-positive; HEX-positive droplets = *GAPDH* internal control.

First, we screened 34 candidate ZF pairs targeting the human *RHO* 5′-UTR by SSA assay and identified 5461S + 5562AS as the lead candidate for downstream refinement. The SSA assay format and HITI workflow were established and validated using SpCas9 at the mouse *Rho* 5′-UTR, where HITI-mediated donor insertion was achieved in 80–90% of electroporated rod photoreceptor cells[23]. Of the 34 candidate pairs screened, 10 produced detectable EGFP reconstitution above background (Fig. 1B); the remaining 24 expressed the mCherry transfection control but generated no discernible GFP signal and were considered non-cleaving (data not shown). Among the 10 active pairs, 5 carried 5-bp spacers and 5 carried 6-bp spacers. Pairs with 5-bp spacers showed a tendency toward stronger GFP reconstitution; however, this trend was not consistent across all candidates and appeared dependent on the specific ZF–target sequence combination rather than a general property of spacer length. Among the 5-bp spacer group, the pairs 5461S + 5562AS and 5865AS + 5764S produced stronger GFP reconstitution signal, comparable to the SpCas9/mRho gRNA1 positive control. The 5461S + 5562AS combination was selected for downstream refinement based on its high SSA activity.

Next, we optimized the 5562AS antisense subunit based on concordant superiority across SSA, Cel-1, and TIDE analyses. Eight variants of the 5562AS antisense subunit (C1–C8) were evaluated in combination with 5461S (Fig. 1C–D, Supplementary Fig. S1). SSA assay identified C6 and C8 as the two variants with the strongest GFP reconstitution, with Cel-1 assay confirming prominent cleavage bands for both. TIDE analysis yielded indel frequencies for C1 through C8, and C6 achieved the highest TIDE efficiency, exceeding all other variants. Based on these results, 5461S + 5562AS-C6 was selected as the final ZF-ND1 pair, hereafter referred to as ZF-ND1 (FirmCut ND1).

Genome-wide off-target profiling by GUIDE-seq[33] in HEK293T and ARPE19 cells identified no candidate off-target sites with known photoreceptor function for ZF-ND1; comprehensive analysis of the complete GUIDE-seq dataset is ongoing and will be reported in the final manuscript. Of note, the alternative candidate pair 5865AS + 5764S, which also showed high SSA activity in the initial screen, was deprioritized following GUIDE-seq analysis that revealed a substantially higher number of candidate off-target sites. These findings further support the selection of ZF-ND1 as the lead pair. Amplicon-seq-based off-target validation in hiPSC-derived retinal organoids is also in progress.

We then evaluated NLS configurations[34] for efficient nuclear import in post-mitotic rod photoreceptors and identified 3×multiNLS as the optimal configuration. Because mature rod photoreceptors are terminally differentiated post-mitotic cells, nuclear import of ZFN protein must depend on active NLS-mediated transport. Comparison of four NLS configurations by *in vivo* electroporation in the mouse retina showed that nuclear import efficiency was dependent on NLS multiplicity and composition (Supplementary Fig. S2). The 1×SV40 NLS configuration yielded minimal HITI-mediated AcGFP signal, with progressive improvement observed with 3×SV40 NLS and 3×multiNLS (SV40–AAA–cMyc–GSG–SV40). The 5×multiNLS configuration did not further improve efficiency, consistent with the established requirement for redundant NLS sequences in post-mitotic neurons. The 3×multiNLS configuration was therefore incorporated into the final ZF-ND1 cassette.

### AAV Vector Optimization for HITI-Mediated Editing in Human Retinal Cells

Adeno-associated virus (AAV) is the preferred delivery vector for subretinal injection in post-mitotic photoreceptors, and AAV serotype 5 (AAV5) was selected based on its well-characterized high transduction efficiency to photoreceptor cells in the outer nuclear layer[35,36]. Using hiPSC-derived retinal organoids as a human cellular platform, we optimized three parameters for clinically preferred vector design: (1) promoter architecture for dual ZFN subunit co-expression, (2) the 3′ regulatory element for self-complementary AAV (scAAV) packaging, and (3) the ZFN:donor ratio. The resulting constructs are shown schematically in Fig. 2A.

Prior to vector optimization, a T-to-A transversion was introduced, corresponding to the third base of the second ZF module of 5562AS-C6, within the donor vector on the 5′-ITR-proximal side. Because the donor vector is designed with the ZF-ND1 recognition sequence in reverse orientation to enable directional HITI insertion, correct genomic integration would expose an ATG start codon on the sense strand at this position. The T base at this site exhibits low specificity based on position weight matrix (PWM) prediction (Interactive PWM Predictor; https://zf.princeton.edu/)[37,38], making it amenable to substitution without compromising ZF-ND1 binding. In fact, the T-to-A transversion eliminates the spurious ATG while retaining cleavage efficiency equivalent to the unmodified sequence, as confirmed by SSA assay targeting the donor vector sequence (Supplementary Fig. S3B).

We first evaluated multiple vector architectures for co-expression of ZF-ND1 subunits and found that the bidirectional EF-1α/CMV configuration enabled co-expression of both ZF-ND1 subunits from a single AAV vector while maintaining cleavage efficiency comparable to the individual promoter systems requiring two separate plasmids. Individual CMV and EF-1α/CMV configurations achieved indel efficiencies of 26% and 23%, respectively (Fig. 2B–C, Supplementary Fig. S4). By contrast, the Bicistronic Furin-P2A configuration yielded a substantially lower efficiency of 5%, reproduced across two independent experiments. We attribute this reduction to the residual peptide sequence remaining on the ND1 domain following 2A-based self-cleavage, which is predicted to interfere with ND1-DDD/ND1-RRR obligate heterodimerization[39]. The Bidirectional EF-1α/CMV configuration was adopted for the final AAV5-ZFN vector. Of note, the lower SSA ratio observed for the Bidirectional configuration (0.60 relative to Individual CMV = 1.00) likely reflects plasmid context-specific constraints on bidirectional transcription, rather than reduced genomic editing capacity as confirmed by Cel-1 and TIDE.

We next optimized the 3’ regulatory element for the donor construct. In the mouse system, the 3’-UTR of *Rho* supported highly efficient donor gene expression[23,40,41]; however, human growth hormone (hGH) polyA outperformed hRHO 3’-UTR in hiPSC-derived retinal organoids and enabled scAAV packaging. hRHO 3’-UTR yielded minimal signal above the no-AAV control despite confirmed transduction by the AAV5-CMV-AcGFP internal control, whereas hGH polyA produced substantially stronger AcGFP fluorescence (Fig. 2D–E). In addition, hGH polyA reduced the donor cassette to 2.26 kb (AAV5-RHO) or 1.97 kb (AAV5-GFP), both within the scAAV packaging limit of approximately 2.3 kb. As scAAV delivers a self-complementary dsDNA genome that bypasses the rate-limiting second-strand synthesis required for single-stranded AAV (ssAAV)[42,43], ZF-ND1-mediated cleavage is expected to initiate earlier following transduction, potentially enhancing HITI efficiency. hGH polyA was therefore adopted as the 3’ regulatory element for all subsequent donor vectors.

hiPSC-derived retinal organoids retinal organoids (DD150–200) were transduced with AAV5-ZFN and increasing concentrations of AAV5-GFP (ZFN:GFP = 1:1 to 1:4; Fig. 2F). Droplet digital PCR (ddPCR) analysis confirmed HITI-mediated donor insertion at all tested ZFN:donor ratios in hiPSC-derived retinal organoids (Fig. 2G). KI allele frequency was below 1% (approximately 0.3–0.5%) under all conditions tested, with no clear dose-dependent increase observed across ZFN:donor ratios, suggesting the lower AAV transduction efficiency achievable in three-dimensional organoid cultures compared with *in vivo* delivery. The robust KI allele frequency confirmed by ddPCR was comparable to that to HEK293T cells co-transfected with AAV5-ZFN and AAV5-RHO plasmids at a ZFN:RHO ratio of 1:1 (Fig. 2G). This value is therefore not used as a primary efficacy metric; efficacy is instead addressed by the rat and NHP experiments (Figs. 3 and 4). The ZFN:RHO ratio of 1:1 was not included in the organoid experiment owing to limited availability of the AAV5-RHO preparation at the time of the assay.

**Figure 3.**
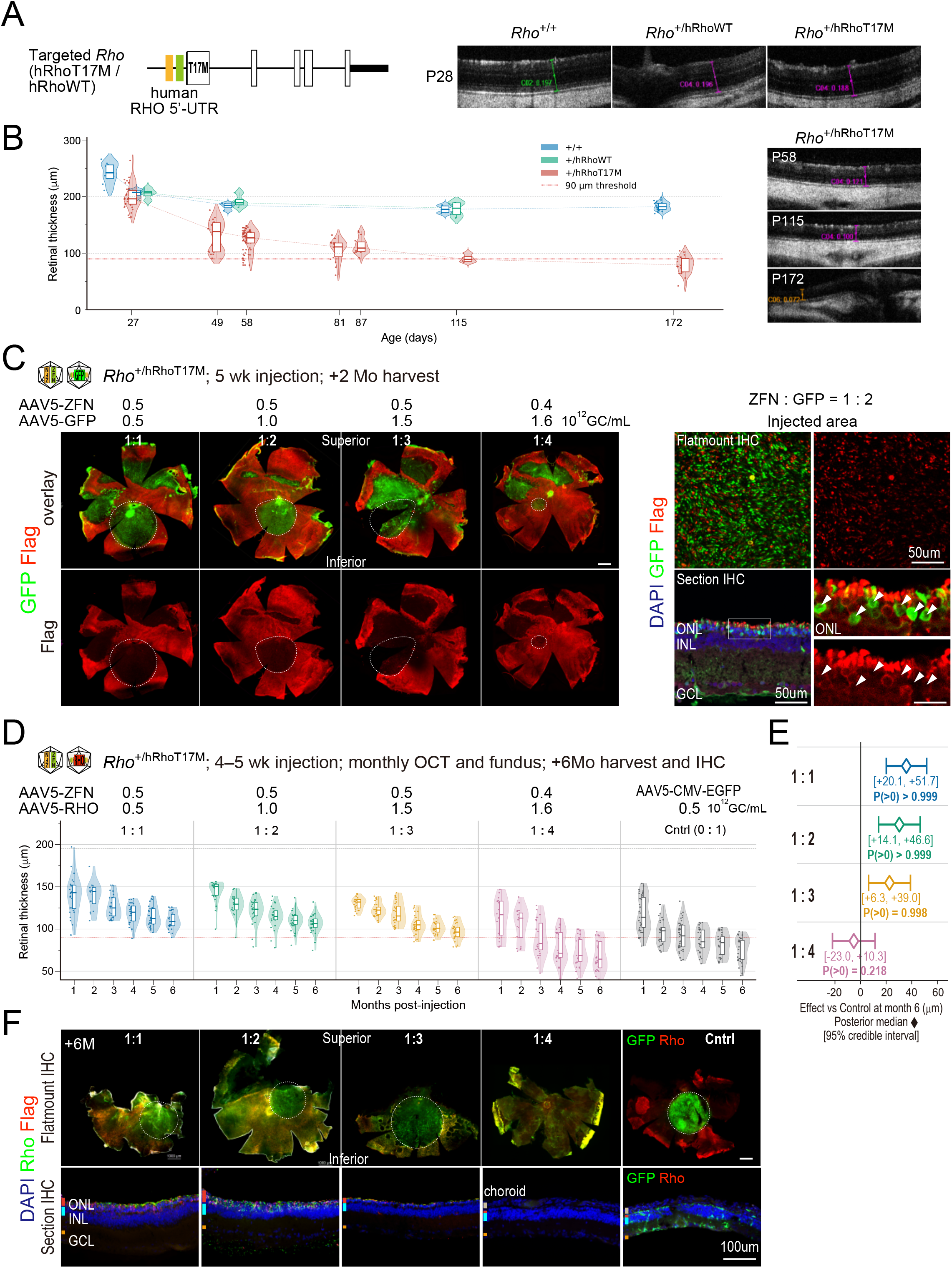
Therapeutic Efficacy of ZF-ND1-Mediated HITI in a Humanized RHO-T17M Rat Disease Model. **(A)** Experimental scheme and OCT images. Left: allele structure of targeted Rho locus in *Rho*^+/hRhoT17M^ and *Rho*^+/hRhoWT^ rats, showing the human RHO 5′-UTR cassette (containing ZF-ND1 target site, FLAG tag, and T17M mutation) knocked into endogenous rat Rho exon 1. Right: representative OCT B-scan images from *Rho*^+/+^, *Rho*^+/hRhoWT^, and *Rho*^+/hRhoT17M^ animals at P28. Calipers indicate ONL thickness measurement. **(B)** Natural history of retinal thickness. Left: violin plots showing ONL thickness (µm) in *Rho*^+/+^ (blue), *Rho*^+/hRhoWT^ (teal), and *Rho*^+/hRhoT17M^ (red) animals at the indicated postnatal days (n = 3–4 per genotype; 7 OCT measurements per animal per time point; postnatal days 58 and 59 combined). Red dotted line: 90 µm threshold. *Rho*^+/hRhoWT^ was not significantly different from *Rho*^+/+^ at any time point (Wilcoxon rank-sum test, all P > 0.05). Right: representative OCT B-scans of *Rho*^+/hRhoT17M^ at P58, P115, and P172 showing progressive ONL thinning. **(C)** HITI-mediated knock-in verification at 2 months. *Rho*^+/hRhoT17M^ retinas injected with PoM verification vector (AAV5-ZFN + AAV5-GFP) at ZFN:GFP = 1:1 to 1:4. Upper row: flatmount overlay (GFP green, FLAG red); lower row: FLAG channel only. Dashed circles indicate injection area. GFP-positive cells within the injection area show loss of FLAG signal. Right: high-magnification flatmount IHC (ZFN:GFP = 1:2 representative) and section IHC showing GFP (green) and FLAG (red) in the ONL. Scale bar = 50 µm. n = 2 eyes per group. **(D)** Longitudinal ONL thickness in *Rho*^+/hRhoT17M^ animals treated with AAV5-ZFN + AAV5-RHO at ZFN:RHO = 1:1, 1:2, 1:3, or 1:4 (n = 4 per group) or Control (AAV5-CMV-EGFP + AAV5-RHO, no ZFN; n = 6). Violin plots show all 7 OCT measurements per animal per time point. Red dotted line: 90 µm threshold. ZFN:RHO 1:4: n = 3 animals at M1–M2 (failed injection in one animal). **(E)** Bayesian posterior probability analysis of treatment effect at month 6. Points (L) indicate posterior medians; horizontal bars indicate 95% credible intervals (CrI). Effect sizes relative to Control: ZFN:RHO 1:1, +35.8 µm (95% CrI +20.1 to +51.7; P(>0) > 0.999); 1:2, +30.3 µm (+14.1 to +46.6; P(>0) > 0.999); 1:3, +22.6 µm (+6.3 to +39.0; P(>0) = 0.998); 1:4, −6.3 µm (−23.0 to +10.3; P(>0) = 0.218). Bayesian LMM: thickness L group × month + (1|animal); 4 chains × 2,000 MCMC samples; R-hat ≤ 1.002. **(F)** Long-term HITI donor expression at 6-month harvest. Upper row: retinal flatmount IHC (GFP green, RHO red) from AAV5-ZFN + AAV5-RHO-treated *Rho*^+/hRhoT17M^ animals (ZFN:RHO = 1:1, 1:2, 1:3, 1:4) and Control. Dashed circles indicate injection area. GFP-positive cells are detected within the injection area in the 1:1 and 1:2 groups. Lower row: section IHC (DAPI blue, GFP green, RHO red) showing GFP-positive photoreceptors in the ONL in treated animals. Scale bar = 100 µm. Antibodies: anti-GFP (rat monoclonal, Nacalai Tesque cat# 04404-84, 1:1000); anti-Rhodopsin (rabbit polyclonal, Abcam ab112576, 1:2000); DAPI.

**Figure 4.**
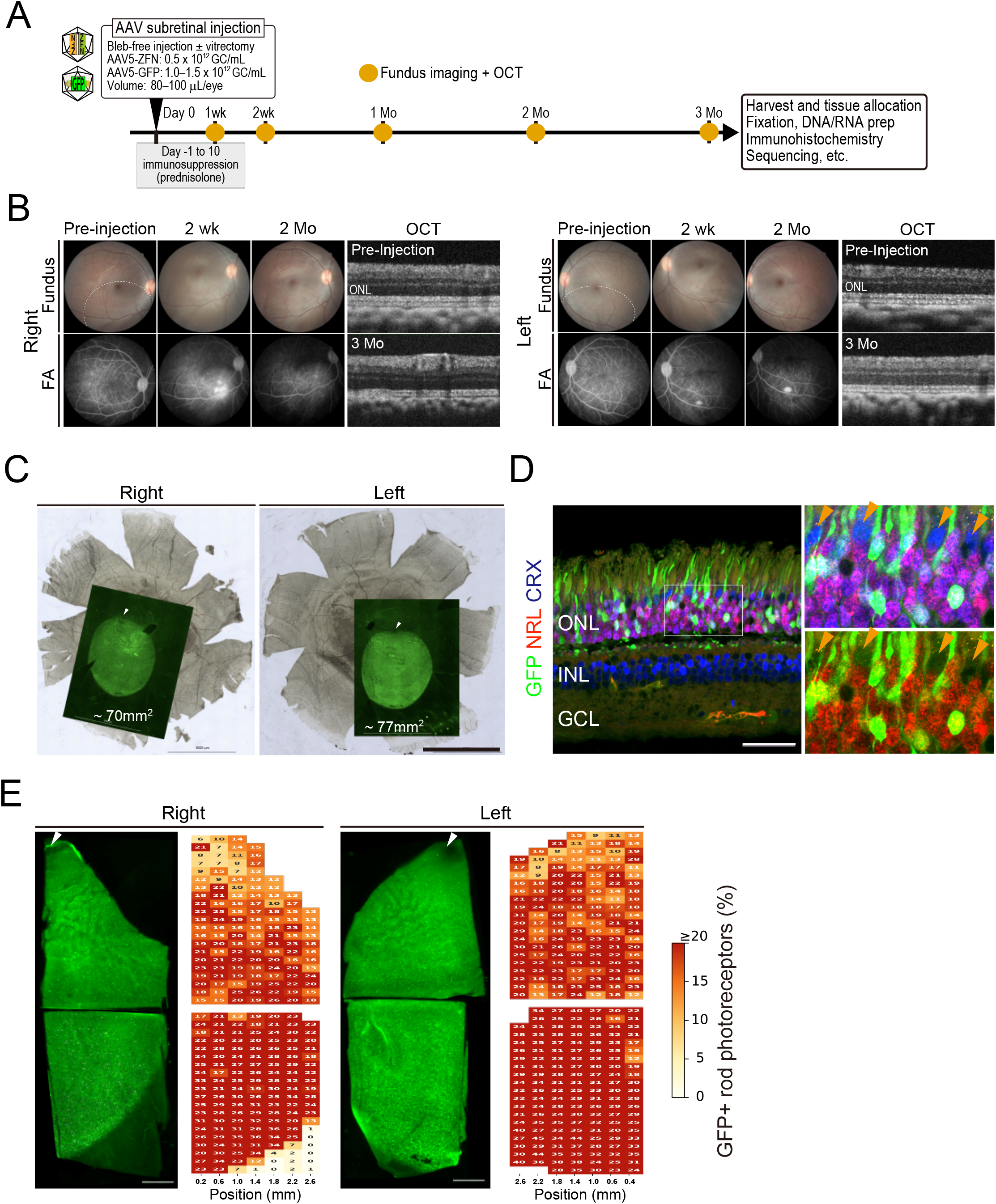
Proof-of-Mechanism for ZF-ND1-Mediated HITI Editing in the Crab-Eating Monkey Retina. **(A)** Experimental scheme. Two crab-eating monkeys (NHP1, ZFN:GFP = 1:3; NHP2 ZFN:GFP = 1:2) received bilateral subretinal injection of PoM verification vector (AAV5-ZFN + AAV5-nGFP; 100 µL per eye) without prior vitrectomy. Immunosuppression: prednisolone 1 mg/kg/day i.m. (Day −1 to Day 3) tapered to 0.5 mg/kg/day (Day 4–10), plus triamcinolone acetonide sub-Tenon injection (STTA) at the day of injection and Days 14, 28, and 56. Retinas were harvested at 3 months post-injection for flatmount fluorescence imaging, IHC, and genomic DNA/RNA extraction. **(B)** Fundus photographs, fluorescein angiograms (FA), and OCT images from NHP1 at pre-injection baseline and at post-injection months 1 and 2. FA at 2 months post-injection confirms successful subretinal delivery (bleb area) with no active inflammation in either eye. OCT shows preserved ONL structure through 3 months in both eyes. **(C)** Retinal flatmount fluorescence images showing GFP+ (AcGFP) photoreceptors within the injection area at 3-month harvest. Imaging was restricted to the GFP-positive injection area; full-flatmount imaging was not performed to preserve tissue quality for DNA/RNA analysis. Total retinal area: left eye ∼77 mm², right eye ∼70 mm². GFP+ cells are visible within the injection area. Scale bar = 10 mm. **(D)** Immunohistochemistry of retinal cryosections from the injection area. Triple-label staining: GFP (green; HITI-mediated GFP donor insertion), NRL (red; rod photoreceptors), CRX (blue; all photoreceptors including cones). GFP+/NRL+ cells confirm HITI editing in rod photoreceptors. CRX+/NRL− cone photoreceptors are GFP-negative. Scale bar = 50 µm. **(E)** Spatial heatmap of GFP+/NRL+ photoreceptor fraction (%) overlaid on the retinal flatmount. Each value represents the GFP+/NRL+ fraction from one immunostained cryosection sampled at 100 µm intervals across the GFP-positive area; sections correspond to the tissue fragments shown in the flatmount. Arrowheads indicate the approximate macular region. The red dashed line (20%) indicates the therapeutic correction threshold, defined as the minimum GFP+/NRL+ fraction required for clinically meaningful retinal preservation in heterozygous RHO-adRP (see Methods).

Sanger sequencing of KI junction PCR products from hiPSC-derived retinal organoids (ZFN:GFP = 1:2, 1:3, 1:4) and MeWo melanoma cells (ZFN:GFP = 1:1; included as a non-photoreceptor control) confirmed HITI-mediated donor insertion at the predicted genomic target site in all samples (Supplementary Fig. S5). The two most frequent junction patterns were 0-bp precise ligation and 1-bp deletion at the ZFN cleavage site on the genomic side, both consistent with NHEJ-mediated end-joining following ZF-ND1-induced cleavage. Neither pattern disrupts the downstream reading frame due to the chimeric intron design[44–47], and the consistent junction profiles across cell types and ZFN:donor ratios indicate consistent ZF-ND1-mediated HITI integration. KI junction sequencing in NHP retinal tissue is described in the following section (Fig. 4).

### Therapeutic Efficacy of ZF-ND1-Mediated HITI Editing in a Humanized RHO-T17M Rat Disease Model

To evaluate therapeutic efficacy *in vivo*, we generated a humanized rat disease model carrying the human *RHO* 5′-UTR with the T17M mutation[48–50] knocked into the endogenous *Rho* locus (Supplementary Fig. S6). The T17M mutation is associated with a severe, early-onset phenotype in patients[17,51]. We assessed the therapeutic vector in a 6-month longitudinal study with monthly OCT monitoring (Fig. 3A). We replaced endogenous rat *Rho* exon 1 with the human *RHO* 5′-UTR (including the ZF-ND1 target sequence) and T17M mutation, generating *Rho*^+/hRhoT17M^ and *Rho*^+/hRhoWT^ lines. Serial OCT monitoring showed progressive outer nuclear layer (ONL) thinning in *Rho*^+/hRhoT17M^ animals after 4 weeks of age, reaching ∼80 µm by P115 (Fig. 3B). Based on this natural history, an ONL thickness of ≤90 µm by OCT was defined as the criterion for near-complete photoreceptor loss in this model. *Rho*^+/hRhoWT^ animals maintained retinal thickness equivalent to wild-type (*Rho*^+/+^) at all time points (Wilcoxon rank-sum test, all P > 0.05), confirming that humanization without T17M does not cause degeneration. One month of age was therefore established as the optimal intervention window.

Prior to the proof-of-concept study, we determined the working dose range of the AAV cocktail. *Rho*^+/hRhoWT^ animals were injected subretinally with the proof-of-mechanism (PoM) verification vector (AAV5-ZFN + AAV5-GFP; ZFN:GFP = 1:1) at three dose levels and assessed at 2 months post-injection. Retinal flatmounts showed GFP-positive rod photoreceptors within the injection area at all doses tested; the dose of 1.0 × 10¹² GC/mL yielded clear GFP signal and was selected as the working dose for subsequent experiments (Supplementary Fig. S7). Subsequent experiments were therefore conducted at doses within this order of magnitude. In addition, we performed PoM verification experiments prior to therapeutic assessment and confirmed simultaneous HITI-mediated knock-in and mutant RHO silencing in the HITI-edited rod photoreceptors. *Rho*^+/hRhoT17M^ animals injected with PoM verification vector (AAV5-ZFN + AAV5-GFP; ZFN:GFP = 1:1 and 1:2) showed GFP-positive rod photoreceptors within the injection area at 2 months post-injection, with loss of FLAG signal (T17M mutant hRHO) in GFP-positive cells (Fig. 3C; n = 2 eyes per group), confirming the mechanism of action at cellular resolution.

We then assessed the therapeutic efficacy of the therapeutic vector. The therapeutic vector suppressed retinal degeneration over 6 months, with ZFN:RHO ratios of 1:1 and 1:2 achieving ONL protection above the 90 µm threshold. *Rho*^+/hRhoT17M^ animals were injected with the therapeutic vector (AAV5-ZFN + AAV5-RHO; ZFN:RHO = 1:1 to 1:4) at 4 weeks of age and monitored monthly by OCT for 6 months (Fig. 3D). At month 6, the 1:1 group maintained a median ONL of 109 µm (n = 4, 28 measurements; range 90–126 µm) and the 1:2 group maintained 107 µm (n = 4, range 75–132 µm), both above the 90 µm threshold. The 1:3 group maintained 97 µm (range 79–115 µm) with greater inter-individual variation, while the 1:4 group declined to 65 µm (range 41–97 µm), comparable to Control (80 µm; n = 6, range 45–97 µm). Bayesian posterior probability analysis confirmed significant preservation in the 1:1, 1:2, and 1:3 groups relative to Control, whereas the 1:4 group showed no credible positive effect (Fig. 3E). Effect sizes at month 6: 1:1 group, +35.8 µm (95% CrI +20.1 to +51.7; P(>0) > 0.999); 1:2, +30.3 µm (+14.1 to +46.6; P(>0) > 0.999); 1:3, +22.6 µm (+6.3 to +39.0; P(>0) = 0.998); 1:4, −6.3 µm (−23.0 to +10.3; P(>0) = 0.218). The 1:1 and 1:2 groups had 95% credible interval (CrI) entirely above zero. These results suggest that ZFN:RHO ratios of 1:1 and 1:2 are the optimal dosing regimens.

At the 6-month endpoint, retinal flatmount and cryosection immunohistochemistry (IHC) confirmed long-term structural preservation of the ONL in treated animals (Fig. 3F). Rhodopsin immunoreactivity was detected throughout the injection area in the 1:1 and 1:2 groups, with preservation of ONL laminar architecture on cryosections. The 1:3 group showed intermediate signal with greater spatial heterogeneity, while the 1:4 group and Control (AAV5-CMV-EGFP) showed markedly reduced rhodopsin immunoreactivity and ONL thinning, consistent with advanced photoreceptor degeneration. These data confirm that the structural preservation observed by OCT is accompanied by sustained photoreceptor integrity at the cellular level through 6 months post-injection. Note that the superior retina could not be flatmounted in any group, owing to a combination of severe thinning and adhesion to the retinal pigment epithelium (RPE). For the 1:4 group and Control, dissection of the inferior retina was also technically challenging; flatmounts were therefore prepared by choroidal dissection after scleral removal, which may result in apparent overestimation of retinal thickness on DAPI staining.

In this descriptive safety assessment, the therapeutic vector did not cause progressive retinal thinning in wild-type rats over 5 months of observation at either dose level tested. Wild-type Wistar rats injected at the standard dose (1.5 × 10¹² GC/mL total) and one-third dilution (0.5 × 10¹² GC/mL) maintained ONL thickness of 173–211 µm throughout 5 months of observation (Supplementary Fig. S8), comparable to those of non-treated WT rats. Representative OCT (months 2–5) confirm preservation of normal retinal laminar architecture. These data confirm the absence of vector-associated retinal toxicity at the doses tested[52].

### Proof-of-Mechanism for ZF-ND1-Mediated HITI Editing in the NHP Retina

To extend preclinical validation to a non-human primate model, we assessed proof-of-mechanism activity of ZF-ND1-mediated HITI editing in crab-eating monkey retina, which shares structural and cellular organization with the human retina[36].

Prior to the experiment, we confirmed ZF-ND1 cleavage activity at the crab-eating monkey *RHO* 5′-UTR. Sequence alignment revealed three nucleotide mismatches between the human and crab-eating monkey target sequences in the 5461S recognition region (Supplementary Fig. S3A); PWM-based prediction indicated that the mismatch at the third position of ZF73 (C→A), which corresponds to a high-specificity position in the ZF recognition motif, is expected to reduce cleavage activity. SSA assay confirmed an approximately 30–40% reduction in GFP reconstitution efficiency for the NHP target relative to the human sequence (Supplementary Fig. S3B). This level of cleavage activity was considered sufficient for proof-of-mechanism validation, which was the primary objective of the NHP experiment.

We then administered PoM vectors (AAV5-ZFN + AAV5-GFP) to both eyes of NHP1 (ZFN:GFP = 1:3) and NHP2 (ZFN:GFP = 1:2) by subretinal injection (∼100 µL/eye) (Fig. 4A). The bleb-free direct subretinal delivery method was adopted based on our prior study in which the same approach without vitrectomy was validated in crab-eating monkeys. Injection was targeted to the paramacular and mid-peripheral inferior retina, as visualization of the peripheral retina was limited by the available surgical microscope.

Serial fundus, fluorescein angiography (FA), and OCT examinations confirmed retinal integrity through 3 months in NHP1 (Fig. 4B), while FA findings in NHP2 indicated intraocular inflammation in the injection area (Supplementary Fig. S9). Fundus examination and FA at post-injection months 1 and 2 confirmed successful subretinal delivery in both animals. FA at 2 months showed no evidence of active intraocular inflammation in NHP1 in either eye. In NHP2, we observed chorioretinal changes and demarcation within the injection area in fundus images, and FA revealed findings consistent with intraocular inflammation. OCT imaging through 3 months confirmed preservation of ONL structure in both animals, with no progressive thinning attributable to vector toxicity. The inflammatory data for NHP2 are retained as a reference observation and presented separately in Supplementary Fig. S9.

At the 3-month harvest, GFP-positive photoreceptors were detected across the injection area in both animals, and triple-label IHC confirmed rod-selective HITI integration. Retinal flatmount IHC staining confirmed GFP+ photoreceptors within the injection area in both animals (Fig. 4C). Triple-label immunohistochemistry (GFP/NRL/CRX) on cryosections confirmed GFP+/NRL+ rod photoreceptors within the HITI-edited area (Fig. 4D). GFP signal was absent in CRX+/NRL− cone photoreceptors, confirming rod-selective HITI integration.

Spatial heatmap quantification of GFP+/NRL+ photoreceptor fractions confirmed HITI editing across multiple retinal sections of NHP1 (Fig. 4E). Sections were sampled at 400 µm intervals and results overlaid on retinal flatmount images. A tendency toward lower GFP+/NRL+ fraction was observed in sections closer to the macula, consistent with the rod-selective expression of the inserted GFP cassette driven by the endogenous *RHO* promoter, which is not active in cone photoreceptors. In NHP1, GFP+/NRL+ fractions exceeding 20% were observed in many sections in both eyes, with comparable spatial distribution between the right and left eyes. In NHP2, GFP+ fractions were reduced in areas corresponding to FA-visible inflammatory changes, while sections outside these areas showed editing levels comparable to NHP1.

These editing levels were evaluated against a therapeutic correction threshold of 20%, defined based on the minimum photoreceptor preservation required to maintain visual function[53,54] (≥10%; Geller & Sieving, *Vision Res*, 1993; Geller et al., *J Opt Soc Am A*, 1992), multiplied by a factor of two to account for the heterozygous dominant-negative mechanism of RHO-adRP. In NHP1, the majority of sections exceeded this threshold, suggesting that ZF-ND1-mediated HITI editing can achieve therapeutic-level photoreceptor correction in the primate retina.

Genomic and cDNA sequencing confirmed NHEJ-mediated HITI integration and active transcription of the inserted cassette in NHP retinal tissue. Sanger sequencing of KI junction PCR products from NHP retinal tissue (ZFN:GFP = 1:3) confirmed HITI-mediated donor insertion at the predicted genomic target site (Supplementary Fig. S5). As in human cells, the two most frequent junction patterns were 0-bp precise ligation and 1-bp deletion; however, a higher proportion of 1-bp deletion junctions was observed in NHP compared with hiPSC-derived retinal organoids and MeWo cells, consistent with the three-nucleotide mismatch in the 5461S recognition region possibly causing a subtle shift in ZF array binding position and thereby altering the geometry of ND1-mediated cleavage at the NHP 5′-UTR. RT-PCR of retinal RNA from the injection area confirmed the presence of a correctly spliced HITI-derived cDNA transcript, with chimeric intron processing at the predicted splice acceptor site and seamless continuity into the inserted *hRHO*/GFP cassette (Supplementary Fig. S10). A minor sequence deviation was observed in the 5′-UTR region, consistent with the same three-nucleotide mismatch possibly causing a subtle shift in ZF-ND1 cleavage geometry; this pattern is expected to be absent when ZF-ND1 is administered to patients carrying the fully matched human *RHO* sequence. These findings confirm stable HITI integration, correct chimeric intron processing, and active transcription of the inserted *hRHO* cassette in NHP rod photoreceptors.

## Discussion

HITI was originally established by Suzuki et al.[20] using a blunt-end-cutting CRISPR-Cas9, exploiting non-homologous end joining (NHEJ), the predominant DNA repair pathway in post-mitotic cells, to achieve stable donor insertion in terminally differentiated tissues. In this system, the donor cassette is flanked at both ends by the nuclease target sequence in reverse orientation. Donor cassettes integrated in the reverse orientation regenerate the target site at both junctions and are re-cleaved and removed, while those integrated in the forward orientation disrupt the target site and are stably retained, thereby enriching for productive integration. This ObLiGaRe-type architectural constraint[22] enforces integration directionality independently of the initial cleavage geometry. The original ObLiGaRe study reported that 0-bp precise junctions were the most frequent outcome across multiple ZFN-targeted genomic loci, with minor short deletions or insertions in a subset of clones. KI junction sequencing from human-derived samples (hiPSC retinal organoids and MeWo cells) showed 0-bp precise ligation and 1-bp deletion as the two most frequent junction patterns across all conditions tested (Supplementary Fig. S5), confirming that ZF-ND1 generates double-strand breaks (DSBs) compatible with productive NHEJ-mediated HITI regardless of the precise cut geometry. In crab-eating monkey retina, where three nucleotide mismatches in the 5461S recognition region shift the ZF array binding position, a higher proportion of 1-bp deletion junctions was observed relative to human-derived samples, consistent with an altered ZF-DNA binding at the NHP 5′-UTR. These results are consistent with the prediction that the fully matched human RHO sequence will generate the canonical cleavage and integration pattern expected in patients.

The *in vitro* screening strategy underlying ZF-ND1 selection is itself supported by precedent from the prior SpCas9-based HITI system, which validates SSA cleavage efficiency as a reliable selection criterion. We previously reported that gRNA/Cas9 complexes with higher *in vitro* SSA cleavage efficiency consistently demonstrated superior *in vivo* HITI insertion efficiency in mouse rod photoreceptors[23]; the mouse Rho-targeting ZF-ND1 pair 5057S/AS, which exhibited lower SSA efficiency relative to the high-performing SpCas9 gRNAs, similarly showed reduced *in vivo* insertion efficiency by *in vivo* electroporation. This parallel validates the use of SSA efficiency as the primary selection criterion in the human ZF-ND1 screening campaign in the present study.

The contribution of hiPSC-derived retinal organoids to this preclinical dataset warrants careful interpretation. The KI allele frequency measured by ddPCR (0.3–0.5%) was substantially lower than the editing efficiency observed i*n vivo* in the NHP experiments. This discrepancy reflects the inherent limitation of AAV transduction efficiency in three-dimensional organoid cultures, where vector penetration and intercellular distribution differ fundamentally from the subretinal injection setting. This challenge is consistent with previous reports demonstrating that efficient AAV transduction of human retinal organoids requires engineered capsid variants and optimized delivery conditions, with wild-type AAV serotypes (including AAV5) achieving limited transduction of photoreceptor cells within the organoid interior[55,56]. In this study, organoids served as a platform for vector optimization and mechanism verification rather than as a quantitative efficacy model; primary efficacy assessment was therefore conducted in the rat disease model and NHP experiments, where clinically meaningful HITI editing levels were achieved.

The format of the donor vector provides a further design consideration relevant to HITI efficiency. Unlike ssAAV, scAAV delivers a self-complementary dsDNA genome that bypasses second-strand synthesis[42], enabling earlier ZF-ND1-mediated cleavage and donor linearization following transduction. This property, combined with the T-to-A transversion that preserves donor cleavage efficiency while reducing re-cleavage risk[57], is expected to support productive HITI even under conditions of reduced genomic cleavage efficiency, as observed in NHP.

Genomic specificity is an equally critical requirement for clinical translation. Following selection of 5461S + 5562AS based on cleavage activity, in silico off-target prediction using PROGNOS[58] identified a limited number of candidate sites in the human genome for this ZF-ND1 pair. The architectural properties of ZF-ND1, including the extended target recognition sequence of up to 36 bp per 6-finger pair and the obligate heterodimer constraint of the ND1 domain, are predicted to confer a high degree of genomic specificity. Unbiased experimental profiling of the off-target landscape by GUIDE-seq has been completed and will be reported in a forthcoming version of this manuscript. The therapeutic efficacy of ZF-ND1-mediated HITI was evaluated in a humanized rat disease model. The therapeutic vector is designed to arrest or decelerate progressive photoreceptor degeneration rather than to restore lost visual function. ONL thickness measured by serial OCT was selected as the primary efficacy endpoint, consistent with its increasing use as a primary structural endpoint in clinical trials for inherited retinal dystrophies[51]. OCT is particularly suited to evaluating subretinal gene therapy, where therapeutic transduction is confined to the injection site and local improvements are diluted in whole-field electrophysiological measures.

The PoM and proof-of-concept (PoC) experiments provide concordant evidence of a dose-dependent therapeutic mechanism. In the PoM experiment, injection of the verification vector (AAV5-ZFN + AAV5-GFP) at ZFN:GFP = 1:1 and 1:2 yielded GFP-positive rod photoreceptors broadly distributed across the injection area with concurrent loss of FLAG (T17M mutant hRHO) immunoreactivity in GFP-positive cells, confirming simultaneous HITI-mediated knock-in and mutant allele silencing at the cellular level. At ZFN:GFP = 1:3 and 1:4, the GFP-positive area was markedly reduced, indicating lower editing efficiency at higher donor ratios. This pattern was mirrored in the PoC experiment: ZFN:RHO = 1:1 and 1:2 maintained ONL thickness above the 90 µm preservation threshold throughout 6 months (Bayesian effect sizes +35.8 µm and +30.3 µm, respectively; P(>0) > 0.999 for both), while the 1:3 group showed statistically credible but variable preservation (+22.6 µm; P(>0) = 0.998; range 79–115 µm), and the 1:4 group showed no credible treatment benefit (−6.3 µm; P(>0) = 0.218). At the 6-month endpoint, retinal flatmount and cryosection immunohistochemistry confirmed sustained rhodopsin immunoreactivity and preserved ONL laminar architecture in the 1:1 and 1:2 groups, corroborating the OCT-based efficacy findings at the cellular level (Fig. 3F).

The parallel dose-dependent patterns across PoM and PoC experiments provide strong mechanistic linkage between HITI-mediated editing and structural photoreceptor preservation. ZFN:RHO = 1:2 is identified as the preferred ratio, achieving efficacy equivalent to 1:1 while minimizing vector load. The diminishing and ultimately absent efficacy at higher donor ratios is likely attributable to a combination of reduced AAV5-ZFN delivery per cell at a fixed total viral load and cellular stress responses induced by excess AAV genome, both of which would impair the DSB formation required for productive HITI.

The NHP PoM experiment extended this validation to the primate retina. The 20% therapeutic correction threshold applied in the NHP experiment was defined based on the minimum photoreceptor preservation required to maintain retinal function (≥10%)[53,54], multiplied by a factor of two to account for the heterozygous dominant-negative mechanism of RHO-adRP, in which disruption of a single allele per targeted cell is required for therapeutic benefit. In NHP1, subretinal delivery of the PoM verification vector achieved GFP+/NRL+ fractions exceeding 20% in the majority of retinal sections in both eyes, despite an approximately 30–40% reduction in ZF-ND1 cleavage efficiency at the crab-eating monkey RHO 5′-UTR resulting from the three-nucleotide mismatch in the 5461S recognition region described above. The higher cleavage activity expected in patients with the fully matched human RHO sequence will, however, also increase competition from NHEJ-mediated indel formation at the DSB site; the net HITI efficiency in patients will therefore depend on the balance between productive HITI and competing repair pathways, and cannot be assumed to scale linearly with cleavage efficiency alone. Nevertheless, the NHP data demonstrate that therapeutically relevant editing levels are achievable even under the suboptimal cleavage conditions of the NHP target sequence, providing a conservative lower-bound estimate for the correction expected in patients.

The ZF-ND1 target site in the RHO 5′-UTR lies within a region containing common single-nucleotide variants annotated in gnomAD v4.1.1[59]. HITI-mediated integration replaces the targeted 5′-UTR with a defined donor sequence, ensuring that germline polymorphisms at the target site do not persist in the post-editing locus configuration. The NHP experiments in this study provide direct empirical support for this design principle: therapeutic-level HITI editing was achieved despite the three-nucleotide sequence divergence between the human and crab-eating monkey target substrates, demonstrating that the therapeutic outcome is not compromised by local sequence variation at the target site.

The interpretation of the NHP efficacy data requires consideration of the inflammatory findings in NHP2. The response observed in NHP2, manifested as FA-visible chorioretinal changes in the injection area, is consistent with immune reactivity to the AAV5 capsid, potentially exacerbated by pre-existing neutralizing antibody titers (not assessed in this study). A similar finding was noted in the preclinical development of EDIT-101 for Leber congenital amaurosis type 10 (LCA10)[60]. Pre-operative screening for AAV5 serum neutralizing antibody titers will therefore be included in the clinical protocol to identify and appropriately manage patients at risk of immune-mediated attenuation of therapeutic efficacy.

The ZFN:donor ratio identified in the rat model provides an initial reference for clinical dose selection, though direct extrapolation to humans requires consideration of several species-specific factors. AAV transduction efficiency and biodistribution differ between rat subretinal injection and human subretinal delivery; the reduced genomic cleavage efficiency in NHP alters the relative demand for linearized donor substrate; and the larger retinal volume in humans introduces spatial heterogeneity in vector distribution not captured in the rat model. Consistent with this, therapeutic-level editing was achieved in NHP1 at ZFN:GFP = 1:3, a ratio at which efficacy was diminishing in the rat model, suggesting that the optimal ratio may differ between species. Clinical dose-finding studies will therefore be required to independently establish the optimal ZFN:donor ratio for human application, using the rat- and NHP-derived data as an initial reference.

In the broader context of RHO-adRP therapeutic development, the present approach offers a mechanistic advantage over EDIT-103 and OPGx-RHO[13,14], which deliver exogenous RHO supplementation from AAV-encoded expression cassettes. Because supplementation-based approaches express rhodopsin from exogenous promoters, the level of RHO expression is proportional to vector dose and transduction efficiency, creating a risk of overexpression toxicity at high vector doses[15] or haploinsufficiency under suboptimal transduction[9]. By inserting the therapeutic donor under the transcriptional control of the endogenous RHO promoter via HITI, the present approach subjects rhodopsin expression to the native regulatory architecture of the RHO locus, an approach predicted to avoid the dose-dependency of exogenous supplementation and to provide a more physiologically regulated expression profile. In the NHP experiment, HITI-mediated GFP expression was restricted to NRL-positive rod photoreceptors and absent in CRX-positive/NRL-negative cones, confirming cell-type-specific expression under endogenous RHO promoter control (Fig. 4D). These features, including mutation-agnostic design, endogenous promoter-driven expression, and freedom-to-operate as an IP-independent platform, support the potential of this therapeutic strategy for the broad RHO-adRP patient population[10,19].

This study establishes a preclinical proof-of-concept for ZF-ND1-mediated HITI as a mutation-agnostic therapeutic strategy for RHO-adRP. Mechanistic consistency across human-derived cells, a humanized rat disease model, and the primate retina provides cross-species validation of the editing mechanism. Dose-dependent structural preservation over 6 months in the rat model, with a well-defined optimal ZFN:donor ratio and consistent PoM-PoC correlation, supports rational dose selection for clinical development. In the primate retina, therapeutic-level photoreceptor correction was achieved under the conservative cleavage conditions imposed by target sequence mismatches, supporting the expectation that clinically relevant editing levels are attainable in patients with the fully matched human RHO sequence. The principal limitations of this dataset include the small NHP cohort size, the 3-month observation period in NHP, and the descriptive nature of the safety assessment; formal GLP toxicology studies and comprehensive off-target profiling are to be conducted prior to clinical translation. These data provide a foundation for advancing ZF-ND1-mediated HITI toward first-in-human investigation for RHO-adRP.

## Methods

### Plasmids and AAV Vectors

The details of all DNA constructs used in this study are listed in Supplementary Table S1. ZF pair target sequences and amino acid sequences of ZF modules are provided in Supplementary Tables S2 and S3. Zinc finger (ZF) pairs targeting the human *RHO* 5′-UTR were designed using the Scripps Research Institute ZF Tools algorithm[30] (http://www.zincfingertools.org). ZF module numbers correspond to those in the Zinc Finger Consortium modular assembly set[61] (Addgene Kit #1000000005). The target region comprised the ∼170 bp sequence proximal to the *RHO* translation start codon within the 5′-UTR. Candidate pairs with either 5-bp or 6-bp spacers between the two half-sites were generated, yielding 34 ZF pair combinations. 21 ZF pairs targeting the mouse *Rho* 5′-UTR were designed in a similar manner. Each ZF array was fused to either the ND1-DDD or ND1-RRR heterodimeric nuclease domain of FirmCut via a TGGS linker, generating paired ZF-ND1 expression plasmids (pCMV-XXXX-S/AS-ND1DDD/RRR).

The final optimized AAV5-ZFN vector (insert size 4.73 kb) incorporates both ZFN subunits in opposing orientations from a central bidirectional EF-1α/CMV promoter: [synthetic pA]–[ND1-RRR]–[5562AS-C6]–[EF-1α]←→[CMV]–[5461S]–[ND1-DDD]–[bGHpA]. The 5′ ITR incorporates an 11-bp deletion (5′-AAAGCCCGGGC-3′) to facilitate stable plasmid propagation and single-stranded AAV packaging. AAV5-RHO (insert size 2.26 kb) and AAV5-GFP (insert size 1.97 kb) were packaged as self-complementary AAV (scAAV), with insert sizes below the scAAV packaging threshold of approximately 2.3 kb (including the Δ-ITR). The donor cassette architecture is: [Δ-ITR]–[ZFN target]–[Chimeric intron]–[hRHO CDS or AcGFP]–[hGH polyA]–[ZFN target]–[ITR].

All AAV vectors (AAV5-ZFN, AAV5-RHO, and AAV5-GFP) were produced by Synprogen Co., Ltd. (Kobe, Japan) by triple-plasmid transfection of HEK293T cells followed by cesium chloride gradient ultracentrifugation (one to two rounds). Vector genomic copies were determined by qPCR.

### Animals

All animal experiments were conducted in accordance with the guidelines of the Institutional Animal Care and Use Committee of RIKEN Kobe Campus (approval numbers: A2022-03-11 and A2001-03). Wild-type CD1 mice were used for *in vivo* electroporation experiments. A humanized *RHO* rat model was generated by CRISPR/Cas9-mediated knock-in[62,63] of a 163 bp cassette comprising the human *RHO* 5′-UTR sequence (containing the ZF-ND1 target site) and a FLAG-tagged T17M point mutation (p.Thr17Met) into the endogenous rat *Rho* locus at exon 1 (Supplementary Fig. S6). Two guide RNAs flanking the insertion site (T1: 5′-TCTGTCTACGAACAGCCCGT<u>GGG</u>-3′; T2: 5′-CTTCTCCAACATCACGGGCG<u>TGG</u>- 3′; PAM sequences underlined) were used to excise the endogenous exon 1 sequence and replace it with the humanized donor cassette by homology-directed repair. Wistar rats (Albino) were used throughout. Heterozygous *Rho*^+/hRhoWT^ (carrying the human 5′-UTR without T17M; Accession No. CDB0102R: https://large.riken.jp/distribution/mutant-list.html) and *Rho*^+/hRhoT17M^ (carrying the human 5′-UTR with T17M; Accession No. CDB0101RE) animals were generated by crossing founder animals with wild-type Wistar rats. Genotyping was performed by PCR of tail-tip genomic DNA using primers 5′-TCAGTGCCTGGAGTTGTGCTGTG-3′ (forward) and 5′-CCAGGTAGTACTGCGGCTGCTCAAAG-3′ (reverse), yielding a 200 bp product from the wild-type allele and a 284 bp product from the knock-in allele, followed by Sanger sequencing for confirmation. Proof-of-mechanism studies were conducted in two crab-eating monkeys (*Macaca fascicularis*), designated NHP1 (tattoo ID 7881940874; male; 3 years 4 months at study entry) and NHP2 (tattoo ID 8274723288; male; 3 years 1 month at study entry), supplied by Eve Bioscience Co., Ltd. (Wakayama, Japan).

### Genomic DNA and cDNA preparation

Genomic DNA from cultured cells, hiPSC-derived retinal organoids, and NHP retinal tissue was extracted using the Puregene Tissue Kit (QIAGEN, cat# 158063). NHP retinal cDNA was synthesized using SuperScript III Reverse Transcriptase (Thermo Fisher Scientific, cat# 18080400) with random hexamer primers from the total retinal RNAs extracted using the RNeasy Mini Kit (QIAGEN, cat# 74104). All procedures were performed according to the manufacturer’s recommended protocols.

### Single-Strand Annealing (SSA) Assay

Cleavage activity of ZF-ND1 pairs was assessed by single-strand annealing (SSA) assay as previously described[23]. Briefly, HEK293T cells (RIKEN BioResource Research Center, Tsukuba, Japan) were seeded in 24-well plates and co-transfected using FuGENE6 (Roche) with 300 ng pCAG-EGxxFP SSA reporter carrying the human RHO ZF target sequence between two split EGFP fragments in the same orientation, 100 ng each of ZF-ND1-DDD and ZF-ND1-RRR expression plasmids, and 100 ng pCAG-mCherry as a transfection control. . Targeted double-strand cleavage triggers SSA-mediated EGFP reconstitution. Fluorescence images were acquired 48 hours post-transfection by fluorescence microscopy, and GFP reconstitution efficiency was quantified as the EGFP:mCherry fluorescence intensity ratio using ImageJ. Within each experimental run, ratios were normalized to the designated reference condition, set to 1.00. SpCas9 with a mouse Rho-targeting gRNA and the corresponding pCAG-EGxxFP reporter (Supplementary Table S1) was included as a positive control in SSA screening. For NHP sequence compatibility assessment, the 5461S recognition sequence was aligned with the corresponding NHP *RHO* 5′-UTR region (Supplementary Fig. S3A), and SSA assays comparing cleavage of the human, T-to-A mutant, and NHP *RHO* target sequences were performed in HEK293T cells using identical ZF-ND1 expression plasmids. GFP reconstitution efficiency was expressed as a percentage relative to the human target signal.

### Cel-1 (Surveyor) Nuclease Assay and TIDE Analysis

On-target cleavage activity was confirmed by Cel-1 (Surveyor) nuclease assay using the GeneArt Genomic Cleavage Detection Kit (Invitrogen, A24372) with the manufacturer-provided procedure. Briefly, HEK293T cells transfected with ZF-ND1 expression plasmids were harvested 3 days post-transfection, and a region flanking the ZF-ND1 target site was PCR-amplified using PrimeSTAR Max DNA Polymerase (TaKaRa, R045A) with forward primer 5′-ATCGATAAGCTTGATTATGAACACCCCCAATCTCCCAGATG-3′ and reverse primer 5′-AGAGCGTGAGGAAGTTGATGGGGAAGC-3′ (amplicon 378 bp; thermocycling: 94°C 2 min; 35 cycles of 98°C 10 s, 55°C 5 s, 72°C 10 s). PCR products were denatured and slowly re-annealed (95°C 5 min; 95→85°C at −2°C/s; 85→25°C at −0.1°C/s), followed by Detection Enzyme digestion (37°C, 1 h). Cleavage band intensity was assessed qualitatively by visual inspection; quantitative indel frequency was determined exclusively by TIDE analysis as described below.

For TIDE analysis, the same PCR product was submitted for Sanger sequencing and analyzed using the TIDE web tool[64] (https://apps.datacurators.nl/tide/) with the following parameters: guide sequence 5′-CAGGCCTTCGCAGCATTCTT-3′; alignment window 60 bp; decomposition window 115–685 bp; indel size range ±10 bp; P-value threshold 0.001. Background indel frequency was determined from the electropherogram of untransfected wild-type genomic DNA. TIDE analysis was performed on two independent transfection experiments for each variant; values shown represent one determination per experiment.

### *In Vivo* Electroporation in Neonatal Mouse Retina (Supplementary)

For NLS optimization in post-mitotic rod photoreceptors, four NLS variants were constructed using the mouse *Rho*-targeting 5057S/5057AS ZF-ND1 pair: 1×SV40 NLS (PKKKRKV), 3×SV40 NLS, 3×multiNLS (SV40–AAA–cMyc–GSG–SV40; cMyc NLS: PAAKRVKLD), and 5×multiNLS (Supplementary Table S1). Each variant was introduced into P0 wild-type mouse retina by *in vivo* electroporation as previously described[23,65], with a mouse *Rho* 5′-UTR-targeting HITI donor vector carrying AcGFP. Retinas were harvested at P21 for section immunohistochemistry and quantification of AcGFP-positive rod photoreceptors (Supplementary Fig. S2).

### hiPSC-Derived Retinal Organoid Preparation and AAV Transduction

The experiments were conducted in accordance with the Kobe City Eye Hospital Research Ethics Committee guidelines (approval number: Ki-O21-03). Human iPSC-derived retinal organoids were prepared as previously described[66] and used at differentiation day (DD) 150–200. For polyA signal optimization, organoids were transduced with AAV5-ZFN (0.5 × 10¹² GC/mL) plus either AAV5-GFP carrying hGH polyA or hRHO 3′-UTR (1.0 × 10¹² GC/mL each), or with AAV5-CMV-AcGFP-hGHpA (1.0 × 10¹² GC/mL) as an internal transduction efficiency control; organoids were assessed by fluorescence microscopy at DD150+30 (Fig. 2D-E). For ZFN:donor ratio optimization, AAV5-ZFN was held constant at 0.5 × 10¹² GC/mL while AAV5-GFP was titrated at 0.5, 1.0, 1.5, and 2.0 × 10¹² GC/mL (ZFN:GFP = 1:1, 1:2, 1:3, and 1:4), with AAV5-GFP alone (1.0 × 10¹² GC/mL, no ZFN) as a negative control; organoids were incubated for 30 days post-transduction (Fig. 2F). Complementary stoichiometric titration was performed in HEK293T cells by co-transfection of two plasmids for AAV5-ZFN and AAV5-RHO at ratios of 1:1; genomic DNA was extracted 48 hours post-transfection.

### Subretinal Injection

Subretinal injection of AAVs for rats was performed as previously described[67] with minor modifications. Therapeutic (AAV5-ZFN + AAV5-RHO) or PoM verification vector (AAV5-ZFN + AAV5-GFP) was administered at approximately 1 month of age. Rats were anesthetized with medetomidine (0.375 mg/kg), midazolam (2 mg/kg), and butorphanol (2.5 mg/kg) by intraperitoneal injection. Glass injection pipettes were prepared using a micropipette puller (Sutter P-97) and microgrinder (Narishige EG-400). AAV solution (2 µL per eye) was injected through a trans-scleral approach under surgical microscope visualization using a Micro4 microinjector (WPI). Atipamezole (0.75 mg/kg i.p.) was administered at the end of the procedure to reverse anesthesia. Group assignments: ZFN:RHO 1:1 (AAV5-ZFN 0.5 × 10¹² + AAV5-RHO 0.5 × 10¹² GC/mL), n = 4; 1:2 (0.5 + 1.0 × 10¹²), n = 4; 1:3 (0.5 + 1.5 × 10¹²), n = 4; 1:4 (0.4 + 1.6 × 10¹²), n = 4; Control (AAV5-CMV-EGFP + AAV5-RHO, no ZFN), n = 6. For long-term safety assessment, wild-type Wistar rats received AAV5-ZFN + AAV5-RHO (ZFN:RHO = 1:2) at standard dose (AAV5-ZFN 0.5 × 10¹² + AAV5-RHO 1.0 × 10¹² GC/mL; n = 1) or one-third dilution (total 0.5 × 10¹² GC/mL; n = 1).

For crab-eating monkey injections, animals were anesthetized with ketamine (15 mg/kg i.m.) and xylazine (1 mg/kg i.m.) for immobilization and maintained on isoflurane inhalation anesthesia via endotracheal intubation. Subretinal injection was performed without prior vitrectomy (bleb-free direct subretinal delivery) using a 25G/38G Nano Cannula (MedOne, cat# 3263). The injection needle was inserted through a trans-scleral approach and advanced to the paramacular region. AAV solution was delivered at a slow, steady rate; the full syringe volume was pushed to ensure accurate delivery. A target volume of 100 µL per eye was used. Fast Green (0.05%) was included as a visual indicator of subretinal bleb formation. Vitrectomy-assisted delivery (Hamilton syringe with PolyVent cannula, MedOne cat# 3247) was available as an alternative in the event of incomplete delivery. Right-eye injection was performed first for each animal. To attenuate immune responses to AAV5 vector, animals received prednisolone by intramuscular injection at 1 mg/kg/day from Day −1 through Day 2 post-injection, followed by tapering to 0.5 mg/kg/day from Day 3 through Day 10 (with rest days on Day 7 and Day 9). Triamcinolone acetonide (sub-Tenon injection, STTA; 0.5 mL/eye) was administered on the day of injection and at post-injection Days 14, 28, and 56. Note: pre-operative screening for AAV5 serum neutralizing antibodies was not performed in this study; the potential contribution of pre-existing immunity to the inflammatory findings observed in NHP2 cannot be excluded.

### Section Immunohistochemistry and Retinal Flatmounts

Retinal eyecups were fixed with 4% paraformaldehyde in PBS for 1 hour or overnight. For cryosections, fixed tissue was cryoprotected in 30% sucrose/PBS overnight, embedded in OCT compound, and sectioned at 14∼20 µm. Sections and flatmounts were blocked for 1 hour at room temperature with 5% heat-inactivated horse serum (Thermo Fisher Scientific, cat# 26050070) and 0.5% Triton X-100 in PBS.

For knock-in verification (Fig. 3C), rat retinal sections were immunostained with anti-GFP (rat monoclonal, Nacalai Tesque cat# 04404-26, 1:1000) and anti-FLAG (mouse monoclonal, Millipore cat# F1804, 1:1000). For 6-month harvest analysis (Fig. 3F), sections were immunostained with anti-GFP (rat monoclonal, Nacalai Tesque cat# 04404-84, 1:1000) and anti-RHO (rabbit polyclonal, Abcam cat# ab112576, 1:2000). For NHP retinal sections (Fig. 4D), cryosections were immunostained with anti-GFP (rat monoclonal, Nacalai Tesque cat# 04404-26, 1:1000), anti-NRL (goat polyclonal, R&D Systems cat# AF2945, 1:1000), and anti-CRX (mouse monoclonal, Abnova cat# H00001406-M02, 1:20000). Secondary antibodies were Alexa Fluor 488/594/647 donkey anti-mouse/rabbit/rat/goat (Thermo Fisher Scientific) in 1:1000 dilution. Images were acquired on a BZ-X810 (Keyence) fluorescence microscope; selected sections were additionally imaged on an LSM980 (Zeiss) confocal microscope.

GFP-positive rod photoreceptor fraction was quantified from immunostained NHP sections sampled at 400 µm intervals across the GFP-positive area. In each section, GFP⁺/NRL⁺ cells (HITI-edited rod photoreceptors) and NRL⁺ cells (total rod photoreceptors) were counted. GFP⁺/NRL⁺ fraction (%) = [GFP⁺/NRL⁺ cells] / [NRL⁺ cells] × 100. Each data point in the spatial heatmap corresponds to the GFP⁺/NRL⁺ fraction from one section mapped onto the corresponding retinal flatmount image.

### Optical Coherence Tomography (OCT) Imaging

Retinal structure in rats was monitored longitudinally by OCT using the Envisu 2210 (Leica Microsystems). Animals were anesthetized with isoflurane and pupils dilated with topical tropicamide. ONL thickness was measured manually from OCT B-scans at 7 spatially distributed locations within the inferior retinal quadrant. For natural history characterization, *Rho*^+/+^ (wild-type), *Rho*^+/hRhoWT^, and *Rho*^+/hRhoT17M^ animals (n = 3–4 per genotype) were assessed at postnatal days 27, 49, 58, 81, 87, 115, and 172. For longitudinal efficacy, OCT imaging was performed monthly for 6 months following injection. A threshold of 90 µm ONL thickness was defined as the lower limit of photoreceptor preservation. All measurements were performed by a single observer. For crab-eating monkeys, OCT imaging was performed at pre-injection baseline and at post-injection weeks 1, 2 and months 1, 2, and 3, using the RS-3000 OCT system (NIDEC) under isoflurane anesthesia with topical pupil dilation. Retinal layer integrity was assessed qualitatively; ONL thickness was not used as a primary efficacy endpoint given differences in layer definition between NHP and human OCT.

### Fundus Photography and Fluorescein Angiography (Non-Human Primates)

Fundus photography and fluorescein angiography (FA) were performed at pre-injection baseline and at post-injection weeks 1, 2 and months 1, 2, and 3, using a Canon CX-01 fundus camera under isoflurane anesthesia. Successful subretinal delivery was confirmed by the presence of a visible injection bleb on fundus imaging. Inflammatory changes were assessed by FA; active intraocular inflammation was defined by hyperfluorescence in the injection area on late-phase FA images.

### Droplet Digital PCR (ddPCR)

HITI-mediated knock-in allele frequency was quantified by droplet digital PCR (ddPCR) using the QX200 system (Bio-Rad). Genomic DNA was digested with BamHI (4 units per 20 µL reaction; NEB, cat# R0136S) prior to droplet partitioning to improve template accessibility. Template DNA was added at 100 ng per reaction. Each 20 µL ddPCR reaction contained 1× ddPCR Supermix for Probes (no dUTP) (Bio-Rad, cat# 1863023), 1× Primer & Probe mix for Rho-HITI (FAM channel; forward primer 5′-TCAGAACCCAGAGTCATCCAG-3′, reverse primer 5′-TCGACAAGCCCAGTTTCTATTGG-3′, probe 5′-[FAM]- CCAGTGCCTCACGACCAACTTCT-[BHQ1]-3′, antisense), and 1× Primer & Probe mix for *GAPDH* (HEX channel; forward primer 5′-tccgggtctttgcagtcgtatg-3′, reverse primer 5′-agtagggacctcctgtttctgg-3′, probe 5′-[HEX]-AAGGAGAGCTCAAGGTCAGCGC-[BHQ1]-3′). Primer and probe mixes were obtained from FASMAC Co., Ltd. (Kanagawa, Japan).

Droplets were generated using a DG8 cartridge (Bio-Rad, cat# 1864008) with 70 µL of Droplet Generator Oil for Probes (Bio-Rad, cat# 1863005) and transferred to ddPCR 96-well plates (Bio-Rad, cat# 12001925), sealed using the PX1 PCR Plate Sealer (180°C, 5 seconds). Thermal cycling: 95°C 10 min; 40 cycles of 94°C 35 sec and 62°C 65 sec; 98°C 10 min; hold at 12°C; ramp rate 2°C/sec throughout. Droplets were read using the QX200 Droplet Reader and analyzed with QX Manager software. FAM-positive droplets were assigned to the KI-positive population and HEX-positive droplets to the *GAPDH*-positive population; amplitude thresholds were determined automatically. KI allele frequency was expressed as copies of KI allele per copies of *GAPDH* (copies/µL, by Poisson statistics).

### KI Junction Sequence and cDNA Sequencing

To characterize the genomic sequence at HITI-mediated knock-in junctions, genomic DNA from AAV5-transduced hiPSC-derived retinal organoids (ZFN:GFP = 1:2, 1:3, and 1:4), MeWo human melanoma cells (obtained from the American Type Culture Collection, ATCC, Manassas, VA, USA; ZFN:GFP = 1:1; included as a non-photoreceptor cell line to confirm ZFN activity in an independent cellular context), and NHP retinal tissue from NHP1 (ZFN:GFP = 1:3) was PCR-amplified using KI junction-spanning primers and subjected to Sanger sequencing (Supplementary Fig. S5). The predicted targeted allele sequence was derived from in silico analysis of ZF-ND1 cleavage at the *RHO* 5′-UTR target site followed by NHEJ-mediated ligation of the donor cassette terminus.

To confirm active transcription of the HITI-integrated cassette in NHP retina, retinal tissue from the injection area of NHP1 was used for RNA extraction and RT-PCR using primers flanking the HITI junction spanning the chimeric intron–*hRHO* cDNA boundary. PCR products were subjected to Sanger sequencing (Supplementary Fig. S10).

### Statistical Analysis

For natural history comparison, retinal thickness at each time point was compared between *Rho*^+/+^ and *Rho*^+/hRhoWT^ using per-animal medians (across 7 OCT measurement locations) by Wilcoxon rank-sum test (n = 3–4 animals per group).

For longitudinal efficacy analysis, a Bayesian linear mixed model (LMM) was fitted using PyMC (version 5.28.3)[68] and ArviZ (version 0.23.4)[69]. The model was specified as: *thickness ∼ group × month + (1|animal)*, where group is a categorical predictor for treatment assignment, month is the continuous time variable, and (1|animal) denotes a random intercept for each animal capturing between-animal variability. Weakly informative priors: μ_group ∼ Normal(120, 50); β_time ∼ Normal(0, 20); β_group×month ∼ Normal(0, 10); σ_animal, σ_obs ∼ HalfNormal(20). NUTS sampler; 4 chains × 2,000 iterations (1,000 warm-up); target acceptance 0.90; random seed 42. Convergence: R-hat ≤ 1.002, ESS > 2,700 for all parameters. Treatment effects are reported as posterior medians with 95% credible intervals (CrI) at month 6. All analyses were performed in Python 3.12 using NumPy 2.4.2, PyMC 5.28.3, and ArviZ 0.23.4. Statistical analyses for NHP data were not performed given the descriptive and exploratory nature of the proof-of-mechanism study (n = 2).

## Acknowledgements

We received generous support from all members of the Vision Care Group, Kobe City Eye Hospital Research Center and Cell and Gene Therapy in Ophthalmology Laboratory, RIKEN DMP and BZP. We thank the members of the Laboratory for Animal Resources and Genetic Engineering (LARGE), RIKEN Center for Biosystems Dynamics Research, for generating and maintaining the rats; Satoshi Shirae and Toshika Senba at Vision Care Inc. for their professional support in NHP experiments, maintenance and data management; Shunsuke Saito and Yoshitomo Takagi at Synplogen Co., Ltd. for technical support for AAV vectors.

This work was supported by Santen Pharmaceutical Co., Ltd. (Osaka, Japan); Kobe University Capital Inc. (Kobe, Japan); Ritsumeikan University Capital Inc. (Kyoto, Japan); and the RIKEN Program for Drug Discovery and Medical Technology Platforms (DMP). This research was also partially supported by the Research Support Project for Life Science and Drug Discovery (Basis for Supporting Innovative Drug Discovery and Life Science Research (BINDS)) from the Japan Agency for Medical Research and Development (AMED) under Grant Number JP26ama121015.

## Author Contributions

Author contributions are defined according to the CRediT (Contributor Roles Taxonomy) framework.

Akishi Onishi: Conceptualization, Methodology, Investigation, Validation, Resources, Software, Formal analysis, Data Curation, Writing – Original Draft, Writing – Review & Editing, Visualization, Supervision, Project administration, Funding acquisition.

Tetsushi Sakuma: Conceptualization, Methodology, Investigation, Validation, Resources, Formal analysis, Writing – Review & Editing.

Michiko Mandai: Methodology, Investigation, Validation, Resources, Writing – Review & Editing.

Takashi Watanabe: Methodology, Investigation, Validation, Resources, Formal analysis, Data Curation, Visualization.

Takaho Endo: Methodology, Investigation, Validation, Resources, Software, Formal analysis, Data Curation, Visualization.

Wataru Nomura: Methodology, Resources, Formal analysis, Writing – Review & Editing. Aiko Ishimaru: Investigation, Validation, Resources, Data Curation.

Ken-ichi Inoue: Methodology, Resources, Investigation, Validation. Junki Sho: Investigation, Validation, Resources, Data Curation.

Yoko Ohigashi: Investigation, Validation, Resources, Data Curation. Yuki Nakano: Investigation, Validation, Data Curation.

Kazushi Yasuda: Methodology, Investigation.

Sota Ozaki: Investigation, Validation.

Akiko Maeda: Validation, Data Curation, Writing – Review & Editing. Chikako Morinaga: Supervision, Project administration.

Tetsuo Itoh: Writing – Review & Editing, Project administration, Funding acquisition.

Yuji Inomata: Writing – Review & Editing, Project administration, Funding acquisition, Writing – Review & Editing.

Yukihide Momozawa: Conceptualization, Methodology, Formal analysis, Supervision, Writing – Review & Editing.

Takashi Yamamoto: Conceptualization, Methodology, Supervision, Project administration, Writing – Review & Editing.

Hiroshi Kiyonari: Methodology, Resources, Supervision, Project administration, Writing – Review & Editing.

Seiji Hori: Supervision, Project administration, Funding acquisition.

Masayo Takahashi: Conceptualization, Methodology, Supervision, Project administration, Funding acquisition, Writing – Review & Editing.

## Data Availability

The datasets supporting the current study have not been deposited in a public repository due to their large size but are available from the corresponding author upon reasonable request. Any additional information required to reanalyze the data is also available upon request.

## Supplementary Figure Legends

**Supplementary Figure S1. TIDE (Tracking of Indels by Decomposition) for 5461S + 5562AS Refined Variants (C1–C8).**

TIDE analysis of Sanger sequencing electropherograms from HEK293T cells transfected with 5461S paired with each 5562AS variant. Total TIDE efficiency (%). TIDE web tool parameters: guide sequence 5′-CAGGCCTTCGCAGCATTCTT-3′; alignment window 60 bp; decomposition window 115–685 bp; indel size range ±10 bp; P-value threshold 0.001. Background determined from untransfected wild-type genomic DNA. Two independent transfection experiments were performed for each variant; electropherograms from one representative experiment are shown.

**Supplementary Figure S2. Mouse Rho 5′-UTR ZF-ND1 Screening and NLS Optimization in Post-Mitotic Rod Photoreceptors.**

**(A)** Schematic illustration of ZF pairs used for SSA assay targeting the mouse Rho 5′-UTR.

**(B)** SSA fluorescence images of HEK293T cells co-transfected with pCAG-EGxxFP, ZF-ND1 plasmids, and pCAG-mCherry (mCherry channel not shown for clarity). Panels grouped by spacer length: 6-bp spacer (5057AS + 5057S, 5360AS + 5360S, 8289AS + 8289S) and 5-bp spacer (5360AS + 5259S, 8289AS + 8288S). Normalized EGFP:mCherry ratios (relative to SpCas9/mRho-gRNA1 positive control from Fig. 1B = 1.00) are shown in the lower-left corner of each panel. Scale bar = 500 µm.

**(C)** Protein domain structures for NLS optimization. Sense (ND1-DDD–5057S–NLS) and antisense (ND1-RRR–5057AS–NLS) subunits shown for 1×SV40 NLS (PKKKRKV), 3×SV40 NLS, 3×multiNLS (SV40–AAA–cMyc–GSG–SV40; cMyc NLS: PAAKRVKLD), and 5×multiNLS.

**(D)** *In vivo* NLS optimization by electroporation into neonatal mouse retina. Left: experimental scheme showing subretinal injection of plasmid cocktail into P0 wild-type mouse eyes followed by electroporation. The cocktail comprised bRho300bp-5057S, bRho300bp-5057AS, a mouse *Rho* HITI donor cassette (*Rho* cDNA–Furin-P2A–AcGFP), and pCAG-mCherry. bRho300bp is a 300-bp proximal promoter of bovine rhodopsin. Right: section IHC at P21; AcGFP (green; HITI insertion) and mCherry (red; electroporated cells) in ONL, INL, GCL. Scale bar = 50 µm.

**Supplementary Figure S3. Compatibility of ZF-ND1 with the Crab-Eating Monkey (*Macaca fascicularis*) RHO Target Sequence.**

**(A)** Sequence alignment of the ZF-ND1 target site in human (Hs) and NHP RHO 5′-UTR. Three nucleotide differences in the 5461S recognition region are indicated in lowercase. Protein domain structures of the final ZF-ND1 sense and antisense subunits are shown. The T-to-A mutation in the donor vector prevents inadvertent ATG translation initiation following HITI; the modified target sequence is used in AAV5-RHO and AAV5-GFP.

**(B)** SSA assay comparing ZF-ND1 cleavage at the unmodified human RHO target (Hs), the T-to-A mutant target (Donor), and the NHP RHO target. pCAG-mCherry included as a transfection efficiency control (mCherry channel not shown for clarity). Normalized EGFP:mCherry ratios (relative to Hs RHO target = 1.00) are shown in the lower-left corner of each panel. T-to-A mutation retained ∼94% of cleavage efficiency relative to the human target. Two independent SSA experiments for the NHP RHO target are shown, each yielding ∼30–40% lower EGFP reconstitution relative to the human target. Scale bar = 500 µm.

**Supplementary Figure S4. TIDE Electropherograms for Promoter Configuration Comparison.**

TIDE analysis of Sanger sequencing electropherograms from HEK293T cells transfected with each of the four promoter configurations. Total TIDE efficiency (%): Individual CMV+CMV, 25.7%; Bicistronic Furin-P2A, 4.6%; Individual EF-1α+CMV, 22.6%; Bidirectional EF-1α/CMV, 27.1%. Note: values shown here represent the second independent experiment; Fig. 2C reports values from the first experiment (26%, 5%, 23%, 27%), demonstrating reproducibility. TIDE parameters identical to those in Supplementary Fig. S1. Two independent experiments performed for each configuration; electropherograms from one representative experiment shown.

**Supplementary Figure S5. KI Junction Sanger Sequencing Across Cell Types and ZFN:Donor Ratios.**

KI junction sequences from MeWo human melanoma cells (ZFN:GFP = 1:1; included as a non-photoreceptor cell line to confirm ZFN activity in an independent cellular context); hiPSC-derived retinal organoids (ZFN:GFP = 1:2, 1:3, and 1:4); and NHP crab-eating monkey retinal tissue from NHP1 (see Fig. 4; ZFN:GFP = 1:3). The predicted targeted allele sequence following HITI-mediated integration (from in silico analysis of ZF-ND1 cleavage followed by NHEJ-mediated ligation of the donor cassette terminus) is shown for reference.

**Supplementary Figure S6. Generation and Characterization of the Humanized RHO-T17M Rat Model.**

Knock-in strategy. Guide RNAs T1 (5′-TCTGTCTACGAACAGCCCGT<u>GGG</u>-3′) and T2 (5′-CTTCTCCAACATCACGGGCG<u>TGG</u>-3′; PAM sequences underlined) directed excision and replacement of rat *Rho* exon 1 with the humanized cassette (human *RHO* 5′-UTR + FLAG-T17M) via HDR with 0.5 kb homology arms flanking the insertion site. Two lines were generated: *Rho*^+/hRhoWT^ (carrying the human 5′-UTR without T17M) and *Rho*^+/hRhoT17M^ (carrying the human 5′-UTR with T17M). Genotyping PCR primers (forward, black arrow: 5′-TCAGTGCCTGGAGTTGTGCTGTG-3′; reverse, red arrow: 5′-CCAGGTAGTACTGCGGCTGCTCAAAG-3′) yield a 200 bp product from the wild-type allele and a 284 bp product from the knock-in allele.

**Supplementary Figure S7. Dose Optimization in *Rho*^+/hRhoWT^ Rats: GFP Expression as a Function of AAV5-ZFN Concentration.**

*Rho*^+/hRhoWT^ rats (n = 3 per dose group) received PoM verification vector (AAV5-ZFN + AAV5-GFP, 1:1) at 0.05, 0.16, or 0.5 × 10¹² GC/mL at 1 month of age; harvested at 2 months. Representative flatmount images (superior and inferior quadrants) per dose shown; overlay of GFP (green) and FLAG (red) channels with GFP-only inset (superior quadrant). GFP signal increased with concentration. Scale bar = 1 mm.

**Supplementary Figure S8. Long-term Safety Assessment of Therapeutic Vector in Wild-Type Rats.**

Wild-type albino Wistar rats (n = 1 per dose group) received subretinal injection of AAV5-ZFN + AAV5-RHO (AAV5-ZFN + AAV5-RHO, ZFN:RHO = 1:2) at 1 month of age. Standard dose (No.28: AAV5-ZFN 0.5 × 10¹² + AAV5-RHO 1.0 × 10¹² GC/mL; blue) and one-third dilution (No.24: 0.167 + 0.333 × 10¹² GC/mL; green) are shown. Left: ONL thickness over 5 months post-injection. Lines: per-time-point medians of 7 OCT measurements; shaded area: full range of 7 spatially distributed measurements per time point (reflecting spatial heterogeneity of the retinal area surrounding the injection site); semi-transparent dots: individual measurements. Red dotted line: 90 µm threshold. Both animals maintained 173–211 µm throughout. Right: representative OCT B-scans from No.28 (standard dose) at months 2–5 showing preserved retinal laminar architecture.

**Supplementary Figure S9. Fundus and OCT Data for NHP2 (Reference, Inflammatory Cohort).**

**(A)** Fundus photographs, fluorescein angiograms (FA), and OCT images from NHP2 (Right and Left eyes) at pre-injection baseline, post-injection 2 weeks, and 3 months. FA-visible inflammatory changes are observed in the injection area at post-injection month 2 in both eyes.

**(B)** Retinal flatmount fluorescence images of NHP2 showing GFPL photoreceptors within the injection area at 3-month harvest. Total retinal area: right eye ∼78 mm², left eye ∼67 mm².

**(C)** Spatial heatmap of GFPL/NRLL photoreceptor fraction (%) overlaid on the retinal flatmount of NHP2. The eye with greater inflammatory changes showed reduced GFPL photoreceptor density on both flatmount imaging and heatmap quantification. These data are presented separately as a reference observation; NHP2 data are not included in the primary efficacy analysis. Note: pre-operative screening for AAV5 serum neutralizing antibodies was not performed; pre-existing immunity may have contributed to the inflammatory response.

**Supplementary Figure S10. cDNA Sequencing Confirms Chimeric Intron Splicing and Active Transcription of HITI-Integrated hRHO Donor.**

RT-PCR of retinal RNA from the injection area in NHP1. Upper: predicted spliced cDNA sequence spanning the chimeric intron–hRHO/GFP donor junction, shown for reference. Middle: Sanger sequencing electropherogram of the RT-PCR product confirming chimeric intron splicing at the predicted splice acceptor site and the expected KI junction sequence. Lower: sequencing trace with reverse primer orientation showing the hRHO/GFP donor–chimeric exon junction. A minor displacement of the deletion pattern from the predicted ZF-ND1 cleavage site (zipper-like trace shift) is observed, attributed to the three-nucleotide mismatch in the 5461S ZF recognition region in NHP, which alters ZF array binding and ND1 heterodimerization geometry, resulting in a slightly shifted NHEJ repair pattern. This alteration is expected to be absent in human patients where the fully matched *RHO* sequence is the substrate.

