## Supplementary material for "Mutation-agnostic gene insertion therapy for RHO-associated autosomal dominant retinitis pigmentosa using zinc finger nucleases": FigS1-S10, TableS1-S3

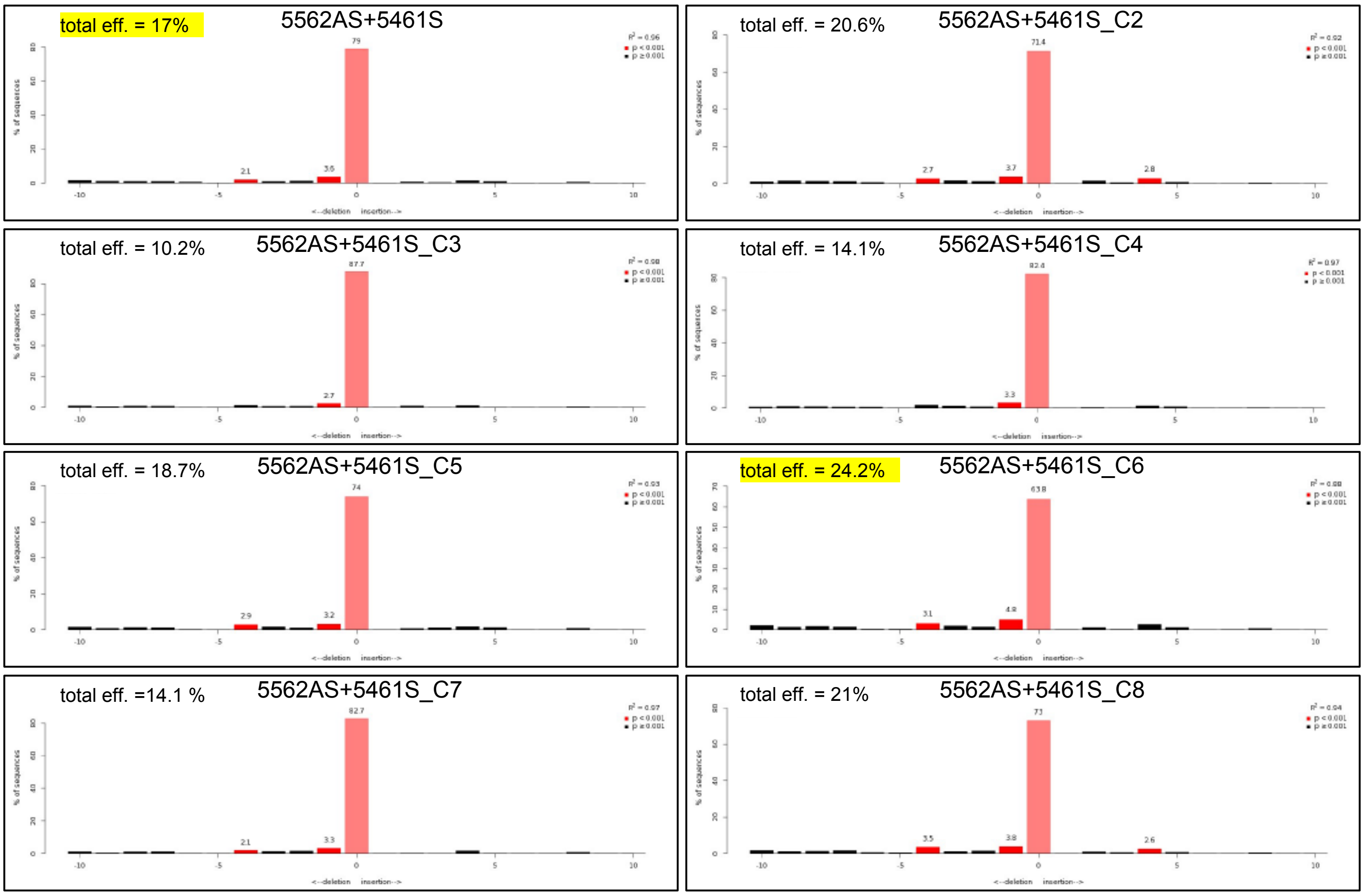


Figure S1. Onishi et al.

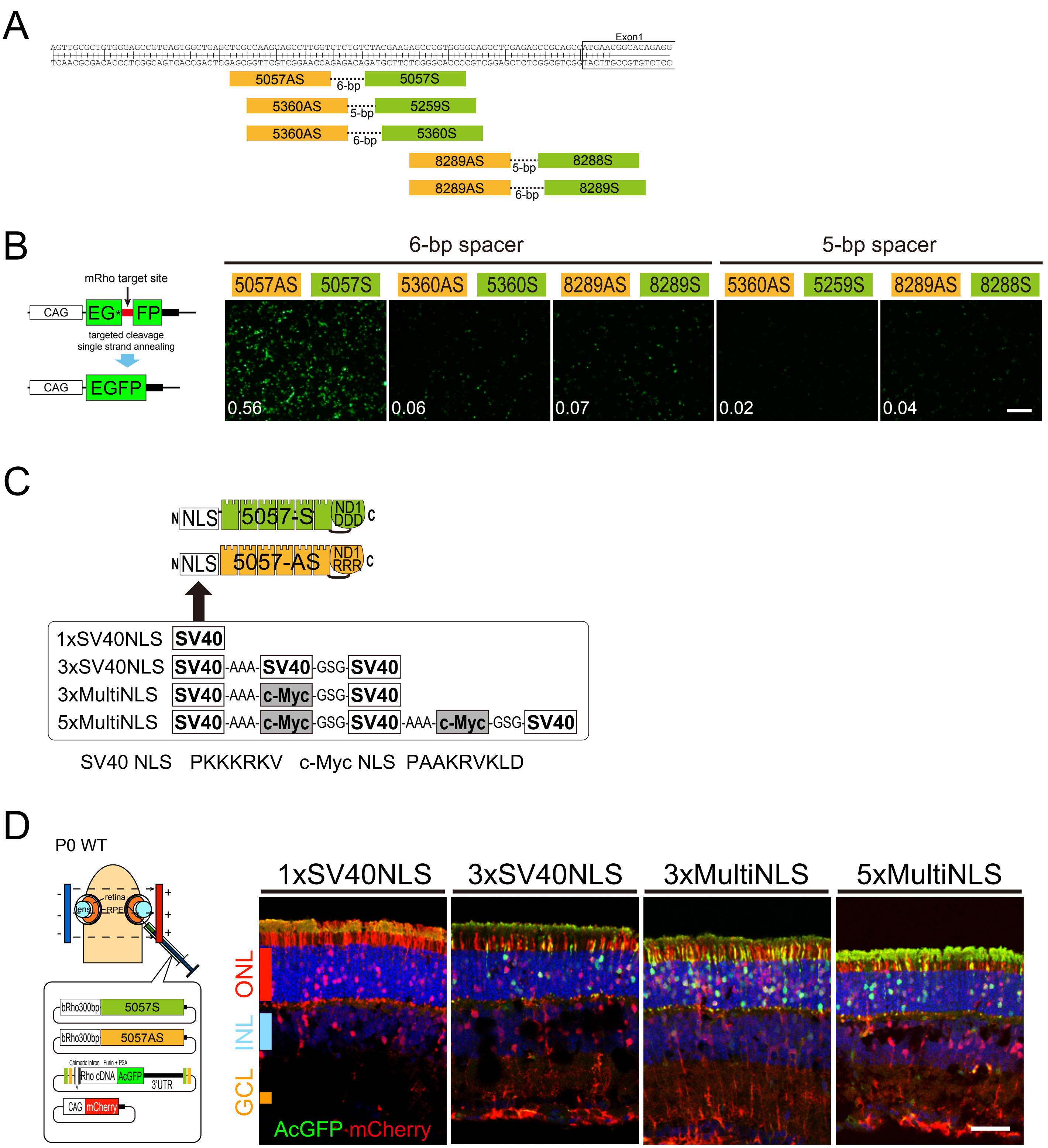


Figure S2. Onishi et al.

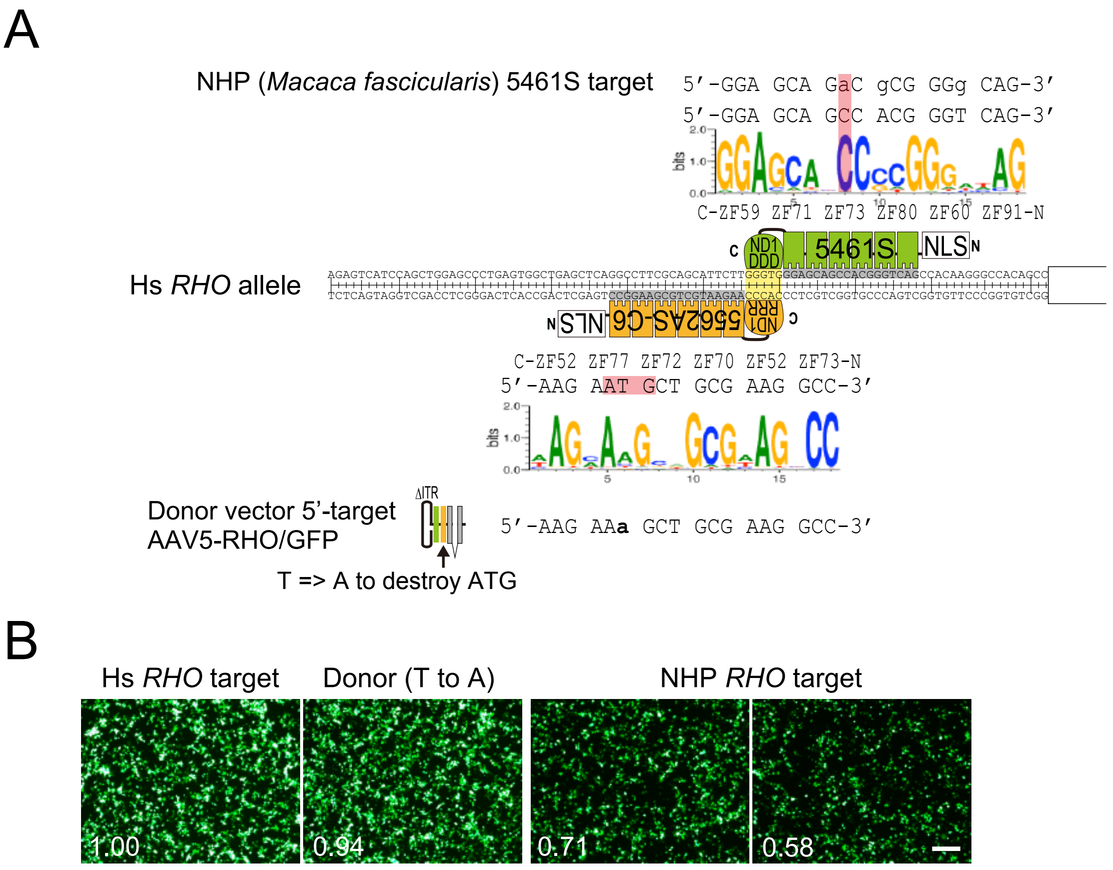


Figure S3. Onishi et al.

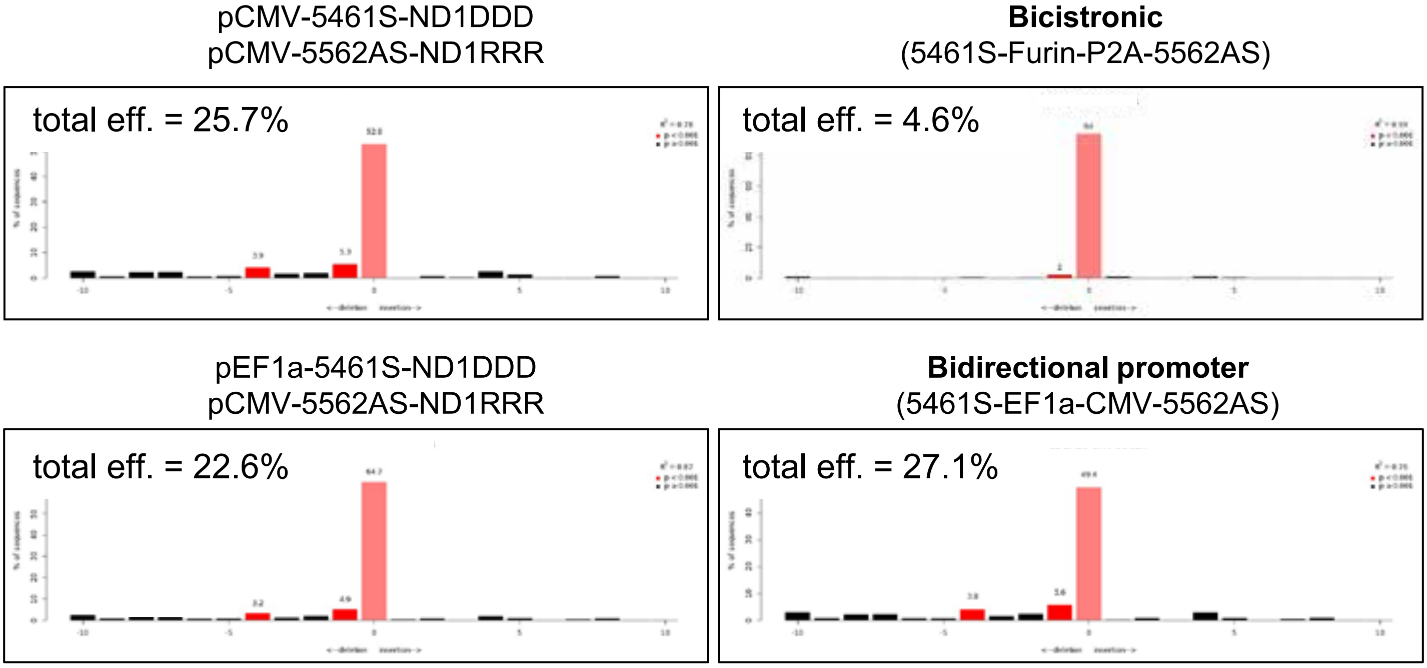


Figure S4. Onishi et al.

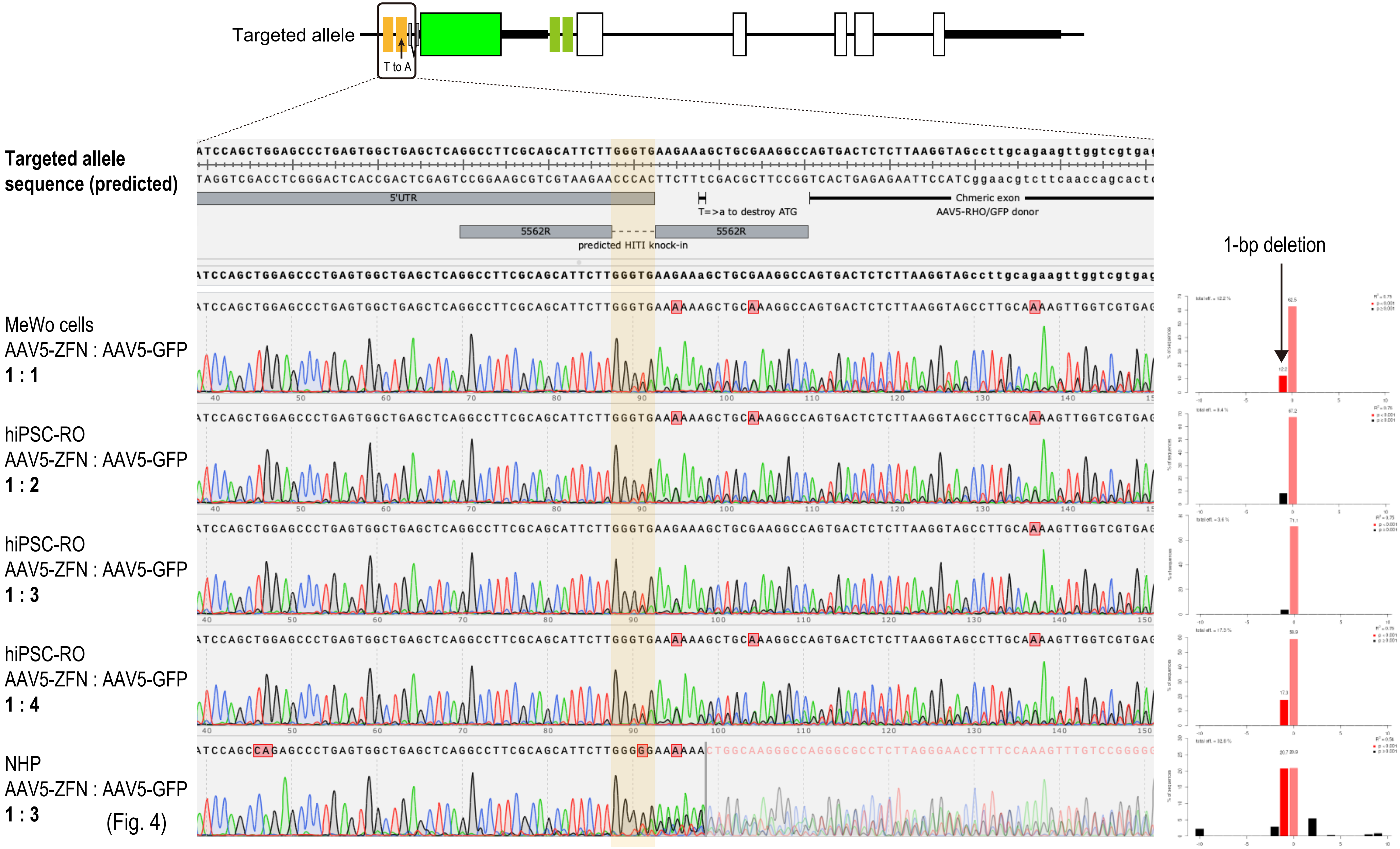


Figure S5. Onishi et al.

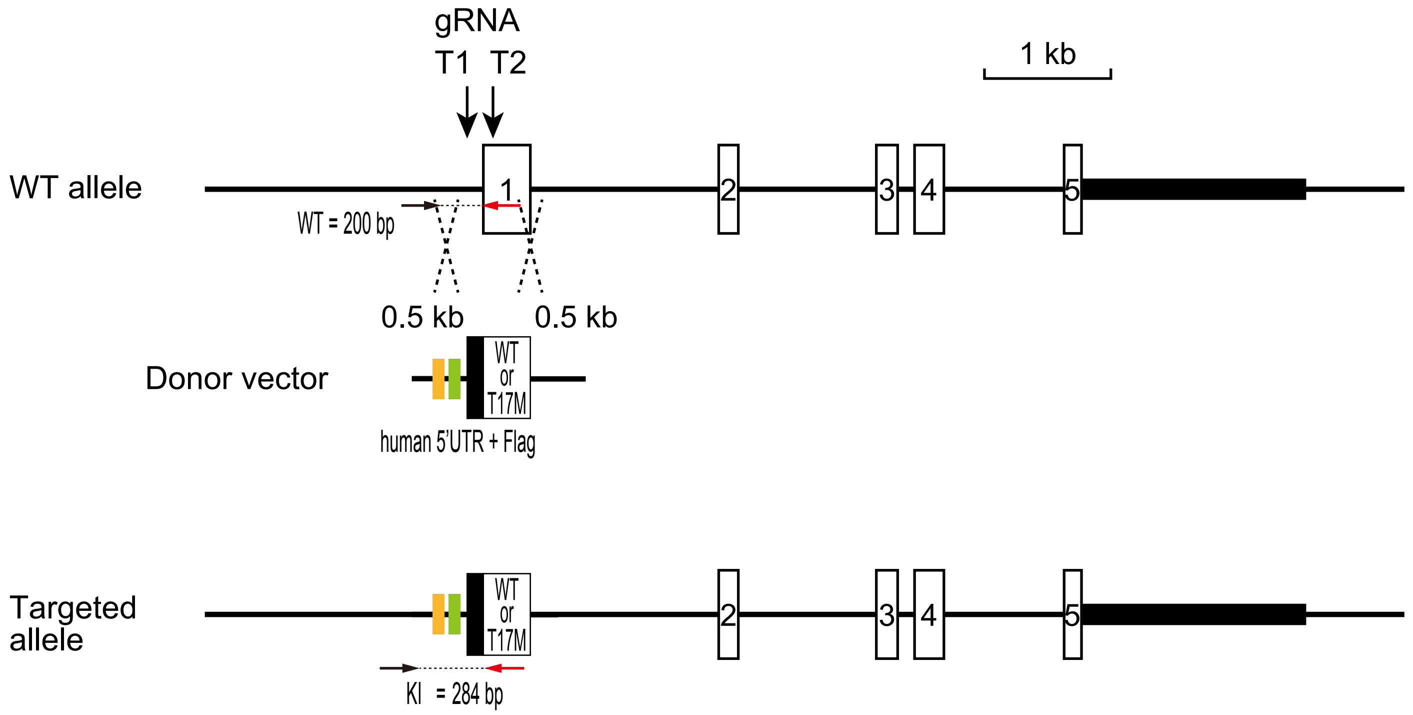


Figure S6. Onishi et al.

Rho<+/hWT>

1Mo injection; +2Mo harvest, flatmount IHC

| | | | | |
|---|---|---|---|---|
| AAV5-ZFN | 0.05 | 0.16 | 0.5 | |
| AAV5-GFP | 0.05 | 0.16 | 0.5 | x10$^{12}$GC/mL |

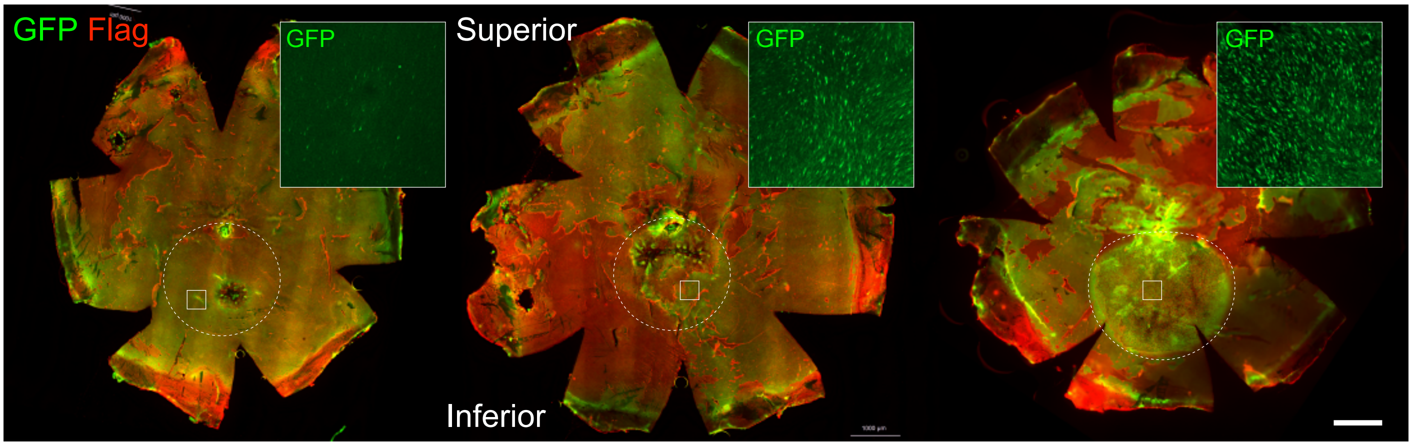


Figure S7. Onishi et al.

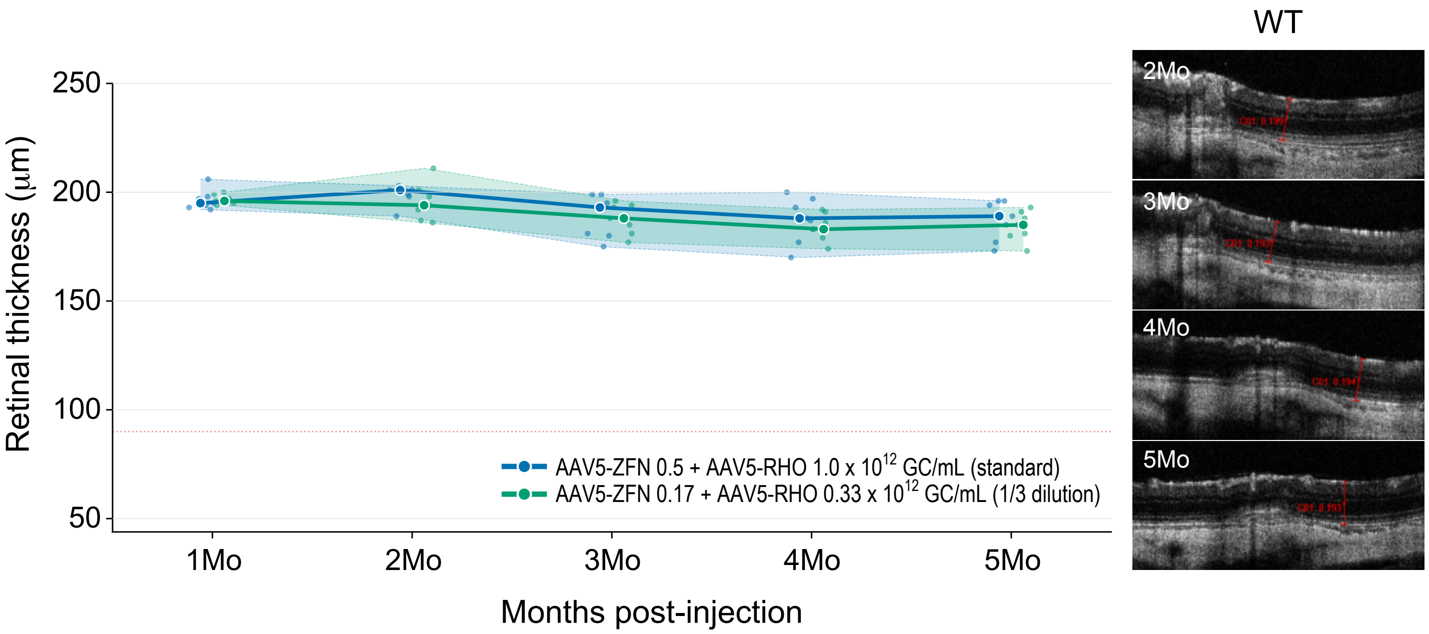


Figure S8. Onishi et al.

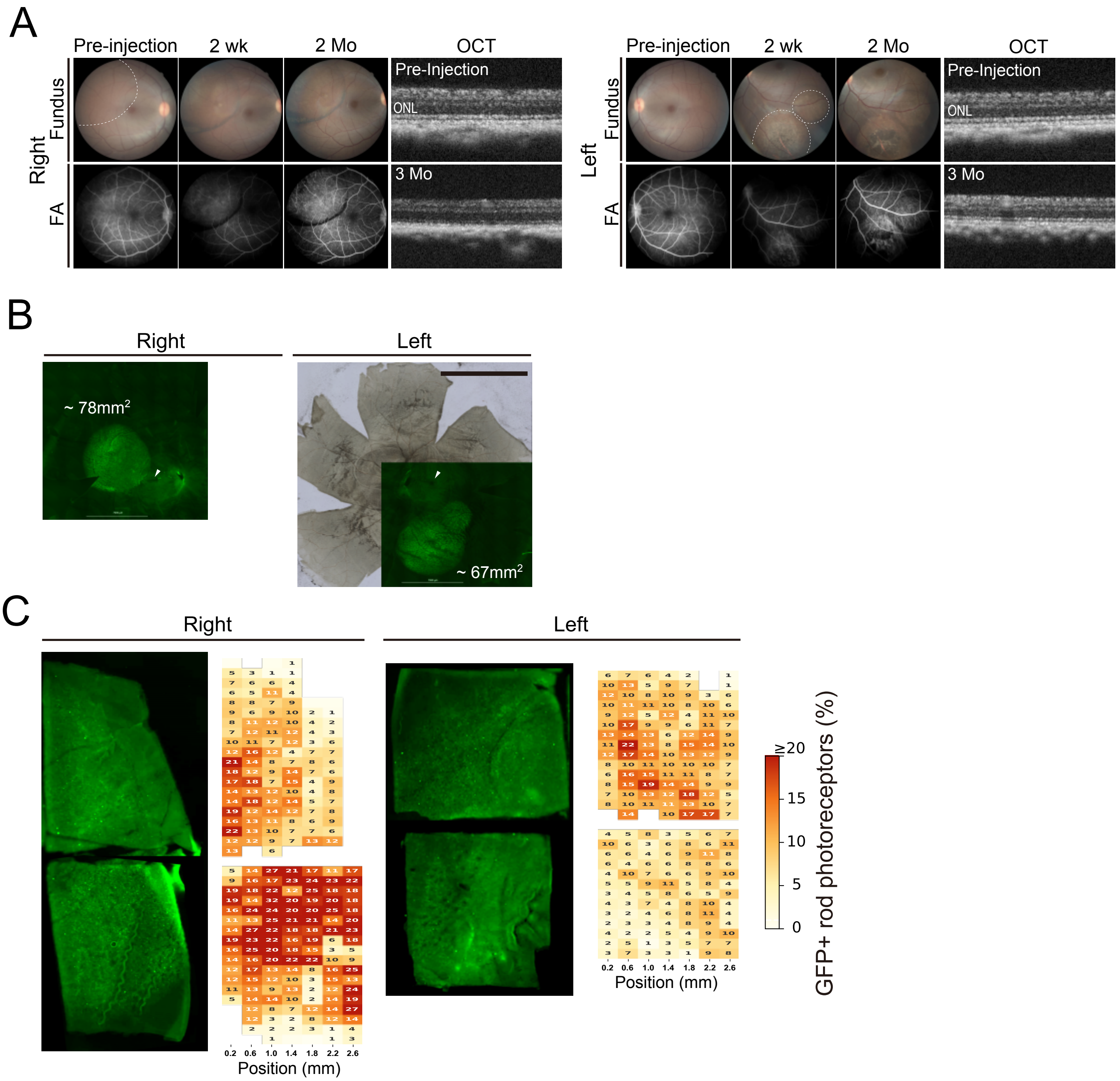


Figure S9. Onishi et al.

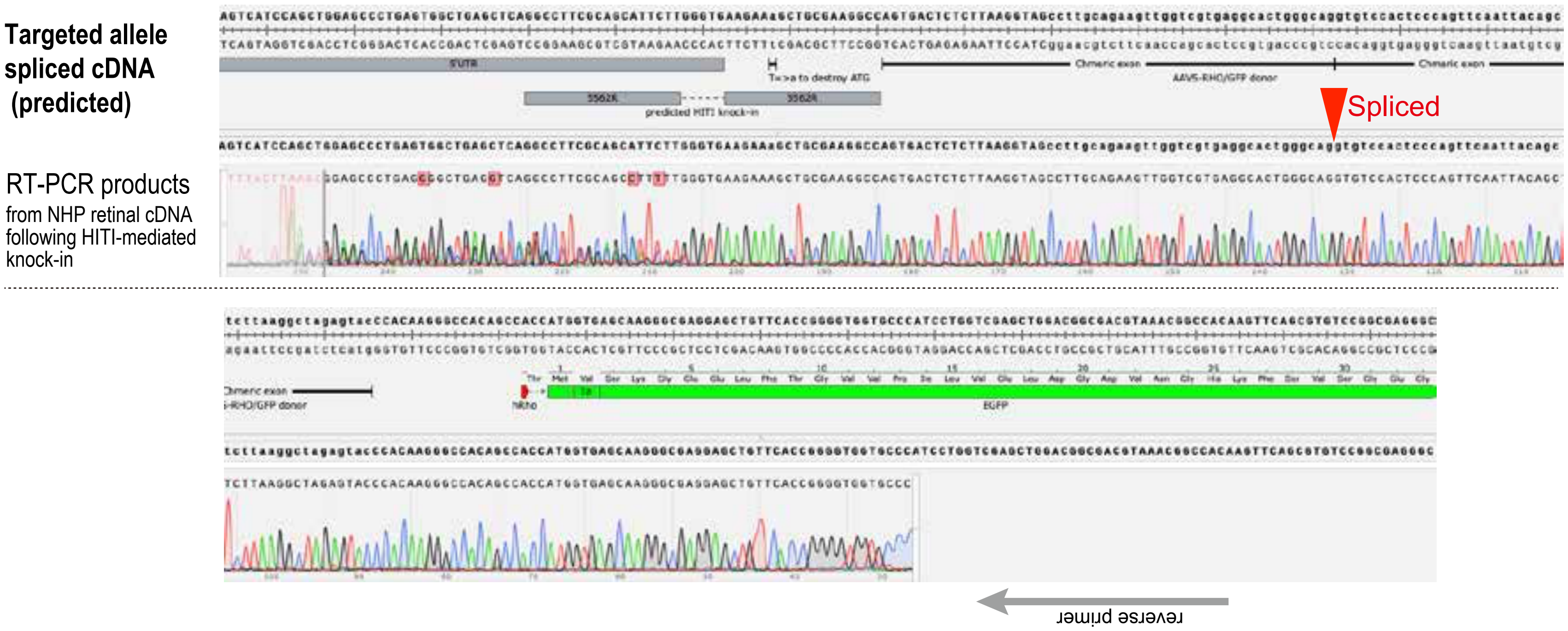


Figure S10. Onishi et al.

**Supplemental Table S1 DNA construction**

### DNAs

| Description in the text | cloned DNA/genes | cloning method |
|---|---|---|
| EGFP fragment for EGxxFP | EGFP (1-600bp) with stop codon, EGFP (120-720bp) | PCR with PrimerStarGXL (TaKaRa)*1 |
| mRho gRNA target | 5'-CTGAGCTCGCCAAGCAGCCTTGGTCTCTGTCTACGAAGAGCCCGTGGGGCAGCCTCGAGAGCC-3' | Oligonucleotide synthesis |
| hRHO ZF target (4000–6000 series) | 5'-TGGCTGAGCTCAGGCCTTCGCAGCATTCTTGGGTGGGAGCAGCCACGGGTCAGCCACAAG-3' | Oligonucleotide synthesis |
| hRHO ZF target (7000–9000 series) | 5'-TTGGGTGGGAGCAGCCACGGGTCAGCCACAAGGGCCACAGCCATGAATGGCACAGAAGG-3' | Oligonucleotide synthesis |
| SpCas9 | CDS of Streptococcus pyogenes Cas9 | PCR with PrimerStarGXL (TaKaRa)*1 |
| mRho-gRNA1 | 5'-TCTGTCTACGAAGAGCCCGTGGG-3' (used in Onishi et al. IOVS 2024) | Oligonucleotide synthesis |
| gRNA scaffold for mRho-gRNA1 | 5'-ATAGCAAGTTAAAATAAGGCTAGTCCGTTATCAACTTGAAAAAGTGGCACCGAGTCGGTGC-3' | Oligonucleotide synthesis |
| ZF modules | see TableS2 and TableS3 for the amino acid sequences | Artificially synthesized |
| ND1DDD | 5'-TTAGTGAAAGGTGAGATGGAGAAGAAAAAAAGTGACCTTAGACACAAGTTAAAGCATGTGCCACACGAGTACATCGAGTTAATCGAAATTGCGCAGGACTCTAAACAGAACAGGCTGTTTGAATTTAAGGTCGTTGAGTTCTTGAAAGAGGTTTACGATTACAACGGGAAGCACCTGGGGGGCAGTCGAAAACCAGACGGGGCCCTGTACACCAACGGGCTGAAAACTGACTACGGCATCATACTCGACACTAAAGCCTATAAGGACGGGTACAGCCTCCCTATTTCTCAGGCAGATGAGATGCAGGATTATGTGGATGAAAATAATAACCGTGATGCGATCATTAATCCCAACGAGTGGTGGAAAGTCTACCCGAACAGTATACTGGATTTCAAGTTCCTGTTCGTCTCCGGGTTCTTCAAGGGGGATTACAAAAAGCAGCTTGCTCGGGTCAGTAACTTAACTAAGCGCAAGGGGGCCGTCTTAAGTGTGGAACAGCTGCTCTTGGGGGGTGAGAAGATCAAGGACGGCTCCTTAACCCTGGAGGATGTCGGCGATAAGTTCAACAACGATGAAATAATCTTC-3' | Artificially synthesized |
| ND1RRR | 5'-CTGGTGAAGGGAGAGATGGAAAAGAAGAAGTCCGATCTCCGCCACAAATTAAAGCACGTACCTCACGAGTATATAGAACTGATCGAGATCGCCCAAGACAGCAAGCAGAACCGTCTTTTCGAGTTCAAGGTCGTGGAGTTTCTGAAAGAAGTCTATGATTATAACGGTAAACACCTGGGAGGGTCTAGGAAGCCTGACGGCGCGCTCTACACCAATGGCTTGAAGACCGACTATGGCATCATCCTGGATACCAAGGCATACAAGGATGGCTACTCCCTTCCCATTTCCCAGGCCCGAGAAATGCAGAGGTACGTGGACGAAAACAACAATCGGAACGCTATCATCAACCCCAATGAGTGGTGGAAAGTGTACCCTAACTCCATTTTAGATTTCAAGTTTCTCTTCGTGAGTGGATTCTTTAAGGGCGACTACAAGAAACAACTTGCGCGGGTTTCCAGGCTGACAAAAAGAAAAGGCGCAGTCCTGTCAGTAGAACAATTGTTACTCGGGGGCGAGAAAATTAAGGACGGCAGTTTGACTTTGGAAGACGTTGGTGATAAATTCAATAACGACGAGATCATTTTT-3' | Artificially synthesized |
| 3xNLS | 5'-CCAAAGAAGAAAAGAAAGGTTGCTGCGGCCCCTGCCGCTAAGCGCGTCAAGCTGGATGGGAGTGGCCCAAAAAAGAAGAGGAAAGTA-3' (SV40NLS-AAA-cMyc NLS-GSG-SV40NLS) | Oligonucleotide synthesis |
| mRho ZF target | 5'-TCGCCAAGCAGCCTTGGTCTCTGTCTACGAAGAGCCCGTGGGGCAGCCTCG-3' | Oligonucleotide synthesis |
| EF-1α promoter | human EEF1A promoter 1182 bp (chr6:73,520,048-73,521,229 in GRCh38/hg38) | Oligonucleotide synthesis |
| NLS | nuclear localization signal from c-Myc (PAAKRVKLD) | Oligonucleotide synthesis |
| Chimeric Intron | Chimeric intron of human beta globin and IgG | Oligonucleotide synthesis |
| synthetic pA | Synthetic polyadelylation signal (5'-AATAAAAGATCTTTATTTTCATTAGATCTGTGTGTTGGTTTTTTGTGTG-3') | Oligonucleotide synthesis |
| FurinP2A | Furin and P2A self digestive peptides with GSG linker (RRKR-GSG-ATNFSLLKQAGDVEENPGP) | Oligonucleotide synthesis |
| hRHO CDS | human RHO CDS (NM_000539.3) | PCR with PrimerStarGXL (TaKaRa)*1 |
| hRHO 3'-UTR | 3'-UTR from human RHO locus (chr3:129,533,726-129,535,393 in GRCh38/hg38) | PCR with PrimerStarGXL (TaKaRa)*1 |
| bRho300bp | bovine rhodopsin 300bp promoter (chr22:56,231,474-56,231,769 in ARS-UCD1.2/bosTau9) | PCR with PrimerStarGXL (TaKaRa)*1 |

### Plasmids

| Description in the text | cloned DNA casettes (5' to 3' direction) | cloning method | Plasmid backbone |
|---|---|---|---|
| pCAG-EGxxFP (for SpCas9 positive control) | [EGFP (1-600bp) with stop codon]-[mRho gRNA target]-[EGFP (120-720bp)] | InFusion HD Cloning Plus system (TaKaRa)*1 | pCAG (Addgene 11159) |
| pCAG-EGxxFP (for ZF-ND1 SSA, 4000–6000 series) | [EGFP (1-600bp) with stop codon]-[hRHO ZF target (4000–6000 series)]-[EGFP (120-720bp)] | InFusion HD Cloning Plus system (TaKaRa)*1 | pCAG (Addgene 11159) |
| pCAG-EGxxFP (for ZF-ND1 SSA, 7000–9000 series) | [EGFP (1-600bp) with stop codon]-[hRHO ZF target (7000–9000 series)]-[EGFP (120-720bp)] | InFusion HD Cloning Plus system (TaKaRa)*1 | pCAG (Addgene 11159) |
| pCAG-mCherry | mCherry | InFusion HD Cloning Plus system (TaKaRa)*1 | pCAG (Addgene 11159) |
| pbRho300-SpCas9 | bRho300bp, SpCas9, synthetic pA | InFusion HD Cloning Plus system (TaKaRa)*1 | pCAG (Addgene 11159) |
| pBAsi-U6-mRho-gRNA1 | mRho-gRNA1(PAM sequence is removed), gRNA scaffold | InFusion HD Cloning Plus system (TaKaRa)*1 | pBASi (TaKaRa) |
| pCMV-XXXX-S/AS-ND1DDD/RRR | SV40NLS-[XXXXS/XXXXAS]-TGGS-[ND1DDD/ND1RRR] | InFusion HD Cloning Plus system (TaKaRa)*1 | pcDNA3.1 (ThermoFischer) |
| pbRho300-5057S/5057AS | [bRho300bp]-[5057S/5057AS] | InFusion HD Cloning Plus system (TaKaRa)*1 | pCAG (Addgene 11159) |
| (Illustrated in Fig.S2D) | [mRhoZF target for 5057S-5057AS]*2-[ChimericIntron, mRhoCDS, FurinP2A, AcGFP, mRho3'UTR]-[mRhoZF target for 5057S-5057AS] | InFusion HD Cloning Plus system (TaKaRa)*1 | pLeaklessIII*3 |
| AAV5-ZFN | [synthetic pA]*2-[ND1RRR]*2-[5562AS-C6]*2-[EF-1a]-[CMV]-[5461S]-[ND1DDD]-[bGHpA] | InFusion HD Cloning Plus system (TaKaRa)*1 | pAAV*4 |
| AAV5-RHO | [hRHO ZF target for 5461S-5562AS with T to A mutation]*2-[ChimericIntron]-[hRHO CDS]-[hGHpA]-[hRHO ZF target for 5461S-5562AS] | InFusion HD Cloning Plus system (TaKaRa)*1 | pscAAV |
| AAV5-GFP | [hRHO ZF target for 5461S-5562AS with T to A mutation]*2-[ChimericIntron]-[AcGFP]-[hGHpA]-[hRHO ZF target for 5461S-5562AS] | InFusion HD Cloning Plus system (TaKaRa)*1 | pscAAV |
| (Illustrated in Fig.2B) Bicistronic | [3xNLS]-[5461S]-[ND1DDD]-[FuP2A]-[5562AS]-[ND1RRR] | InFusion HD Cloning Plus system (TaKaRa)*1 | pcDNA3.1 (ThermoFischer) |
| (Illustrated in Fig.2B) Bidirectional | [synthetic pA]*2-[ND1RRR]*2-[5562AS-C6]*2-[EF-1α]-[CMV]-[5461S]-[ND1DDD]-[bGHpA] | InFusion HD Cloning Plus system (TaKaRa)*1 | pcDNA3.1 (ThermoFischer) |
| (Illustrated in Fig.2D) AAV5-CMV-AcGFP-hRHO 3'-UTR | CMV-AcGFP-[hRHO 3'-UTR] | InFusion HD Cloning Plus system (TaKaRa)*1 | pAAV*4 |

*1 The reagents were used according to the manufacturer's instructions.
*2 These DNAs were inserted by reverse direction.
*3 Development. 2016 Sep 1; 143(17): 3216–3222.
*4 The 5' AAV ITR sequence bears a 11-bp deletion (5'-AAAGCCCGGGC-3').

### Supplemental Table S2 Target sequences of ZF pairs for human RHO 5'-UTR

ZF pairs used in this study | ZF modules (Wright et al. 2006)

| No | Name | antisense | Spacer | sense | Name | F1 | F2 | F3 | F4 | F5 | F6 | Amino acid sequence (6ZF modules) |
|---|---|---|---|---|---|---|---|---|---|---|---|---|
| 1 | 4350S | 5'- TGG CTG AGC TCA GGC CTT | CGCAGC | ATT CTT GGG TGG GAG CAG -3' | 4350S | ZF91 | ZF62 | ZF106 | ZF58 | ZF103 | ZF88 | PGEKPYKCPECGKSFSRADNLTEHQRTHTGEKPYKCPECGKSFSRSDNLVRHQRTHTGEKPYKCPECGKSFSRSDHLTTHQRTHTGEKPYKCPECGKSFSRSDKLVRHQRTHTGEKPYKCPECGKSFSTTGALTEHQRTHTGEKPYKCPECGKSFSHKNALQNHQRTH |
| | 4350AS | 3'- ACC GAC TCG AGT CCG GAA | GCGTCG | TAA GAA CCC ACC CTC GTC -5' | 4350AS | ZF93 | ZF91 | ZF72 | ZF105 | ZF73 | ZF76 | PGEKPYKCPECGKSFSTSHSLTEHQRTHTGEKPYKCPECGKSFSRADNLTEHQRTHTGEKPYKCPECGKSFSTSGELVRHQRTHTGEKPYKCPECGKSFSQAGHLASHQRTHTGEKPYKCPECGKSFSDCRDLARHQRTHTGEKPYKCPECGKSFSRKDNLKNHQRTH |
| 2 | 4653S | 5'- CTG AGC TCA GGC CTT CGC | AGCATT | CTT GGG TGG GAG CAG CCA -3' | 4653S | ZF93 | ZF91 | ZF62 | ZF106 | ZF58 | ZF103 | PGEKPYKCPECGKSFSTSHSLTEHQRTHTGEKPYKCPECGKSFSRADNLTEHQRTHTGEKPYKCPECGKSFSRSDNLVRHQRTHTGEKPYKCPECGKSFSRSDHLTTHQRTHTGEKPYKCPECGKSFSRSDKLVRHQRTHTGEKPYKCPECGKSFSTTGALTEHQRTH |
| | 4653AS | 3'- GAC TCG AGT CCG GAA GCG | TCGTAA | GAA CCC ACC CTC GTC GGT -5' | 4653AS | ZF91 | ZF72 | ZF105 | ZF73 | ZF76 | ZF70 | PGEKPYKCPECGKSFSRADNLTEHQRTHTGEKPYKCPECGKSFSTSGELVRHQRTHTGEKPYKCPECGKSFSQAGHLASHQRTHTGEKPYKCPECGKSFSDCRDLARHQRTHTGEKPYKCPECGKSFSRKDNLKNHQRTHTGEKPYKCPECGKSFSRSDDLVRHQRTH |
| 3 | 4956S | 5'- AGC TCA GGC CTT CGC AGC | ATTCTT | GGG TGG GAG CAG CCA CGG -3' | 4956S | ZF99 | ZF93 | ZF91 | ZF62 | ZF106 | ZF58 | PGEKPYKCPECGKSFSRSDKLTEHQRTHTGEKPYKCPECGKSFSTSHSLTEHQRTHTGEKPYKCPECGKSFSRADNLTEHQRTHTGEKPYKCPECGKSFSRSDNLVRHQRTHTGEKPYKCPECGKSFSRSDHLTTHQRTHTGEKPYKCPECGKSFSRSDKLVRHQRTH |
| | 4956AS | 3'- TCG AGT CCG GAA GCG TCG | TAAGAA | CCC ACC CTC GTC GGT GCC -5' | 4956AS | ZF72 | ZF105 | ZF73 | ZF76 | ZF70 | ZF72 | PGEKPYKCPECGKSFSTSGELVRHQRTHTGEKPYKCPECGKSFSQAGHLASHQRTHTGEKPYKCPECGKSFSDCRDLARHQRTHTGEKPYKCPECGKSFSRKDNLKNHQRTHTGEKPYKCPECGKSFSRSDDLVRHQRTHTGEKPYKCPECGKSFSTSGELVRHQRTH |
| 4 | 5158S | 5'- CTC AGG CCT TCG CAG CAT | TCTTGG | GTG GGA GCA GCC ACG GGT -3' | 5158S | ZF60 | ZF80 | ZF73 | ZF71 | ZF59 | ZF66 | PGEKPYKCPECGKSFSTSGHLVRHQRTHTGEKPYKCPECGKSFSRTDTLRDHQRTHTGEKPYKCPECGKSFSDCRDLARHQRTHTGEKPYKCPECGKSFSQSGDLRRHQRTHTGEKPYKCPECGKSFSQRAHLERHQRTHTGEKPYKCPECGKSFSRSDELVRHQRTH |
| | 5158AS | 3'- GAG TCC GGA AGC GTC GTA | AGAACC | CAC CCT CGT CGG TGC CCA -5' | 5158AS | ZF62 | ZF96 | ZF84 | ZF97 | ZF102 | ZF87 | PGEKPYKCPECGKSFSRSDNLVRHQRTHTGEKPYKCPECGKSFSTKNSLTEHQRTHTGEKPYKCPECGKSFSRSDHLTNHQRTHTGEKPYKCPECGKSFSQSGHLTEHQRTHTGEKPYKCPECGKSFSRNDALTEHQRTHTGEKPYKCPECGKSFSRRDELNVHQRTH |
| 5 | 5259S | 5'- TCA GGC CTT CGC AGC ATT | CTTGGG | TGG GAG CAG CCA CGG GTC -3' | 5259S | ZF69 | ZF99 | ZF93 | ZF91 | ZF62 | ZF106 | PGEKPYKCPECGKSFSDPGALVRHQRTHTGEKPYKCPECGKSFSRSDKLTEHQRTHTGEKPYKCPECGKSFSTSHSLTEHQRTHTGEKPYKCPECGKSFSRADNLTEHQRTHTGEKPYKCPECGKSFSRSDNLVRHQRTHTGEKPYKCPECGKSFSRSDHLTTHQRTH |
| | 5259AS | 3'- AGT CCG GAA GCG TCG TAA | GAACCC | ACC CTC GTC GGT GCC CAG -5' | 5259AS | ZF105 | ZF73 | ZF76 | ZF70 | ZF72 | ZF77 | PGEKPYKCPECGKSFSQAGHLASHQRTHTGEKPYKCPECGKSFSDCRDLARHQRTHTGEKPYKCPECGKSFSRKDNLKNHQRTHTGEKPYKCPECGKSFSRSDDLVRHQRTHTGEKPYKCPECGKSFSTSGELVRHQRTHTGEKPYKCPECGKSFSTTGNLTVHQRTH |
| 6 | 5461S | 5'- AGG CCT TCG CAG CAT TCT | TGGGTG | GGA GCA GCC ACG GGT CAG -3' | 5461S | ZF91 | ZF60 | ZF80 | ZF73 | ZF71 | ZF59 | PGEKPYKCPECGKSFSRADNLTEHQRTHTGEKPYKCPECGKSFSTSGHLVRHQRTHTGEKPYKCPECGKSFSRTDTLRDHQRTHTGEKPYKCPECGKSFSDCRDLARHQRTHTGEKPYKCPECGKSFSQSGDLRRHQRTHTGEKPYKCPECGKSFSQRAHLERHQRTH |
| | 5461AS | 3'- TCC GGA AGC GTC GTA AGA | ACCCAC | CCT CGT CGG TGC CCA GTC -5' | 5461AS | ZF96 | ZF84 | ZF97 | ZF102 | ZF87 | ZF82 | PGEKPYKCPECGKSFSTKNSLTEHQRTHTGEKPYKCPECGKSFSRSDHLTNHQRTHTGEKPYKCPECGKSFSQSGHLTEHQRTHTGEKPYKCPECGKSFSRNDALTEHQRTHTGEKPYKCPECGKSFSRRDELNVHQRTHTGEKPYKCPECGKSFSQLAHLRAHQRTH |
| 7 | 5562S | 5'- GGC CTT CGC AGC ATT CTT | GGGTGG | GAG CAG CCA CGG GTC AGC -3' | 5562S | ZF83 | ZF69 | ZF99 | ZF93 | ZF91 | ZF62 | PGEKPYKCPECGKSFSERSHLREHQRTHTGEKPYKCPECGKSFSDPGALVRHQRTHTGEKPYKCPECGKSFSRSDKLTEHQRTHTGEKPYKCPECGKSFSTSHSLTEHQRTHTGEKPYKCPECGKSFSRADNLTEHQRTHTGEKPYKCPECGKSFSRSDNLVRHQRTH |
| | 5562AS(C1) | 3'- CCG GAA GCG TCG TAA GAA | CCCACC | CTC GTC GGT GCC CAG TCG -5' | 5562AS(C1) | ZF73 | ZF76 | ZF70 | ZF72 | ZF77 | ZF76 | PGEKPYKCPECGKSFSDCRDLARHQRTHTGEKPYKCPECGKSFSRKDNLKNHQRTHTGEKPYKCPECGKSFSRSDDLVRHQRTHTGEKPYKCPECGKSFSTSGELVRHQRTHTGEKPYKCPECGKSFSTTGNLTVHQRTHTGEKPYKCPECGKSFSRKDNLKNHQRTH |
| 8 | 5764S | 5'- CCT TCG CAG CAT TCT TGG | GTGGGA | GCA GCC ACG GGT CAG CCA -3' | 5764S | ZF93 | ZF91 | ZF60 | ZF80 | ZF73 | ZF71 | PGEKPYKCPECGKSFSTSHSLTEHQRTHTGEKPYKCPECGKSFSRADNLTEHQRTHTGEKPYKCPECGKSFSTSGHLVRHQRTHTGEKPYKCPECGKSFSRTDTLRDHQRTHTGEKPYKCPECGKSFSDCRDLARHQRTHTGEKPYKCPECGKSFSQSGDLRRHQRTH |
| | 5764AS | 3'- GGA AGC GTC GTA AGA ACC | CACCCT | CGT CGG TGC CCA GTC GGT -5' | 5764AS | ZF84 | ZF97 | ZF102 | ZF87 | ZF82 | ZF93 | PGEKPYKCPECGKSFSRSDHLTNHQRTHTGEKPYKCPECGKSFSQSGHLTEHQRTHTGEKPYKCPECGKSFSRNDALTEHQRTHTGEKPYKCPECGKSFSRRDELNVHQRTHTGEKPYKCPECGKSFSQLAHLRAHQRTHTGEKPYKCPECGKSFSTSHSLTEHQRTH |
| 9 | 5865S | 5'- CTT CGC AGC ATT CTT GGG | TGGGAG | CAG CCA CGG GTC AGC CAC -3' | 5865S | ZF90 | ZF83 | ZF69 | ZF99 | ZF93 | ZF91 | PGEKPYKCPECGKSFSSKKALTEHQRTHTGEKPYKCPECGKSFSERSHLREHQRTHTGEKPYKCPECGKSFSDPGALVRHQRTHTGEKPYKCPECGKSFSRSDKLTEHQRTHTGEKPYKCPECGKSFSTSHSLTEHQRTHTGEKPYKCPECGKSFSRADNLTEHQRTH |
| | 5865AS | 3'- GAA GCG TCG TAA GAA CCC | ACCCTC | GTC GGT GCC CAG TCG GTG -5' | 5865AS | ZF76 | ZF70 | ZF72 | ZF77 | ZF76 | ZF94 | PGEKPYKCPECGKSFSRKDNLKNHQRTHTGEKPYKCPECGKSFSRSDDLVRHQRTHTGEKPYKCPECGKSFSTSGELVRHQRTHTGEKPYKCPECGKSFSTTGNLTVHQRTHTGEKPYKCPECGKSFSRKDNLKNHQRTHTGEKPYKCPECGKSFSSKKHLAEHQRTH |
| 10 | 6067S | 5'- TCG CAG CAT TCT TGG GTG | GGAGCA | GCC ACG GGT CAG CCA CAA -3' | 6067S | ZF89 | ZF93 | ZF91 | ZF60 | ZF80 | ZF73 | PGEKPYKCPECGKSFSQSGNLTEHQRTHTGEKPYKCPECGKSFSTSHSLTEHQRTHTGEKPYKCPECGKSFSRADNLTEHQRTHTGEKPYKCPECGKSFSTSGHLVRHQRTHTGEKPYKCPECGKSFSRTDTLRDHQRTHTGEKPYKCPECGKSFSDCRDLARHQRTH |
| | 6067AS | 3'- AGC GTC GTA AGA ACC CAC | CCTCGT | CGG TGC CCA GTC GGT GTT -5' | 6067AS | ZF97 | ZF102 | ZF87 | ZF82 | ZF93 | ZF90 | PGEKPYKCPECGKSFSQSGHLTEHQRTHTGEKPYKCPECGKSFSRNDALTEHQRTHTGEKPYKCPECGKSFSRRDELNVHQRTHTGEKPYKCPECGKSFSQLAHLRAHQRTHTGEKPYKCPECGKSFSTSHSLTEHQRTHTGEKPYKCPECGKSFSSKKALTEHQRTH |
| 11 | 6168S | 5'- CGC AGC ATT CTT GGG TGG | GAGCAG | CCA CGG GTC AGC CAC AAG -3' | 6168S | ZF76 | ZF90 | ZF83 | ZF69 | ZF99 | ZF93 | PGEKPYKCPECGKSFSRKDNLKNHQRTHTGEKPYKCPECGKSFSSKKALTEHQRTHTGEKPYKCPECGKSFSERSHLREHQRTHTGEKPYKCPECGKSFSDPGALVRHQRTHTGEKPYKCPECGKSFSRSDKLTEHQRTHTGEKPYKCPECGKSFSTSHSLTEHQRTH |
| | 6168AS | 3'- GCG TCG TAA GAA CCC ACC | CTCGTC | GGT GCC CAG TCG GTG TTC -5' | 6168AS | ZF70 | ZF72 | ZF77 | ZF76 | ZF94 | ZF93 | PGEKPYKCPECGKSFSRSDDLVRHQRTHTGEKPYKCPECGKSFSTSGELVRHQRTHTGEKPYKCPECGKSFSTTGNLTVHQRTHTGEKPYKCPECGKSFSRKDNLKNHQRTHTGEKPYKCPECGKSFSSKKHLAEHQRTHTGEKPYKCPECGKSFSTSHSLTEHQRTH |
| 12 | 7178S | 5'- TTG GGT GGG AGC AGC CAC | GGGTCA | GCC ACA AGG GCC ACA GCC -3' | 7178S | ZF73 | ZF78 | ZF73 | ZF84 | ZF78 | ZF73 | PGEKPYKCPECGKSFSDCRDLARHQRTHTGEKPYKCPECGKSFSSPADLTRHQRTHTGEKPYKCPECGKSFSDCRDLARHQRTHTGEKPYKCPECGKSFSRSDHLTNHQRTHTGEKPYKCPECGKSFSSPADLTRHQRTHTGEKPYKCPECGKSFSDCRDLARHQRTH |
| | 7178AS | 3'- AAC CCA CCC TCG TCG GTG | CCCAGT | CGG TGT TCC CGG TGT CGG -5' | 7178AS | ZF89 | ZF79 | ZF94 | ZF72 | ZF72 | ZF66 | PGEKPYKCPECGKSFSQSGNLTEHQRTHTGEKPYKCPECGKSFSDKKDLTRHQRTHTGEKPYKCPECGKSFSSKKHLAEHQRTHTGEKPYKCPECGKSFSTSGELVRHQRTHTGEKPYKCPECGKSFSTSGELVRHQRTHTGEKPYKCPECGKSFSRSDELVRHQRTH |
| 13 | 7481S | 5'- GGT GGG AGC AGC CAC GGG | TCAGCC | ACA AGG GCC ACA GCC ATG -3' | 7481S | ZF87 | ZF73 | ZF78 | ZF73 | ZF84 | ZF78 | GEKPYKCPECGKSFSRRDELNVHQRTHTGEKPYKCPECGKSFSDCRDLARHQRTHTGEKPYKCPECGKSFSSPADLTRHQRTHTGEKPYKCPECGKSFSDCRDLARHQRTHTGEKPYKCPECGKSFSRSDHLTNHQRTHTGEKPYKCPECGKSFSSPADLTRHQRTH |
| | 7481AS | 3'- CCA CCC TCG TCG GTG CCC | AGTCGG | TGT TCC CGG TGT CGG TAC -5' | 7481AS | ZF79 | ZF94 | ZF72 | ZF72 | ZF66 | ZF94 | GEKPYKCPECGKSFSDKKDLTRHQRTHTGEKPYKCPECGKSFSSKKHLAEHQRTHTGEKPYKCPECGKSFSTSGELVRHQRTHTGEKPYKCPECGKSFSTSGELVRHQRTHTGEKPYKCPECGKSFSRSDELVRHQRTHTGEKPYKCPECGKSFSSKKHLAEHQRTH |
| 14 | 7784S | 5'- GGG AGC AGC CAC GGG TCA | GCCACA | AGG GCC ACA GCC ATG AAT -3' | 7784S | ZF77 | ZF87 | ZF73 | ZF78 | ZF73 | ZF84 | GEKPYKCPECGKSFSTTGNLTVHQRTHTGEKPYKCPECGKSFSRRDELNVHQRTHTGEKPYKCPECGKSFSDCRDLARHQRTHTGEKPYKCPECGKSFSSPADLTRHQRTHTGEKPYKCPECGKSFSDCRDLARHQRTHTGEKPYKCPECGKSFSRSDHLTNHQRTH |
| | 7784AS | 3'- CCC TCG TCG GTG CCC AGT | CGGTGT | TCC CGG TGT CGG TAC TTA -5' | 7784AS | ZF94 | ZF72 | ZF72 | ZF66 | ZF94 | ZF105 | GEKPYKCPECGKSFSSKKHLAEHQRTHTGEKPYKCPECGKSFSTSGELVRHQRTHTGEKPYKCPECGKSFSTSGELVRHQRTHTGEKPYKCPECGKSFSRSDELVRHQRTHTGEKPYKCPECGKSFSSKKHLAEHQRTHTGEKPYKCPECGKSFSQAGHLASHQRTH |
| 15 | 8087S | 5'- AGC AGC CAC GGG TCA GCC | ACAAGG | GCC ACA GCC ATG AAT GGC -3' | 8087S | ZF61 | ZF77 | ZF87 | ZF73 | ZF78 | ZF73 | GEKPYKCPECGKSFSDPGHLVRHQRTHTGEKPYKCPECGKSFSTTGNLTVHQRTHTGEKPYKCPECGKSFSRRDELNVHQRTHTGEKPYKCPECGKSFSDCRDLARHQRTHTGEKPYKCPECGKSFSSPADLTRHQRTHTGEKPYKCPECGKSFSDCRDLARHQRTH |
| | 8087AS | 3'- TCG TCG GTG CCC AGT CGG | TGTTCC | CGG TGT CGG TAC TTA CCG -5' | 8087AS | ZF72 | ZF72 | ZF66 | ZF94 | ZF105 | ZF61 | GEKPYKCPECGKSFSTSGELVRHQRTHTGEKPYKCPECGKSFSTSGELVRHQRTHTGEKPYKCPECGKSFSRSDELVRHQRTHTGEKPYKCPECGKSFSSKKHLAEHQRTHTGEKPYKCPECGKSFSQAGHLASHQRTHTGEKPYKCPECGKSFSDPGHLVRHQRTH |
| 16 | 8289S | 5'- CAG CCA CGG GTC AGC CAC | AAGGGC | CAC AGC CAT GAA TGG CAC -3' | 8289S | ZF90 | ZF106 | ZF63 | ZF92 | ZF83 | ZF90 | GEKPYKCPECGKSFSSKKALTEHQRTHTGEKPYKCPECGKSFSRSDHLTTHQRTHTGEKPYKCPECGKSFSQSSNLVRHQRTHTGEKPYKCPECGKSFSTSGNLTEHQRTHTGEKPYKCPECGKSFSERSHLREHQRTHTGEKPYKCPECGKSFSSKKALTEHQRTH |
| | 8289AS | 3'- GTC GGT GCC CAG TCG GTG | TTCCCG | GTG TCG GTA CTT ACC GTG -5' | 8289AS | ZF102 | ZF106 | ZF95 | ZF65 | ZF72 | ZF66 | GEKPYKCPECGKSFSRNDALTEHQRTHTGEKPYKCPECGKSFSRSDHLTTHQRTHTGEKPYKCPECGKSFSRNDTLTEHQRTHTGEKPYKCPECGKSFSDPGNLVRHQRTHTGEKPYKCPECGKSFSTSGELVRHQRTHTGEKPYKCPECGKSFSRSDELVRHQRTH |
| 17 | 8592S | 5'- CCA CGG GTC AGC CAC AAG | GGCCAC | AGC CAT GAA TGG CAC AGA -3' | 8592S | ZF82 | ZF90 | ZF106 | ZF63 | ZF92 | ZF83 | GEKPYKCPECGKSFSQLAHLRAHQRTHTGEKPYKCPECGKSFSSKKALTEHQRTHTGEKPYKCPECGKSFSRSDHLTTHQRTHTGEKPYKCPECGKSFSQSSNLVRHQRTHTGEKPYKCPECGKSFSTSGNLTEHQRTHTGEKPYKCPECGKSFSERSHLREHQRTH |
| | 8592AS | 3'- GGT GCC CAG TCG GTG TTC | CCGGTG | TCG GTA CTT ACC GTG TCT -5' | 8592AS | ZF106 | ZF95 | ZF65 | ZF72 | ZF66 | ZF103 | GEKPYKCPECGKSFSRSDHLTTHQRTHTGEKPYKCPECGKSFSRNDTLTEHQRTHTGEKPYKCPECGKSFSDPGNLVRHQRTHTGEKPYKCPECGKSFSTSGELVRHQRTHTGEKPYKCPECGKSFSRSDELVRHQRTHTGEKPYKCPECGKSFSTTGALTEHQRTH |
| 18 | 8895S | 5'- CGG GTC AGC CAC AAG GGC | CACAGC | CAT GAA TGG CAC AGA AGG -3' | 8895S | ZF84 | ZF82 | ZF90 | ZF106 | ZF63 | ZF92 | GEKPYKCPECGKSFSRSDHLTNHQRTHTGEKPYKCPECGKSFSQLAHLRAHQRTHTGEKPYKCPECGKSFSSKKALTEHQRTHTGEKPYKCPECGKSFSRSDHLTTHQRTHTGEKPYKCPECGKSFSQSSNLVRHQRTHTGEKPYKCPECGKSFSTSGNLTEHQRTH |
| | 8895AS | 3'- GCC CAG TCG GTG TTC CCG | GTGTCG | GTA CTT ACC GTG TCT TCC -5' | 8895AS | ZF95 | ZF65 | ZF72 | ZF66 | ZF103 | ZF73 | GEKPYKCPECGKSFSRNDTLTEHQRTHTGEKPYKCPECGKSFSDPGNLVRHQRTHTGEKPYKCPECGKSFSTSGELVRHQRTHTGEKPYKCPECGKSFSRSDELVRHQRTHTGEKPYKCPECGKSFSTTGALTEHQRTHTGEKPYKCPECGKSFSDCRDLARHQRTH |
| 19 | 4955S | 5'- AGC TCA GGC CTT CGC AGC | ATTCT | TGG GTG GGA GCA GCC ACG -3' | 4955S | ZF80 | ZF73 | ZF71 | ZF59 | ZF66 | ZF106 | GEKPYKCPECGKSFSRTDTLRDHQRTHTGEKPYKCPECGKSFSDCRDLARHQRTHTGEKPYKCPECGKSFSQSGDLRRHQRTHTGEKPYKCPECGKSFSQRAHLERHQRTHTGEKPYKCPECGKSFSRSDELVRHQRTHTGEKPYKCPECGKSFSRSDHLTTHQRTH |
| | 4956AS | 3'- TCG AGT CCG GAA GCG TCG | TAAGA | ACC CAC CCT CGT CGG TGC -5' | 4956AS | ZF72 | ZF105 | ZF73 | ZF76 | ZF70 | ZF72 | GEKPYKCPECGKSFSTSGELVRHQRTHTGEKPYKCPECGKSFSQAGHLASHQRTHTGEKPYKCPECGKSFSDCRDLARHQRTHTGEKPYKCPECGKSFSRKDNLKNHQRTHTGEKPYKCPECGKSFSRSDDLVRHQRTHTGEKPYKCPECGKSFSTSGELVRHQRTH |
| 20 | 5157S | 5'- CTC AGG CCT TCG CAG CAT | TCTTG | GGT GGG AGC AGC CAC GGG -3' | 5157S | ZF58 | ZF90 | ZF83 | ZF83 | ZF58 | ZF60 | GEKPYKCPECGKSFSRSDKLVRHQRTHTGEKPYKCPECGKSFSSKKALTEHQRTHTGEKPYKCPECGKSFSERSHLREHQRTHTGEKPYKCPECGKSFSERSHLREHQRTHTGEKPYKCPECGKSFSRSDKLVRHQRTHTGEKPYKCPECGKSFSTSGHLVRHQRTH |
| | 5158AS | 3'- GAG TCC GGA AGC GTC GTA | AGAAC | CCA CCC TCG TCG GTG CCC -5' | 5158AS | ZF62 | ZF96 | ZF84 | ZF97 | ZF102 | ZF87 | GEKPYKCPECGKSFSRSDNLVRHQRTHTGEKPYKCPECGKSFSTKNSLTEHQRTHTGEKPYKCPECGKSFSRSDHLTNHQRTHTGEKPYKCPECGKSFSQSGHLTEHQRTHTGEKPYKCPECGKSFSRNDALTEHQRTHTGEKPYKCPECGKSFSRRDELNVHQRTH |
| 21 | 5158S | 5'- TCA GGC CTT CGC AGC ATT | CTTGG | GTG GGA GCA GCC ACG GGT -3' | 5158S | ZF60 | ZF80 | ZF73 | ZF71 | ZF59 | ZF66 | GEKPYKCPECGKSFSTSGHLVRHQRTHTGEKPYKCPECGKSFSRTDTLRDHQRTHTGEKPYKCPECGKSFSDCRDLARHQRTHTGEKPYKCPECGKSFSQSGDLRRHQRTHTGEKPYKCPECGKSFSQRAHLERHQRTHTGEKPYKCPECGKSFSRSDELVRHQRTH |
| | 5259AS | 3'- AGT CCG GAA GCG TCG TAA | GAACC | CAC CCT CGT CGG TGC CCA -5' | 5259AS | ZF105 | ZF73 | ZF76 | ZF70 | ZF72 | ZF77 | GEKPYKCPECGKSFSQAGHLASHQRTHTGEKPYKCPECGKSFSDCRDLARHQRTHTGEKPYKCPECGKSFSRKDNLKNHQRTHTGEKPYKCPECGKSFSRSDDLVRHQRTHTGEKPYKCPECGKSFSTSGELVRHQRTHTGEKPYKCPECGKSFSTTGNLTVHQRTH |
| 22 | 5461S | 5'- GGC CTT CGC AGC ATT CTT | GGGTG | GGA GCA GCC ACG GGT CAG -3' | 5461S | ZF91 | ZF60 | ZF80 | ZF73 | ZF71 | ZF59 | GEKPYKCPECGKSFSRADNLTEHQRTHTGEKPYKCPECGKSFSTSGHLVRHQRTHTGEKPYKCPECGKSFSRTDTLRDHQRTHTGEKPYKCPECGKSFSDCRDLARHQRTHTGEKPYKCPECGKSFSQSGDLRRHQRTHTGEKPYKCPECGKSFSQRAHLERHQRTH |
| | 5562AS(C1) | 3'- CCG GAA GCG TCG TAA GAA | CCCAC | CCT CGT CGG TGC CCA GTC -5' | 5562AS(C1) | ZF73 | ZF76 | ZF70 | ZF72 | ZF77 | ZF76 | GEKPYKCPECGKSFSDCRDLARHQRTHTGEKKFACPECPKRFMRSDNLTQHIKTHTGEKPYKCPECGKSFSRSDDLVRHQRTHTGEKPYKCPECGKSFSTSGELVRHQRTHTGEKPYKCPECGKSFSTTGNLTVHQRTHTGEKKFACPECPKRFMRSDNLTQHIKTH |
| 23 | 5764S | 5'- CTT CGC AGC ATT CTT GGG | TGGGA | GCA GCC ACG GGT CAG CCA -3' | 5764S | ZF93 | ZF91 | ZF60 | ZF80 | ZF73 | ZF71 | GEKPYKCPECGKSFSTSHSLTEHQRTHTGEKPYKCPECGKSFSRADNLTEHQRTHTGEKPYKCPECGKSFSTSGHLVRHQRTHTGEKPYKCPECGKSFSRTDTLRDHQRTHTGEKPYKCPECGKSFSDCRDLARHQRTHTGEKPYKCPECGKSFSQSGDLRRHQRTH |
| | 5865AS | 3'- GAA GCG TCG TAA GAA CCC | ACCCT | CGT CGG TGC CCA GTC GGT -5' | 5865AS | ZF76 | ZF70 | ZF72 | ZF77 | ZF76 | ZF94 | GEKPYKCPECGKSFSRKDNLKNHQRTHTGEKPYKCPECGKSFSRSDDLVRHQRTHTGEKPYKCPECGKSFSTSGELVRHQRTHTGEKPYKCPECGKSFSTTGNLTVHQRTHTGEKPYKCPECGKSFSRKDNLKNHQRTHTGEKPYKCPECGKSFSSKKHLAEHQRTH |
| 24 | 6067S | 5'- CGC AGC ATT CTT GGG TGG | GAGCA | GCC ACG GGT CAG CCA CAA -3' | 6067S | ZF89 | ZF93 | ZF91 | ZF60 | ZF80 | ZF73 | GEKPYKCPECGKSFSQSGNLTEHQRTHTGEKPYKCPECGKSFSTSHSLTEHQRTHTGEKPYKCPECGKSFSRADNLTEHQRTHTGEKPYKCPECGKSFSTSGHLVRHQRTHTGEKPYKCPECGKSFSRTDTLRDHQRTHTGEKPYKCPECGKSFSDCRDLARHQRTH |
| | 6168AS | 3'- GCG TCG TAA GAA CCC ACC | CTCGT | CGG TGC CCA GTC GGT GTT -5' | 6168AS | ZF70 | ZF72 | ZF77 | ZF76 | ZF94 | ZF93 | GEKPYKCPECGKSFSRSDDLVRHQRTHTGEKPYKCPECGKSFSTSGELVRHQRTHTGEKPYKCPECGKSFSTTGNLTVHQRTHTGEKPYKCPECGKSFSRKDNLKNHQRTHTGEKPYKCPECGKSFSSKKHLAEHQRTHTGEKPYKCPECGKSFSTSHSLTEHQRTH |
| 25 | 6874S | 5'- TTC TTG GGT GGG AGC AGC | CACGG | GTC AGC CAC AAG GGC CAC -3' | 6874S | ZF90 | ZF61 | ZF76 | ZF90 | ZF83 | ZF69 | GEKPYKCPECGKSFSSKKALTEHQRTHTGEKPYKCPECGKSFSDPGHLVRHQRTHTGEKPYKCPECGKSFSRKDNLKNHQRTHTGEKPYKCPECGKSFSSKKALTEHQRTHTGEKPYKCPECGKSFSERSHLREHQRTHTGEKPYKCPECGKSFSDPGALVRHQRTH |
| | 6874AS | 3'- AAG AAC CCA CCC TCG TCG | GTGCC | CAG TCG GTG TTC CCG GTG -5' | 6874AS | ZF63 | ZF89 | ZF79 | ZF94 | ZF72 | ZF72 | GEKPYKCPECGKSFSQSSNLVRHQRTHTGEKPYKCPECGKSFSQSGNLTEHQRTHTGEKPYKCPECGKSFSDKKDLTRHQRTHTGEKPYKCPECGKSFSSKKHLAEHQRTHTGEKPYKCPECGKSFSTSGELVRHQRTHTGEKPYKCPECGKSFSTSGELVRHQRTH |
| 26 | 7177S | 5'- TTG GGT GGG AGC AGC CAC | GGGTC | AGC CAC AAG GGC CAC AGC -3' | 7177S | ZF83 | ZF90 | ZF61 | ZF76 | ZF90 | ZF83 | GEKPYKCPECGKSFSERSHLREHQRTHTGEKPYKCPECGKSFSSKKALTEHQRTHTGEKPYKCPECGKSFSDPGHLVRHQRTHTGEKPYKCPECGKSFSRKDNLKNHQRTHTGEKPYKCPECGKSFSSKKALTEHQRTHTGEKPYKCPECGKSFSERSHLREHQRTH |
| | 7178AS | 3'- AAC CCA CCC TCG TCG GTG | CCCAG | TCG GTG TTC CCG GTG TCG -5' | 7178AS | ZF89 | ZF79 | ZF94 | ZF72 | ZF72 | ZF66 | GEKPYKCPECGKSFSQSGNLTEHQRTHTGEKPYKCPECGKSFSDKKDLTRHQRTHTGEKPYKCPECGKSFSSKKHLAEHQRTHTGEKPYKCPECGKSFSTSGELVRHQRTHTGEKPYKCPECGKSFSTSGELVRHQRTHTGEKPYKCPECGKSFSRSDELVRHQRTH |
| 27 | 7480S | 5'- GGT GGG AGC AGC CAC GGG | TCAGC | CAC AAG GGC CAC AGC CAT -3' | 7480S | ZF92 | ZF83 | ZF90 | ZF61 | ZF76 | ZF90 | GEKPYKCPECGKSFSDKKDLTRHQRTHTGEKPYKCPECGKSFSSKKHLAEHQRTHTGEKPYKCPECGKSFSTSGELVRHQRTHTGEKPYKCPECGKSFSTSGELVRHQRTHTGEKPYKCPECGKSFSRSDELVRHQRTHTGEKPYKCPECGKSFSSKKHLAEHQRTH |
| | 7481AS | 3'- CCA CCC TCG TCG GTG CCC | AGTCG | GTG TTC CCG GTG TCG GTA -5' | 7481AS | ZF79 | ZF94 | ZF72 | ZF72 | ZF66 | ZF94 | GEKPYKCPECGKSFSDKKDLTRHQRTHTGEKPYKCPECGKSFSSKKHLAEHQRTHTGEKPYKCPECGKSFSTSGELVRHQRTHTGEKPYKCPECGKSFSTSGELVRHQRTHTGEKPYKCPECGKSFSRSDELVRHQRTHTGEKPYKCPECGKSFSSKKHLAEHQRTH |
| 28 | 7783S | 5'- GGG AGC AGC CAC GGG TCA | GCCAC | AAG GGC CAC AGC CAT GAA -3' | 7783S | ZF63 | ZF92 | ZF83 | ZF90 | ZF61 | ZF76 | GEKPYKCPECGKSFSQSSNLVRHQRTHTGEKPYKCPECGKSFSTSGNLTEHQRTHTGEKPYKCPECGKSFSERSHLREHQRTHTGEKPYKCPECGKSFSSKKALTEHQRTHTGEKPYKCPECGKSFSDPGHLVRHQRTHTGEKPYKCPECGKSFSRKDNLKNHQRTH |
| | 7784AS | 3'- CCC TCG TCG GTG CCC AGT | CGGTG | TTC CCG GTG TCG GTA CTT -5' | 7784AS | ZF94 | ZF72 | ZF72 | ZF66 | ZF94 | ZF105 | GEKPYKCPECGKSFSSKKHLAEHQRTHTGEKPYKCPECGKSFSTSGELVRHQRTHTGEKPYKCPECGKSFSTSGELVRHQRTHTGEKPYKCPECGKSFSRSDELVRHQRTHTGEKPYKCPECGKSFSSKKHLAEHQRTHTGEKPYKCPECGKSFSQAGHLASHQRTH |
| 29 | 8086S | 5'- AGC AGC CAC GGG TCA GCC | ACAAG | GGC CAC AGC CAT GAA TGG -3' | 8086S | ZF106 | ZF63 | ZF92 | ZF83 | ZF90 | ZF61 | GEKPYKCPECGKSFSRSDHLTTHQRTHTGEKPYKCPECGKSFSQSSNLVRHQRTHTGEKPYKCPECGKSFSTSGNLTEHQRTHTGEKPYKCPECGKSFSERSHLREHQRTHTGEKPYKCPECGKSFSSKKALTEHQRTHTGEKPYKCPECGKSFSDPGHLVRHQRTH |
| | 8087AS | 3'- TCG TCG GTG CCC AGT CGG | TGTTC | CCG GTG TCG GTA CTT ACC -5' | 8087AS | ZF72 | ZF72 | ZF66 | ZF94 | ZF105 | ZF61 | GEKPYKCPECGKSFSTSGELVRHQRTHTGEKPYKCPECGKSFSTSGELVRHQRTHTGEKPYKCPECGKSFSRSDELVRHQRTHTGEKPYKCPECGKSFSSKKHLAEHQRTHTGEKPYKCPECGKSFSQAGHLASHQRTHTGEKPYKCPECGKSFSDPGHLVRHQRTH |
| 30 | 8288S | 5'- CAG CCA CGG GTC AGC CAC | AAGGG | CCA CAG CCA TGA ATG GCA -3' | 8288S | ZF71 | ZF87 | ZF105 | ZF93 | ZF91 | ZF93 | GEKPYKCPECGKSFSQSGDLRRHQRTHTGEKPYKCPECGKSFSRRDELNVHQRTHTGEKPYKCPECGKSFSQAGHLASHQRTHTGEKPYKCPECGKSFSTSHSLTEHQRTHTGEKPYKCPECGKSFSRADNLTEHQRTHTGEKPYKCPECGKSFSTSHSLTEHQRTH |
| | 8289AS | 3'- GTC GGT GCC CAG TCG GTG | TTCCC | GGT GTC GGT ACT TAC CGT -5' | 8289AS | ZF102 | ZF106 | ZF95 | ZF65 | ZF72 | ZF66 | GEKPYKCPECGKSFSRNDALTEHQRTHTGEKPYKCPECGKSFSRSDHLTTHQRTHTGEKPYKCPECGKSFSRNDTLTEHQRTHTGEKPYKCPECGKSFSDPGNLVRHQRTHTGEKPYKCPECGKSFSTSGELVRHQRTHTGEKPYKCPECGKSFSRSDELVRHQRTH |
| 31 | 8591S | 5'- CCA CGG GTC AGC CAC AAG | GGCCA | CAG CCA TGA ATG GCA CAG -3' | 8591S | ZF91 | ZF71 | ZF87 | ZF105 | ZF93 | ZF91 | GEKPYKCPECGKSFSRADNLTEHQRTHTGEKPYKCPECGKSFSQSGDLRRHQRTHTGEKPYKCPECGKSFSRRDELNVHQRTHTGEKPYKCPECGKSFSQAGHLASHQRTHTGEKPYKCPECGKSFSTSHSLTEHQRTHTGEKPYKCPECGKSFSRADNLTEHQRTH |
| | 8592AS | 3'- GGT GCC CAG TCG GTG TTC | CCGGT | GTC GGT ACT TAC CGT GTC -5' | 8592AS | ZF106 | ZF95 | ZF65 | ZF72 | ZF66 | ZF103 | GEKPYKCPECGKSFSRSDHLTTHQRTHTGEKPYKCPECGKSFSRNDTLTEHQRTHTGEKPYKCPECGKSFSDPGNLVRHQRTHTGEKPYKCPECGKSFSTSGELVRHQRTHTGEKPYKCPECGKSFSRSDELVRHQRTHTGEKPYKCPECGKSFSTTGALTEHQRTH |
| 32 | 8892S | 5'- CGG GTC AGC CAC AAG GGC | CACAG | CCA TGA ATG GCA CAG AAG -3' | 8892S | ZF76 | ZF91 | ZF71 | ZF87 | ZF105 | ZF93 | GEKPYKCPECGKSFSRKDNLKNHQRTHTGEKPYKCPECGKSFSRADNLTEHQRTHTGEKPYKCPECGKSFSQSGDLRRHQRTHTGEKPYKCPECGKSFSRRDELNVHQRTHTGEKPYKCPECGKSFSQAGHLASHQRTHTGEKPYKCPECGKSFSTSHSLTEHQRTH |
| | 8895AS | 3'- GCC CAG TCG GTG TTC CCG | GTGTC | GGT ACT TAC CGT GTC TTC -5' | 8895AS | ZF95 | ZF65 | ZF72 | ZF66 | ZF103 | ZF73 | GEKPYKCPECGKSFSRNDTLTEHQRTHTGEKPYKCPECGKSFSDPGNLVRHQRTHTGEKPYKCPECGKSFSTSGELVRHQRTHTGEKPYKCPECGKSFSRSDELVRHQRTHTGEKPYKCPECGKSFSTTGALTEHQRTHTGEKPYKCPECGKSFSDCRDLARHQRTH |
| 33 | 9197S | 5'- GTC AGC CAC AAG GGC CAC | AGCCA | TGA ATG GCA CAG AAG GCC -3' | 9197S | ZF73 | ZF76 | ZF91 | ZF71 | ZF87 | ZF105 | GEKPYKCPECGKSFSDCRDLARHQRTHTGEKPYKCPECGKSFSRKDNLKNHQRTHTGEKPYKCPECGKSFSRADNLTEHQRTHTGEKPYKCPECGKSFSQSGDLRRHQRTHTGEKPYKCPECGKSFSRRDELNVHQRTHTGEKPYKCPECGKSFSQAGHLASHQRTH |
| | 9197AS | 3'- CAG TCG GTG TTC CCG GTG | TCGGT | ACT TAC CGT GTC TTC CGG -5' | 9197AS | ZF65 | ZF72 | ZF66 | ZF103 | ZF73 | ZF66 | GEKPYKCPECGKSFSDPGNLVRHQRTHTGEKPYKCPECGKSFSTSGELVRHQRTHTGEKPYKCPECGKSFSRSDELVRHQRTHTGEKPYKCPECGKSFSTTGALTEHQRTHTGEKPYKCPECGKSFSDCRDLARHQRTHTGEKPYKCPECGKSFSRSDELVRHQRTH |
| 34 | 9410S | 5'- AGC CAC AAG GGC CAC AGC | CATGA | ATG GCA CAG AAG GCC CTA -3' | 9410S | ZF10 | ZF73 | ZF76 | ZF91 | ZF71 | ZF87 | GEKPYKCPECGKSFSQNSTLTEHQRTHTGEKPYKCPECGKSFSDCRDLARHQRTHTGEKPYKCPECGKSFSRKDNLKNHQRTHTGEKPYKCPECGKSFSRADNLTEHQRTHTGEKPYKCPECGKSFSQSGDLRRHQRTHTGEKPYKCPECGKSFSRRDELNVHQRTH |
| | 9410AS | 3'- TCG GTG TTC CCG GTG TCG | GTACT | TAC CGT GTC TTC CGG GAT -5' | 9410AS | ZF72 | ZF66 | ZF10 | ZF73 | ZF66 | ZF72 | GEKPYKCPECGKSFSTSGELVRHQRTHTGEKPYKCPECGKSFSRSDELVRHQRTHTGEKPYKCPECGKSFSTTGALTEHQRTHTGEKPYKCPECGKSFSDCRDLARHQRTHTGEKPYKCPECGKSFSRSDELVRHQRTHTGEKPYKCPECGKSFSTSGELVRHQRTH |
| Compatible ZF | 5562AS-C2 | | | | 5562AS-C2 | ZF73 | ZF76 | ZF70 | ZF72 | ZF77 | ZF52 | GEKPYKCPECGKSFSDCRDLARHQRTHTGEKPYKCPECGKSFSRKDNLKNHQRTHTGEKPYKCPECGKSFSRSDDLVRHQRTHTGEKPYKCPECGKSFSTSGELVRHQRTHTGEKPYKCPECGKSFSTTGNLTVHQRTHTGEKKFACPECPKRFMRSDNLTQHIKTH |
| | 5562AS-C3 | | | | 5562AS-C3 | ZF73 | ZF76 | ZF70 | ZF72 | ZF137 | ZF76 | GEKPYKCPECGKSFSDCRDLARHQRTHTGEKPYKCPECGKSFSRKDNLKNHQRTHTGEKPYKCPECGKSFSRSDDLVRHQRTHTGEKPYKCPECGKSFSTSGELVRHQRTHTGEKPYECDHCGKAFSVSSNLNVHRRIHTGEKPYKCPECGKSFSRKDNLKNHQRTH |
| | 5562AS-C4 | | | | 5562AS-C4 | ZF73 | ZF76 | ZF70 | ZF72 | ZF137 | ZF52 | GEKPYKCPECGKSFSDCRDLARHQRTHTGEKPYKCPECGKSFSRKDNLKNHQRTHTGEKPYKCPECGKSFSRSDDLVRHQRTHTGEKPYKCPECGKSFSTSGELVRHQRTHTGEKPYECDHCGKAFSVSSNLNVHRRIHTGEKKFACPECPKRFMRSDNLTQHIKTH |
| | 5562AS-C5 | 3'- CCG GAA GCG TCG TAA GAA | CCCAC | CCT CGT CGG TGC CCA GTC -5' | 5562AS-C5 | ZF73 | ZF52 | ZF70 | ZF72 | ZF77 | ZF76 | GEKPYKCPECGKSFSDCRDLARHQRTHTGEKKFACPECPKRFMRSDNLTQHIKTHTGEKPYKCPECGKSFSRSDDLVRHQRTHTGEKPYKCPECGKSFSTSGELVRHQRTHTGEKPYKCPECGKSFSTTGNLTVHQRTHTGEKPYKCPECGKSFSRKDNLKNHQRTH |
| | 5562AS-C6 | | | | 5562AS-C6 | ZF73 | ZF52 | ZF70 | ZF72 | ZF77 | ZF52 | GEKPYKCPECGKSFSDCRDLARHQRTHTGEKKFACPECPKRFMRSDNLTQHIKTHTGEKPYKCPECGKSFSRSDDLVRHQRTHTGEKPYKCPECGKSFSTSGELVRHQRTHTGEKPYKCPECGKSFSTTGNLTVHQRTHTGEKKFACPECPKRFMRSDNLTQHIKTH |
| | 5562AS-C7 | | | | 5562AS-C7 | ZF73 | ZF52 | ZF70 | ZF72 | ZF137 | ZF76 | GEKPYKCPECGKSFSDCRDLARHQRTHTGEKKFACPECPKRFMRSDNLTQHIKTHTGEKPYKCPECGKSFSRSDDLVRHQRTHTGEKPYKCPECGKSFSTSGELVRHQRTHTGEKPYECDHCGKAFSVSSNLNVHRRIHTGEKPYKCPECGKSFSRKDNLKNHQRTH |
| | 5562AS-C8 | | | | 5562AS-C8 | ZF73 | ZF52 | ZF70 | ZF72 | ZF137 | ZF52 | GEKPYKCPECGKSFSDCRDLARHQRTHTGEKKFACPECPKRFMRSDNLTQHIKTHTGEKPYKCPECGKSFSRSDDLVRHQRTHTGEKPYKCPECGKSFSTSGELVRHQRTHTGEKPYECDHCGKAFSVSSNLNVHRRIHTGEKKFACPECPKRFMRSDNLTQHIKTH |

ZF module numbers and amino acid sequences follow the Zinc Finger Consortium nomenclature (Addgene Kit #1000000005; https://www.addgene.org/kits/zfc-modular-assembly/). Module selection was performed using ZF Tools.

### Supplemental Table S3 Target sequences of ZF pairs for mouse Rho 5'-UTR

| ZF pairs used in this study | | | | | ZF modules (Wright et al. 2006) | | | | | | | |
|---|---|---|---|---|---|---|---|---|---|---|---|---|
| No | Name | antisense | Spacer | sense | Name | F1 | F2 | F3 | F4 | F5 | F6 | Amino acid sequence (6ZF modules) |
| 1 | 5057S | 5'- TCG CCA AGC AGC CTT GGT | CTCTGT | CTA CGA AGA GCC CGT GGG -3' | 5057S | ZF58 | ZF100 | ZF73 | ZF82 | ZF97 | ZF101 | GEKPYKCPECGKSFSRSDKLVRHQRTHTGEKPYKCPECGKSFSSRRTCRAHQRTHTGEKPYKCPECGKSFSDCRDLARHQRTHTGEKPYKCPECGKSFSQLAHLRAHQRTHTGEKPYKCPECGKSFSQSGHLTEHQRTHTGEKPYKCPECGKSFSQNSTLTEHQRTHT |
| | 5057AS | 3'- AGC GGT TCG TCG GAA CCA | GAGACA | GAT GCT TCT CGG GCA CCC -5' | 5057AS | ZF97 | ZF106 | ZF72 | ZF72 | ZF76 | ZF79 | GEKPYKCPECGKSFSQSGHLTEHQRTHTGEKPYKCPECGKSFSRSDHLTTHQRTHTGEKPYKCPECGKSFSTSGELVRHQRTHTGEKPYKCPECGKSFSTSGELVRHQRTHTGEKPYKCPECGKSFSRKDNLKNHQRTHTGEKPYKCPECGKSFSDKKDLTRHQRTHT |
| 2 | 5259S | 5'- GCC AAG CAG CCT TGG TCT | CTGTCT | ACG AAG AGC CCG TGG GGC -3' | 5259S | ZF61 | ZF106 | ZF95 | ZF83 | ZF76 | ZF80 | GEKPYKCPECGKSFSDPGHLVRHQRTHTGEKPYKCPECGKSFSRSDHLTTHQRTHTGEKPYKCPECGKSFSRNDTLTEHQRTHTGEKPYKCPECGKSFSERSHLREHQRTHTGEKPYKCPECGKSFSRKDNLKNHQRTHTGEKPYKCPECGKSFSRTDTLRDHQRTHT |
| | 5259AS | 3'- CGG TTC GTC GGA ACC AGA | GACAGA | TGC TTC TCG GGC ACC CCG -5' | 5259AS | ZF61 | ZF103 | ZF102 | ZF84 | ZF93 | ZF82 | GEKPYKCPECGKSFSDPGHLVRHQRTHTGEKPYKCPECGKSFSTTGALTEHQRTHTGEKPYKCPECGKSFSRNDALTEHQRTHTGEKPYKCPECGKSFSRSDHLTNHQRTHTGEKPYKCPECGKSFSTSHSLTEHQRTHTGEKPYKCPECGKSFSQLAHLRAHQRTHT |
| 3 | 5360S | 5'- CCA AGC AGC CTT GGT CTC | TGTCTA | CGA AGA GCC CGT GGG GCA -3' | 5360S | ZF71 | ZF58 | ZF100 | ZF73 | ZF82 | ZF97 | GEKPYKCPECGKSFSQSGDLRRHQRTHTGEKPYKCPECGKSFSRSDKLVRHQRTHTGEKPYKCPECGKSFSSRRTCRAHQRTHTGEKPYKCPECGKSFSDCRDLARHQRTHTGEKPYKCPECGKSFSQLAHLRAHQRTHTGEKPYKCPECGKSFSQSGHLTEHQRTHT |
| | 5360AS | 3'- GGT TCG TCG GAA CCA GAG | ACAGAT | GCT TCT CGG GCA CCC CGT -5' | 5360AS | ZF106 | ZF72 | ZF72 | ZF76 | ZF79 | ZF62 | GEKPYKCPECGKSFSRSDHLTTHQRTHTGEKPYKCPECGKSFSTSGELVRHQRTHTGEKPYKCPECGKSFSTSGELVRHQRTHTGEKPYKCPECGKSFSRKDNLKNHQRTHTGEKPYKCPECGKSFSDKKDLTRHQRTHTGEKPYKCPECGKSFSRSDNLVRHQRTHT |
| 4 | 5562S | 5'- AAG CAG CCT TGG TCT CTG | TCTACG | AAG AGC CCG TGG GGC AGC -3' | 5562S | ZF83 | ZF61 | ZF106 | ZF95 | ZF83 | ZF76 | GEKPYKCPECGKSFSERSHLREHQRTHTGEKPYKCPECGKSFSDPGHLVRHQRTHTGEKPYKCPECGKSFSRSDHLTTHQRTHTGEKPYKCPECGKSFSRNDTLTEHQRTHTGEKPYKCPECGKSFSERSHLREHQRTHTGEKPYKCPECGKSFSRKDNLKNHQRTHT |
| | 5562AS | 3'- TTC GTC GGA ACC AGA GAC | AGATGC | TTC TCG GGC ACC CCG TCG -5' | 5562AS | ZF103 | ZF102 | ZF84 | ZF93 | ZF82 | ZF91 | GEKPYKCPECGKSFSTTGALTEHQRTHTGEKPYKCPECGKSFSRNDALTEHQRTHTGEKPYKCPECGKSFSRSDHLTNHQRTHTGEKPYKCPECGKSFSTSHSLTEHQRTHTGEKPYKCPECGKSFSQLAHLRAHQRTHTGEKPYKCPECGKSFSRADNLTEHQRTHT |
| 5 | 5663S | 5'- AGC AGC CTT GGT CTC TGT | CTACGA | AGA GCC CGT GGG GCA GCC -3' | 5663S | ZF73 | ZF71 | ZF58 | ZF100 | ZF73 | ZF82 | GEKPYKCPECGKSFSDCRDLARHQRTHTGEKPYKCPECGKSFSQSGDLRRHQRTHTGEKPYKCPECGKSFSRSDKLVRHQRTHTGEKPYKCPECGKSFSSRRTCRAHQRTHTGEKPYKCPECGKSFSDCRDLARHQRTHTGEKPYKCPECGKSFSQLAHLRAHQRTHT |
| | 5663AS | 3'- TCG TCG GAA CCA GAG ACA | GATGCT | TCT CGG GCA CCC CGT CGG -5' | 5663AS | ZF72 | ZF72 | ZF76 | ZF79 | ZF62 | ZF78 | GEKPYKCPECGKSFSTSGELVRHQRTHTGEKPYKCPECGKSFSTSGELVRHQRTHTGEKPYKCPECGKSFSRKDNLKNHQRTHTGEKPYKCPECGKSFSDKKDLTRHQRTHTGEKPYKCPECGKSFSRSDNLVRHQRTHTGEKPYKCPECGKSFSSPADLTRHQRTHT |
| 6 | 6067S | 5'- GCC TTG GTC TCT GTC TAC | GAAGAG | CCC GTG GGG CAG CCT CGA -3' | 6067S | ZF97 | ZF96 | ZF91 | ZF58 | ZF66 | ZF94 | GEKPYKCPECGKSFSQSGHLTEHQRTHTGEKPYKCPECGKSFSTKNSLTEHQRTHTGEKPYKCPECGKSFSRADNLTEHQRTHTGEKPYKCPECGKSFSRSDKLVRHQRTHTGEKPYKCPECGKSFSRSDELVRHQRTHTGEKPYKCPECGKSFSSKKHLAEHQRTHT |
| | 6067AS | 3'- CGG AAC CAG AGA CAG ATG | CTTCTC | GGG CAC CCC GTC GGA GCT -5' | 6067AS | ZF61 | ZF89 | ZF65 | ZF82 | ZF65 | ZF67 | GEKPYKCPECGKSFSDPGHLVRHQRTHTGEKPYKCPECGKSFSQSGNLTEHQRTHTGEKPYKCPECGKSFSDPGNLVRHQRTHTGEKPYKCPECGKSFSQLAHLRAHQRTHTGEKPYKCPECGKSFSDPGNLVRHQRTHTGEKPYKCPECGKSFSQSSSLVRHQRTHT |
| 7 | 7683S | 5'- ACG AAG AGC CCG TGG GGC | AGCCTC | GAG AGC CGC AGC CAT GAA -3' | 7683S | ZF63 | ZF92 | ZF83 | ZF98 | ZF83 | ZF62 | GEKPYKCPECGKSFSQSSNLVRHQRTHTGEKPYKCPECGKSFSTSGNLTEHQRTHTGEKPYKCPECGKSFSERSHLREHQRTHTGEKPYKCPECGKSFSHTGHLLEHQRTHTGEKPYKCPECGKSFSERSHLREHQRTHTGEKPYKCPECGKSFSRSDNLVRHQRTHT |
| | 7683AS | 3'- TGC TTC TCG GGC ACC CCG | TCGGAG | CTC TCG GCG TCG GTA CTT -5' | 7683AS | ZF100 | ZF103 | ZF72 | ZF99 | ZF93 | ZF73 | GEKPYKCPECGKSFSSRRTCRAHQRTHTGEKPYKCPECGKSFSTTGALTEHQRTHTGEKPYKCPECGKSFSTSGELVRHQRTHTGEKPYKCPECGKSFSRSDKLTEHQRTHTGEKPYKCPECGKSFSTSHSLTEHQRTHTGEKPYKCPECGKSFSDCRDLARHQRTHT |
| 8 | 7986S | 5'- AAG AGC CCG TGG GGC AGC | CTCGAG | AGC CGC AGC CAT GAA CGG -3' | 7986S | ZF99 | ZF63 | ZF92 | ZF83 | ZF98 | ZF83 | GEKPYKCPECGKSFSRSDKLTEHQRTHTGEKPYKCPECGKSFSQSSNLVRHQRTHTGEKPYKCPECGKSFSTSGNLTEHQRTHTGEKPYKCPECGKSFSERSHLREHQRTHTGEKPYKCPECGKSFSHTGHLLEHQRTHTGEKPYKCPECGKSFSERSHLREHQRTHT |
| | 7986AS | 3'- TTC TCG GGC ACC CCG TCG | GAGCTC | TCG GCG TCG GTA CTT GCC -5' | 7986AS | ZF103 | ZF72 | ZF99 | ZF93 | ZF73 | ZF72 | GEKPYKCPECGKSFSTTGALTEHQRTHTGEKPYKCPECGKSFSTSGELVRHQRTHTGEKPYKCPECGKSFSRSDKLTEHQRTHTGEKPYKCPECGKSFSTSHSLTEHQRTHTGEKPYKCPECGKSFSDCRDLARHQRTHTGEKPYKCPECGKSFSTSGELVRHQRTHT |
| 9 | 8289S | 5'- AGC CCG TGG GGC AGC CTC | GAGAGC | CGC AGC CAT GAA CGG CAC -3' | 8289S | ZF90 | ZF99 | ZF63 | ZF92 | ZF83 | ZF98 | GEKPYKCPECGKSFSSKKALTEHQRTHTGEKPYKCPECGKSFSRSDKLTEHQRTHTGEKPYKCPECGKSFSQSSNLVRHQRTHTGEKPYKCPECGKSFSTSGNLTEHQRTHTGEKPYKCPECGKSFSERSHLREHQRTHTGEKPYKCPECGKSFSHTGHLLEHQRTHT |
| | 8289AS | 3'- TCG GGC ACC CCG TCG GAG | CTCTCG | GCG TCG GTA CTT GCC GTG -5' | 8289AS | ZF72 | ZF99 | ZF93 | ZF73 | ZF72 | ZF62 | GEKPYKCPECGKSFSTSGELVRHQRTHTGEKPYKCPECGKSFSRSDKLTEHQRTHTGEKPYKCPECGKSFSTSHSLTEHQRTHTGEKPYKCPECGKSFSDCRDLARHQRTHTGEKPYKCPECGKSFSTSGELVRHQRTHTGEKPYKCPECGKSFSRSDNLVRHQRTHT |
| 10 | 5259S | 5'- CCA AGC AGC CTT GGT CTC | TGTCT | ACG AAG AGC CCG TGG GGC -3' | 5259S | ZF61 | ZF106 | ZF95 | ZF83 | ZF76 | ZF80 | GEKPYKCPECGKSFSDPGHLVRHQRTHTGEKPYKCPECGKSFSRSDHLTTHQRTHTGEKPYKCPECGKSFSRNDTLTEHQRTHTGEKPYKCPECGKSFSERSHLREHQRTHTGEKPYKCPECGKSFSRKDNLKNHQRTHTGEKPYKCPECGKSFSRTDTLRDHQRTHT |
| | 5360AS | 3'- GGT TCG TCG GAA CCA GAG | ACAGA | TGC TTC TCG GGC ACC CCG -5' | 5359AS | ZF106 | ZF72 | ZF72 | ZF76 | ZF79 | ZF62 | GEKPYKCPECGKSFSRSDHLTTHQRTHTGEKPYKCPECGKSFSTSGELVRHQRTHTGEKPYKCPECGKSFSTSGELVRHQRTHTGEKPYKCPECGKSFSRKDNLKNHQRTHTGEKPYKCPECGKSFSDKKDLTRHQRTHTGEKPYKCPECGKSFSRSDNLVRHQRTHT |
| 11 | 5561S | 5'- AAG CAG CCT TGG TCT CTG | TCTAC | GAA GAG CCC GTG GGG CAG -3' | 5561S | ZF91 | ZF58 | ZF66 | ZF94 | ZF62 | ZF63 | GEKPYKCPECGKSFSRADNLTEHQRTHTGEKPYKCPECGKSFSRSDKLVRHQRTHTGEKPYKCPECGKSFSRSDELVRHQRTHTGEKPYKCPECGKSFSSKKHLAEHQRTHTGEKPYKCPECGKSFSRSDNLVRHQRTHTGEKPYKCPECGKSFSQSSNLVRHQRTHT |
| | 5562AS | 3'- TTC GTC GGA ACC AGA GAC | AGATG | CTT CTC GGG CAC CCC GTC -5' | 5562AS | ZF103 | ZF102 | ZF84 | ZF93 | ZF82 | ZF91 | GEKPYKCPECGKSFSTTGALTEHQRTHTGEKPYKCPECGKSFSRNDALTEHQRTHTGEKPYKCPECGKSFSRSDHLTNHQRTHTGEKPYKCPECGKSFSTSHSLTEHQRTHTGEKPYKCPECGKSFSQLAHLRAHQRTHTGEKPYKCPECGKSFSRADNLTEHQRTHT |
| 12 | 5562S | 5'- AGC AGC CTT GGT CTC TGT | CTACG | AAG AGC CCG TGG GGC AGC -3' | 5562S | ZF83 | ZF61 | ZF106 | ZF95 | ZF83 | ZF76 | GEKPYKCPECGKSFSERSHLREHQRTHTGEKPYKCPECGKSFSDPGHLVRHQRTHTGEKPYKCPECGKSFSRSDHLTTHQRTHTGEKPYKCPECGKSFSRNDTLTEHQRTHTGEKPYKCPECGKSFSERSHLREHQRTHTGEKPYKCPECGKSFSRKDNLKNHQRTHT |
| | 5663AS | 3'- TCG TCG GAA CCA GAG ACA | GATGC | TTC TCG GGC ACC CCG TCG -5' | 5663AS | ZF72 | ZF72 | ZF76 | ZF79 | ZF62 | ZF78 | GEKPYKCPECGKSFSTSGELVRHQRTHTGEKPYKCPECGKSFSTSGELVRHQRTHTGEKPYKCPECGKSFSRKDNLKNHQRTHTGEKPYKCPECGKSFSDKKDLTRHQRTHTGEKPYKCPECGKSFSRSDNLVRHQRTHTGEKPYKCPECGKSFSSPADLTRHQRTHT |
| 13 | 5864S | 5'- CAG CCT TGG TCT CTG TCT | ACGAA | GAG CCC GTG GGG CAG CCT -3' | 5864S | ZF96 | ZF91 | ZF58 | ZF66 | ZF94 | ZF62 | GEKPYKCPECGKSFSTKNSLTEHQRTHTGEKPYKCPECGKSFSRADNLTEHQRTHTGEKPYKCPECGKSFSRSDKLVRHQRTHTGEKPYKCPECGKSFSRSDELVRHQRTHTGEKPYKCPECGKSFSSKKHLAEHQRTHTGEKPYKCPECGKSFSRSDNLVRHQRTHT |
| | 5864AS | 3'- GTC GGA ACC AGA GAC AGA | TGCTT | CTC GGG CAC CCC GTC GGA -5' | 5864AS | ZF102 | ZF84 | ZF93 | ZF82 | ZF91 | ZF82 | GEKPYKCPECGKSFSRNDALTEHQRTHTGEKPYKCPECGKSFSRSDHLTNHQRTHTGEKPYKCPECGKSFSTSHSLTEHQRTHTGEKPYKCPECGKSFSQLAHLRAHQRTHTGEKPYKCPECGKSFSRADNLTEHQRTHTGEKPYKCPECGKSFSQLAHLRAHQRTHT |
| 14 | 6067S | 5'- CCT TGG TCT CTG TCT ACG | AAGAG | CCC GTG GGG CAG CCT CGA -3' | 6067S | ZF97 | ZF96 | ZF91 | ZF58 | ZF66 | ZF94 | GEKPYKCPECGKSFSQSGHLTEHQRTHTGEKPYKCPECGKSFSTKNSLTEHQRTHTGEKPYKCPECGKSFSRADNLTEHQRTHTGEKPYKCPECGKSFSRSDKLVRHQRTHTGEKPYKCPECGKSFSRSDELVRHQRTHTGEKPYKCPECGKSFSSKKHLAEHQRTHT |
| | 6167AS | 3'- GGA ACC AGA GAC AGA TGC | TTCTC | GGG CAC CCC GTC GGA GCT -5' | 6167AS | ZF84 | ZF93 | ZF82 | ZF91 | ZF82 | ZF100 | GEKPYKCPECGKSFSRSDHLTNHQRTHTGEKPYKCPECGKSFSTSHSLTEHQRTHTGEKPYKCPECGKSFSQLAHLRAHQRTHTGEKPYKCPECGKSFSRADNLTEHQRTHTGEKPYKCPECGKSFSQLAHLRAHQRTHTGEKPYKCPECGKSFSSRRTCRAHQRTHT |
| 15 | 6470S | 5'- TGG TCT CTG TCT ACG AAG | AGCCC | GTG GGG CAG CCT CGA GAG -3' | 6470S | ZF62 | ZF97 | ZF96 | ZF91 | ZF58 | ZF66 | GEKPYKCPECGKSFSRSDNLVRHQRTHTGEKPYKCPECGKSFSQSGHLTEHQRTHTGEKPYKCPECGKSFSTKNSLTEHQRTHTGEKPYKCPECGKSFSRADNLTEHQRTHTGEKPYKCPECGKSFSRSDKLVRHQRTHTGEKPYKCPECGKSFSRSDELVRHQRTHT |
| | 6470AS | 3'- ACC AGA GAC AGA TGC TTC | TCGGG | CAC CCC GTC GGA GCT CTC -5' | 6470AS | ZF93 | ZF82 | ZF91 | ZF82 | ZF100 | ZF103 | GEKPYKCPECGKSFSTSHSLTEHQRTHTGEKPYKCPECGKSFSQLAHLRAHQRTHTGEKPYKCPECGKSFSRADNLTEHQRTHTGEKPYKCPECGKSFSQLAHLRAHQRTHTGEKPYKCPECGKSFSSRRTCRAHQRTHTGEKPYKCPECGKSFSTTGALTEHQRTHT |
| 16 | 6773S | 5'- TCT CTG TCT ACG AAG AGC | CCGTG | GGG CAG CCT CGA GAG CCG -3' | 6773S | ZF95 | ZF62 | ZF97 | ZF96 | ZF91 | ZF58 | GEKPYKCPECGKSFSRNDTLTEHQRTHTGEKPYKCPECGKSFSRSDNLVRHQRTHTGEKPYKCPECGKSFSQSGHLTEHQRTHTGEKPYKCPECGKSFSTKNSLTEHQRTHTGEKPYKCPECGKSFSRADNLTEHQRTHTGEKPYKCPECGKSFSRSDKLVRHQRTHT |
| | 6773AS | 3'- AGA GAC AGA TGC TTC TCG | GGCAC | CCC GTC GGA GCT CTC GGC -5' | 6773AS | ZF82 | ZF91 | ZF82 | ZF100 | ZF103 | ZF72 | GEKPYKCPECGKSFSQLAHLRAHQRTHTGEKPYKCPECGKSFSRADNLTEHQRTHTGEKPYKCPECGKSFSQLAHLRAHQRTHTGEKPYKCPECGKSFSSRRTCRAHQRTHTGEKPYKCPECGKSFSTTGALTEHQRTHTGEKPYKCPECGKSFSTSGELVRHQRTHT |
| 17 | 7076S | 5'- CTG TCT ACG AAG AGC CCG | TGGGG | CAG CCT CGA GAG CCG CAG -3' | 7076S | ZF91 | ZF95 | ZF62 | ZF97 | ZF96 | ZF91 | GEKPYKCPECGKSFSRADNLTEHQRTHTGEKPYKCPECGKSFSRNDTLTEHQRTHTGEKPYKCPECGKSFSRSDNLVRHQRTHTGEKPYKCPECGKSFSQSGHLTEHQRTHTGEKPYKCPECGKSFSTKNSLTEHQRTHTGEKPYKCPECGKSFSRADNLTEHQRTHT |
| | 7076AS | 3'- GAC AGA TGC TTC TCG GGC | ACCCC | GTC GGA GCT CTC GGC GTC -5' | 7076AS | ZF91 | ZF82 | ZF100 | ZF103 | ZF72 | ZF99 | GEKPYKCPECGKSFSRADNLTEHQRTHTGEKPYKCPECGKSFSQLAHLRAHQRTHTGEKPYKCPECGKSFSSRRTCRAHQRTHTGEKPYKCPECGKSFSTTGALTEHQRTHTGEKPYKCPECGKSFSTSGELVRHQRTHTGEKPYKCPECGKSFSRSDKLTEHQRTHT |
| 18 | 7379S | 5'- TCT ACG AAG AGC CCG TGG | GGCAG | CCT CGA GAG CCG CAG CCA -3' | 7379S | ZF93 | ZF91 | ZF95 | ZF62 | ZF97 | ZF96 | GEKPYKCPECGKSFSTSHSLTEHQRTHTGEKPYKCPECGKSFSRADNLTEHQRTHTGEKPYKCPECGKSFSRNDTLTEHQRTHTGEKPYKCPECGKSFSRSDNLVRHQRTHTGEKPYKCPECGKSFSQSGHLTEHQRTHTGEKPYKCPECGKSFSTKNSLTEHQRTHT |
| | 7379AS | 3'- AGA TGC TTC TCG GGC ACC | CCGTC | GGA GCT CTC GGC GTC GGT -5' | 7379AS | ZF82 | ZF100 | ZF103 | ZF72 | ZF99 | ZF93 | GEKPYKCPECGKSFSQLAHLRAHQRTHTGEKPYKCPECGKSFSSRRTCRAHQRTHTGEKPYKCPECGKSFSTTGALTEHQRTHTGEKPYKCPECGKSFSTSGELVRHQRTHTGEKPYKCPECGKSFSRSDKLTEHQRTHTGEKPYKCPECGKSFSTSHSLTEHQRTHT |
| 19 | 7682S | 5'- ACG AAG AGC CCG TGG GGC | AGCCT | CGA GAG CCG CAG CCA TGA -3' | 7682S | ZF105 | ZF93 | ZF91 | ZF95 | ZF62 | ZF97 | GEKPYKCPECGKSFSQAGHLASHQRTHTGEKPYKCPECGKSFSTSHSLTEHQRTHTGEKPYKCPECGKSFSRADNLTEHQRTHTGEKPYKCPECGKSFSRNDTLTEHQRTHTGEKPYKCPECGKSFSRSDNLVRHQRTHTGEKPYKCPECGKSFSQSGHLTEHQRTHT |
| | 7683AS | 3'- TGC TTC TCG GGC ACC CCG | TCGGA | GCT CTC GGC GTC GGT ACT -5' | 7683AS | ZF100 | ZF103 | ZF72 | ZF99 | ZF93 | ZF73 | GEKPYKCPECGKSFSSRRTCRAHQRTHTGEKPYKCPECGKSFSTTGALTEHQRTHTGEKPYKCPECGKSFSTSGELVRHQRTHTGEKPYKCPECGKSFSRSDKLTEHQRTHTGEKPYKCPECGKSFSTSHSLTEHQRTHTGEKPYKCPECGKSFSDCRDLARHQRTHT |
| 20 | 7985S | 5'- AAG AGC CCG TGG GGC AGC | CTCGA | GAG CCG CAG CCA TGA ACG -3' | 7985S | ZF80 | ZF105 | ZF93 | ZF91 | ZF95 | ZF62 | GEKPYKCPECGKSFSRTDTLRDHQRTHTGEKPYKCPECGKSFSQAGHLASHQRTHTGEKPYKCPECGKSFSTSHSLTEHQRTHTGEKPYKCPECGKSFSRADNLTEHQRTHTGEKPYKCPECGKSFSRNDTLTEHQRTHTGEKPYKCPECGKSFSRSDNLVRHQRTHT |
| | 7986AS | 3'- TTC TCG GGC ACC CCG TCG | GAGCT | CTC GGC GTC GGT ACT TGC -5' | 7986AS | ZF103 | ZF72 | ZF99 | ZF93 | ZF73 | ZF72 | GEKPYKCPECGKSFSTTGALTEHQRTHTGEKPYKCPECGKSFSTSGELVRHQRTHTGEKPYKCPECGKSFSRSDKLTEHQRTHTGEKPYKCPECGKSFSTSHSLTEHQRTHTGEKPYKCPECGKSFSDCRDLARHQRTHTGEKPYKCPECGKSFSTSGELVRHQRTHT |
| 21 | 8288S | 5'- AGC CCG TGG GGC AGC CTC | GAGAG | CCG CAG CCA TGA ACG GCA -3' | 8288S | ZF71 | ZF80 | ZF105 | ZF93 | ZF91 | ZF95 | GEKPYKCPECGKSFSQSGDLRRHQRTHTGEKPYKCPECGKSFSRTDTLRDHQRTHTGEKPYKCPECGKSFSQAGHLASHQRTHTGEKPYKCPECGKSFSTSHSLTEHQRTHTGEKPYKCPECGKSFSRADNLTEHQRTHTGEKPYKCPECGKSFSRNDTLTEHQRTHT |
| | 8289AS | 3'- TCG GGC ACC CCG TCG GAG | CTCTC | GGC GTC GGT ACT TGC CGT -5' | 8289AS | ZF72 | ZF99 | ZF93 | ZF73 | ZF72 | ZF62 | GEKPYKCPECGKSFSTSGELVRHQRTHTGEKPYKCPECGKSFSRSDKLTEHQRTHTGEKPYKCPECGKSFSTSHSLTEHQRTHTGEKPYKCPECGKSFSDCRDLARHQRTHTGEKPYKCPECGKSFSTSGELVRHQRTHTGEKPYKCPECGKSFSRSDNLVRHQRTHT |

ZF module numbers and amino acid sequences follow the Zinc Finger Consortium nomenclature (Addgene Kit #1000000005; https://www.addgene.org/kits/zfc-modular-assembly/). Module selection was performed using ZF Tools.
